# Vitamin D inhibits osteosarcoma by reprogramming nonsense-mediated RNA decay and SNAI2-mediated epithelial-to-mesenchymal transition

**DOI:** 10.1101/2023.01.04.522778

**Authors:** Enrico Capobianco, Vanessa McGaughey, Gerbenn Seraphin, John Heckel, Sandra Rieger, Thomas S. Lisse

**Affiliations:** University of Miami, Department of Biology, Coral Gables, Florida USA; The Jackson Laboratory, Bar Harbor Maine USA; Sylvester Comprehensive Cancer Center, Miller School of Medicine, University of Miami, Miami, Florida USA

**Keywords:** Osteosarcoma, cancer, tumor, vitamin D, vitamin D deficiency, vitamin D receptor, migration, VDR, ROS, MG-63, LM7, EMT, epithelial to mesenchymal transition, osteoblast, SNAI2, snail, metastasis, desmoplasia, microenvironment, carcinoid, oxidative stress, stem cell

## Abstract

Osteosarcomas are immune-resistant and metastatic as a result of elevated nonsense-mediated RNA decay (NMD), reactive oxygen species (ROS), and epithelial-to-mesenchymal transition (EMT). Although vitamin D has anti-cancer effects, its effectiveness and mechanism of action against osteosarcomas are poorly understood. In this study, we assessed the impact of vitamin D and its receptor (VDR) on the NMD-ROS-EMT signaling axis in *in vitro* and *in vivo* osteosarcoma animal models. Initiation of VDR signaling facilitated the enrichment of EMT pathway genes, after which 1,25(OH)_2_D, the active vitamin D derivative, inhibited the EMT pathway in osteosarcoma subtypes. The ligand-bound VDR directly downregulated the EMT inducer *SNAI2*, differentiating highly metastatic from low metastatic subtypes and 1,25(OH)_2_D sensitivity. Moreover, epigenome-wide motif and putative target gene analysis revealed the VDR’s integration with NMD tumorigenic and immunogenic pathways. In an autoregulatory manner, 1,25(OH)_2_D inhibited NMD machinery genes and upregulated NMD target genes implicated in anti-oncogenic activity, immunorecognition, and cell-to-cell adhesion. Dicer substrate siRNA knockdown of *SNAI2* revealed superoxide dismutase 2 (SOD2)-mediated antioxidative responses and 1,25(OH)_2_D sensitization via non-canonical SOD2 nuclear-to-mitochondrial translocalization leading to overall ROS suppression. In a mouse xenograft metastasis model, the therapeutically relevant vitamin D derivative calcipotriol inhibited osteosarcoma metastasis and tumor growth shown for the first time. Our results uncover novel osteosarcoma-inhibiting mechanisms for vitamin D and calcipotriol that may be translated to human patients.

## Introduction

Osteosarcoma (OS) is the most common primary malignant bone tumor in children and adolescents, affecting >1,000 patients annually in the United States alone^1–4^. Osteosarcomas are characterized by a loss of mineralization^5^; moreover, the majority of OS patients have subclinical micrometastases, whereas 30-40% have full-blown metastatic disease^6, 7^. Currently, the standard OS treatments consist of surgical removal of the tumor followed by adjuvant chemotherapy, which causes severe side effects and chemoresistance in patients^3, 4, 8–11^. Several clinical trials evaluating nonconventional OS therapies that combine immunotherapy and cytostatic drug treatments are currently underway in an effort to address this problem^11^. However, despite current treatment options, 30 to 40% of OS patients still die within 5 years of diagnosis^10^. The prognosis for osteosarcoma patients depends on whether it has metastasized at the time of diagnosis, which involves induction of the epithelial-to-mesenchymal transition (EMT), a process that is not only activated in cancer but also in disease-induced fibrosis^12^, and non-disease processes like wound re-epithelialization and embryonic mesoderm formation^13^.

Immunotherapy has thus far been ineffective against OS due to its ability to evade the immune system^14–17^. It is unknown how resistance develops, but it may result from the complex immunological and/or epigenetic reprogramming of OS that creates a tumor permissive environment to support malignancy^3, 4, 10^. In addition, recent studies have shown that the OS microenvironment contains activated cancer associated fibroblasts (CAFs) that can influence desmoplasia, a process in which excessive fibrosis leads to chemoresistance and metastasis of malignant cancer cells, and inhibition of immune cell infiltration that increases tumor burden^18^. Furthermore, genome-wide association studies have linked genetic variations within a neurotransmitter receptor gene to increased risk of OS development, suggesting neuronal-tumor interactions in the manifestation of the disease^19^. Evidence also suggests that nonsense-mediated RNA decay (NMD) is uniquely increased in OS, which, for example, decreases the number of NMD-target neoantigens on OS tumor cells that are required to elicit the cytotoxic killing of cancer cells by immune cells^14–17^. NMD is part of the post-transcriptional RNA surveillance pathway to clear aberrant transcripts harboring premature termination codons (PTCs) often generated by abnormal or ineffective biogenesis of mRNAs or by somatic mutations in cancers^15, 20–23^. To date, the selective targeting and modulation of the NMD pathway to overcome OS resistance for therapeutic considerations has not been reported.

Anti-cancer properties of vitamin D have been studied in major cancer types, however our knowledge of vitamin D’s impact on OS is still very limited. Vitamin D deficiency is a global health issue, with the lowest levels linked to more advanced cancers^24–32^. Clinical randomized studies have demonstrated that vitamin D supplementation reduces the risk of invasive cancer and/or mortality in subgroups of people^25, 26, 33–38^, suggesting that increased circulating vitamin D plays a protective role. Vitamin D is composed of two major forms, vitamin D_2_ (ergocalciferol) and vitamin D_3_ (cholecalciferol)^30^. In humans, vitamin D_3_ is synthesized in the skin in response to UVB exposure, while vitamin D_2_ is obtained from plant sources in our diets with lower efficacy. Both forms of vitamin D are biologically inert and must be converted and hydroxylated to 25(OH)D by vitamin D-25-hydroxylase in the liver, which is the primary measurement of vitamin D status^39^. 25(OH)D is further hydroxylated in the kidneys or within specialized cell types by 25(OH)D-1-OHase (CYP27B1) to produce 1-alpha, 25-dihydroxyvitamin D (also known as 1,25(OH)_2_D), the biologically active derivative of vitamin D^30, 40–44^. The 1,25(OH)_2_D effects are mediated by the vitamin D receptor (VDR), an intracellular nuclear receptor superfamily member, that can promote cell cycle arrest and apoptosis via post-transcriptional and gene regulatory means^40, 45–55^. In mammals, the physiological and molecular effects of vitamin D_3_ on bone health and function have been experimentally and clinically validated^30^. Vitamin D_3_ is known to directly promote the differentiation of normal bone-forming osteoblast-lineage cells and primary human and murine osteoblast cells *in vivo*^56^ and *in vitro*^48, 51^, respectively. Although numerous studies have implicated a suppressive role for vitamin D_3_ in invasive cancer development (e.g., breast, prostate) and improved cancer patient and animal survival^1–4, 26–30^, the functional role of vitamin D_3_ and the VDR in OS is unclear. Uncertain is whether vitamin D_3_ can be utilized to address a number of crucial concerns about OS biology and therapy, such as the regulation of transformation, cancer fibrosis, NMD, and metastasis. Using both *in vitro* and *in vivo* systems, this work examined the influence of the VDR and vitamin D_3_ on the regulation and pathogenesis of OS through impact on OS growth, migration and potential cancer cell immunorecognition.

## Results

### 1,25(OH)_2_D promotes the enrichment of the EMT pathway in MG63 osteosarcoma cells

Gene set enrichment analysis (GSEA)^57^ was used to identify novel pathways using RNAseq data from MG63 OS cells treated with 1,25(OH)_2_D (GEO accession: GSE220948). Using the annotated gene set file M5930 from the Molecular Signatures Database (MSigDB), the EMT gene set was identified as being significantly enriched (**Fig. 1A**)^58^. Using a physiologically-relevant concentration range (10-100nM)^59^, 1,25(OH)_2_D inhibited the expression of EMT inducers and regulators such as *SNAI2* (also known as the transrepressor *SLUG*), *CD44*, and *MMP3*, as confirmed by quantitative PCR (qPCR) (**Fig. 1B**). Higher 1,25(OH)_2_D concentrations increased *MMP3*, both of which are known to induce apoptosis^(47, 48, 60)^. In addition, 1,25(OH)_2_D downregulated and upregulated, respectively, *VIM* and *CLDN1*, which were not annotated in M593, indicating that vitamin D_3_ may inhibit EMT via downregulation of intermediate filaments and the promotion of cell-to-cell adhesion (**Fig. 1C**)^61, 62^. ATACseq was used to assess 1,25(OH)_2_D-dependent genome-wide chromatin accessibility of genes (**Fig. 1D**), and potential promoter regulatory regions were identified within *CD44* and *SNAI2*, but not *MMP3*, *VIM*, and *CLDN1* (**Fig. 1E & S1A**). Intriguingly, 1,25(OH)_2_D did not regulate the major epithelial cadherin, *CDH1*, which is known to be repressed by SNAI2^63^, indicating that other essential cell-cell mediators (such as claudins) are involved in OS (**Fig. 1E**). Moreover, after 1,25(OH)_2_D treatment, the protein expression of SNAI2, MMP3, and CD44 all decreased (**Fig. 1F-H & S1B**). Surprisingly, 1,25(OH)_2_D treatment decreased CD44 nuclear localization, indicating that 1,25(OH)_2_D may inhibit the proteolytic processing of CD44’s intracellular cytoplasmic carboxy-terminal domain in order to limit tumor progression and metastasis via alternative genomic interactions (**Fig. 1I**)^64^.

**Figure 1.**
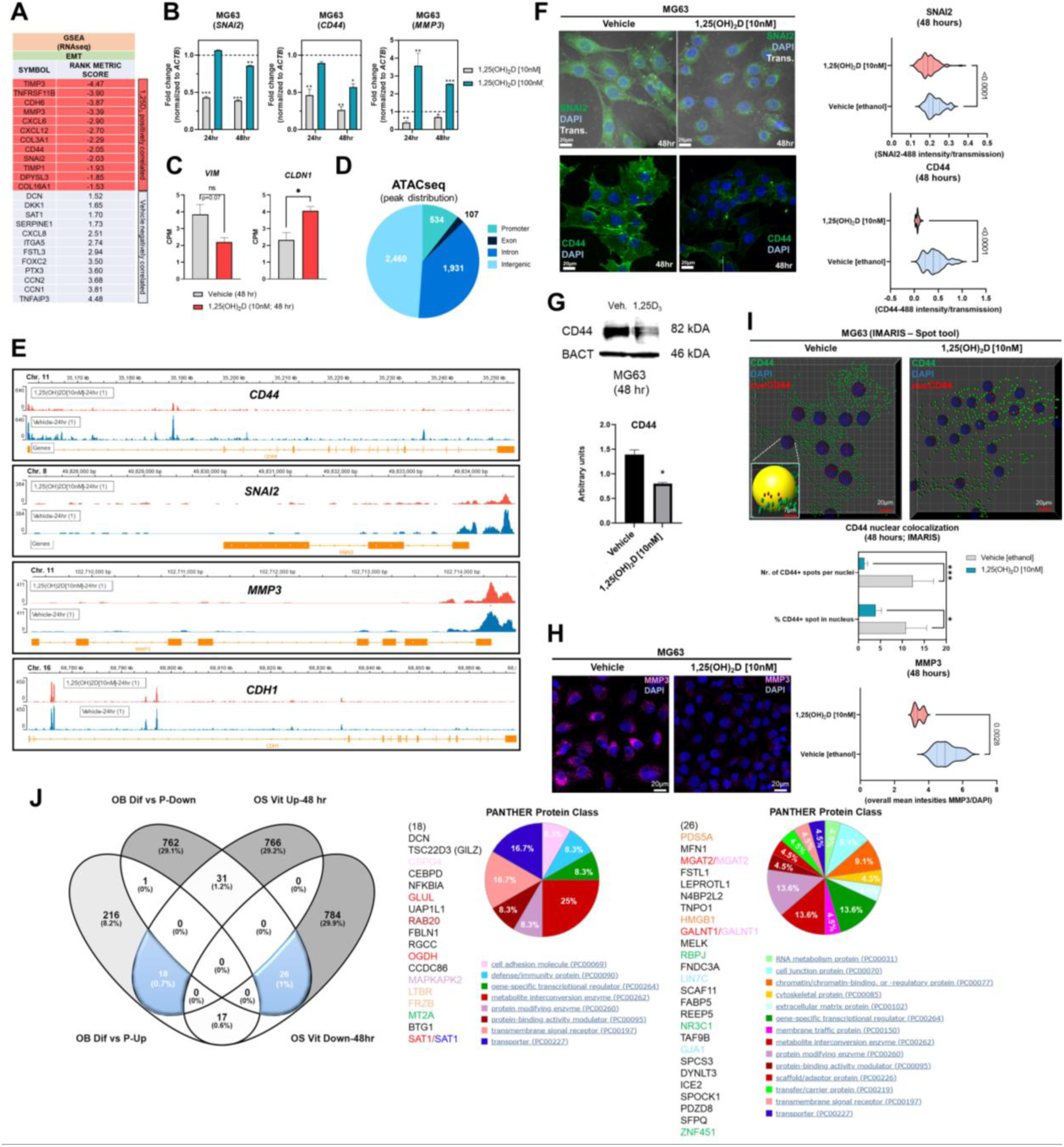
1,25(OH)_2_D treatment of MG63 cells results in enrichment of EMT pathway genes. A) Gene set enrichment analysis (GSEA) was performed to identify a collection of *a priori* EMT-related genes that were over-represented in our RNAseq experiment and may be associated with 1,25(OH)_2_D anti-cancer effects. The MSigDB database was utilized for GSEA at http://www.broadinstitute.org/gsea. B) Quantitative PCR (qPCR) study of genes associated with EMT. Tukey’s multiple comparison test, two-way ANOVA; *p* ≤ ****0.0001, ***0.001, **0.01, *0.05 (n=3). C) MG63 RNA-seq study of EMT genes that are not contained in the MSigDB database. Test of independence (T); *p* ≤ *0.05 (n=1) D) Top: ATAC-seq analysis and peak distribution of MG63 cells treated for 24 hours with 1,25(OH)_2_D. Bottom: ATAC-seq heatmap of 1,25(OH)_2_D regulated genes about the transcription start sites (TSS) E) Visualization of ATAC-seq tracks utilizing the IGV genome browser in order to identify accessible chromatin areas and read peaks. Data represents 24 hours of treatment with 10nM 1,25(OH)_2_D in MG63 cells. All scales indicate the minimum and maximum reads in that window using IGV auto-scale. F) Left: Immunofluorescence study of SNAI2 and CD44 in MG63 cells 48 hours after treatment with 1,25(OH)_2_D. Bars = 20µm. Right: Quantitative study of SNAI2 and CD44-coupled fluorescence intensities in MG63 samples treated with 1,25(OH)_2_D versus vehicle. Individual cell areas and transmitted light were used to equalize intensities. Between the vehicle and treatment data sets, an unpaired t test was conducted, with two-tailed *p*-value summaries displayed in the graphs. The violin plots (n=4) depict the quartiles and medians (darker dashes). G) Western blot study of CD44 utilizing MG63 cells treated for 24 hours with 10nM 1,25(OH)_2_D. To facilitate comparisons, LiCOR band intensities were converted to arbitrary units. Test of independence; *p* ≤ *0.05 (n=3). H) Left: Immunofluorescence confocal imaging study of MG63 cells reveals a decrease in MMP3 expression 48 hours after treatment with 10nM 1,25(OH)_2_D. Right: The violin plots reflect n=4 experimental conditions with an average of 20-40 cells (Unpaired T test). I) An IMARIS colocalization analysis of MG63 cells treated for 48 hours with 10nM 1,25(OH)_2_D. Compared to 1,25(OH)_2_D-treated controls, CD44 nuclear localized expression was more prevalent in untreated controls. Two-way ANOVA Multiple comparisons using Tukey’s test; *p* ≤ ***0.001, *0.05 (n=3). J) Left: A comparative transcriptome study of differentiated (Dif) and proliferating (P) normal osteoblasts in comparison to 1,25(OH)_2_D-treated MG63 OS cells. Normal osteoblast gene set generated from GSE39262. Set expression value (A) to 4 and cut-off to 2-fold change. Right: Panther analysis of subsets of up-and down-regulated genes

A comparison to normal osteoblast states (differentiation *versus* proliferation) was conducted to determine the relative biological effects of 1,25(OH)_2_D on OS. We compared the normal osteoblast transcriptomic data from GSE39262 with our RNAseq data from MG63 OS cells (**Fig. 1J**). Compared to MG63 genes that were either upregulated or downregulated in response to 1,25(OH)_2_D, there was little overlap (0.7% and 1%, respectively) between the genes that define differentiated normal osteoblasts (**Fig. 1J left**). Except for decorin (*DCN*), the remaining genes regulated by 1,25(OH)_2_D did not overlap with either normal proliferation or differentiation genes, indicating that one of vitamin D_3_’s function toward OS may involve other processes such as EMT regulation (**Fig. 1J middle**). 1,25(OH)_2_D upregulated the cell adhesion factor *CSPG4* and the WNT antagonist *FRZB*, but a greater proportion of genes (>29%) were excluded from differentiation or proliferation pathways. Similarly, several cell proliferation/DNA replication cancer-inducing genes, such as *PDS5A*, *HMGB1*, *FABP5*, and *TAF9B*, were downregulated in differentiated osteoblasts, but they represented less than 1% of the downregulated 1,25(OH)_2_D-dependent genes in MG63 cells, indicating that the majority of vitamin D’s effects on OS may involve functional regulation of processes such as EMT (>29%) instead (**Fig. 1J right**).

### 1,25(OH)_2_D inhibits wound induced EMT and migration of osteosarcoma cells

Despite the findings of the GSEA, the biological effects of 1,25(OH)_2_D on EMT was unclear. To determine the effect of 1,25(OH)_2_D on cell migration, we performed an *in vitro* scratch assay on OS cell models MG63 and LM7, which are low and highly metastatic, respectively^65^. After 72 hours of 10nM 1,25(OH)_2_D treatment, MG63 cell migration from the leading edge was reduced by approximately 50% compared to vehicle (**Fig. 2A-C**). In contrast, after 72 hours of treatment with 10nM 1,25(OH)_2_D, scratch migration of LM7 cells was inhibited by 26% (**Fig. 2D,E**), indicating relative resistance. The Boyden Chamber assay, which quantifies the number of invading cells atop a cell-permeable polyester porous membrane over a larger well containing osteogenic media, was also used to examine the response to 1,25(OH)_2_D. MG63 cells remained on the surface of the membrane regardless of treatment conditions, confirming their low metastatic potential (**Fig. 2F**). In contrast, metastatic LM7 cells invaded the membrane after 24 hours, with a statistically significant reduction in the number of invading cells after 1,25(OH)_2_D treatment compared to vehicle control (**Fig. 2G**). The results indicate that 1,25(OH)_2_D inhibits EMT in both OS cell models to varying degrees.

**Figure 2.**
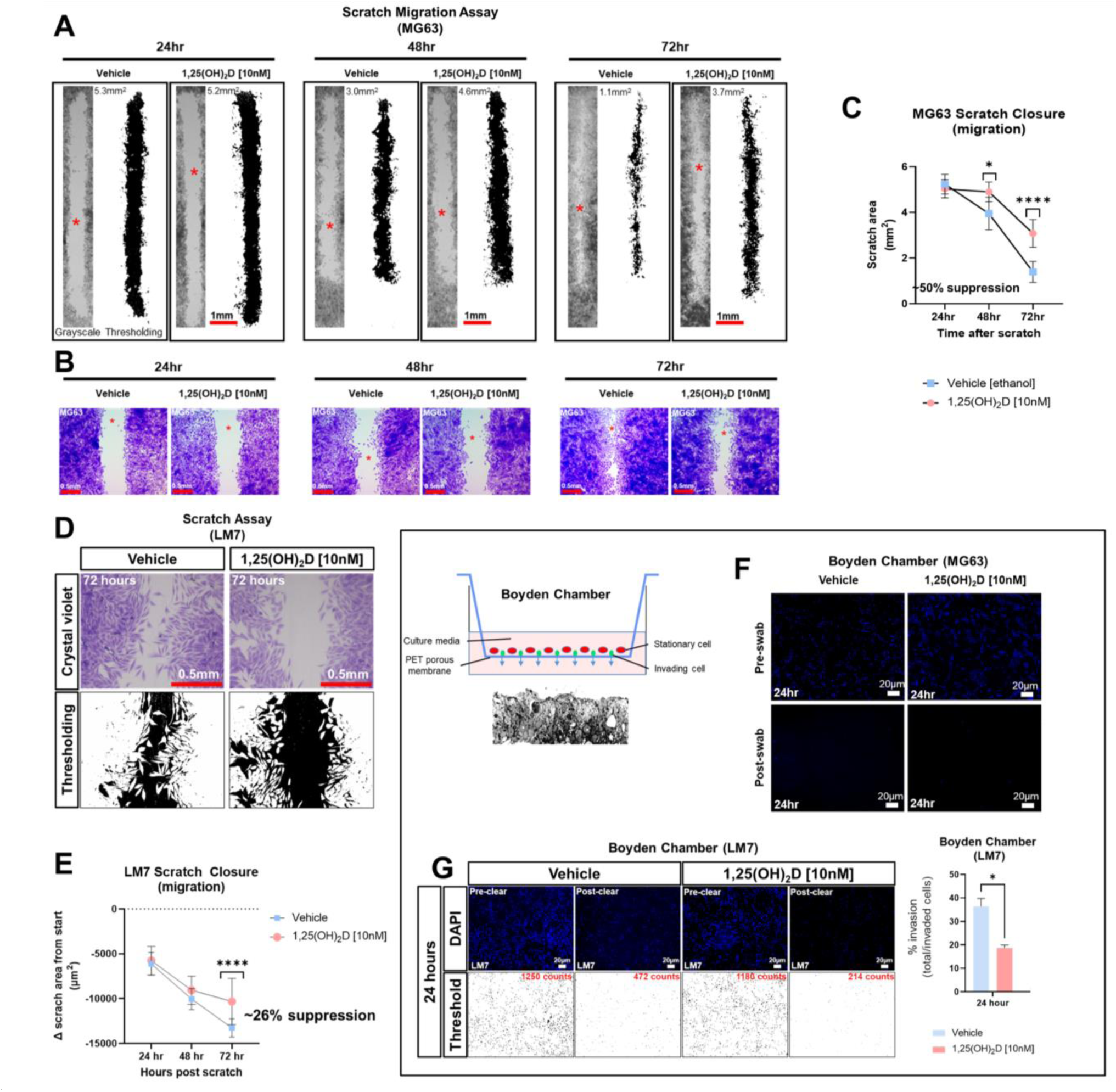
1,25(OH)_2_D inhibits osteosarcoma cell migration and invasion. A) MG63 cells were subjected to scratch assays for 24-72 hours in the presence of vehicle or 10nM 1,25(OH)_2_D. ImageJ thresholding was employed to compute surface areas. The red asterisks indicate the region of increased magnification and crystal violet staining depicted in B. B) Staining with crystal violet of migratory MG63 cells. C) Analysis of the MG63 scratch assay. The graph illustrates n=4 experimental setups for each condition, along with the average scratch area. Two-way ANOVA Multiple comparisons using Tukey’s test; *p* ≤ *0.05 and ****0.0001. D) Metastatic LM7 cells were subjected to scratch studies for 24-72 hours with vehicle or 10nM 1,25(OH)_2_D. E) Analysis of the LM7 scratch assay. The graph illustrates n=4 experimental setups for each condition, with the average scratch area indicated. Two-way ANOVA Tukey’s multiple comparison test; *p* ≤ ****0.0001. Below is a sectional view of the membrane. G) Boyden chamber assay using metastatic LM7 cells after 24 hours. Thresholding performed with ImageJ. Right: Quantitative analysis with Student’s t test; *p* ≤ *0.05 (n=3).

### Experimentally induced EMT of mouse hair follicle stem cells reveals that *Vdr* signaling inhibits cell migration during normal cutaneous wound healing

The role of the VDR in OS migration prompted us to analyze whether migration is also affected in a different context such as skin wound repair. In addition to *in vitro* scratch assays of OS cells, we investigated cell migration of bulge stem cells in hair follicles, which is a well-established model whereby progenitor cells undergo EMT to repair wounds at distant regions (**Fig. 3A**)^13^. We used a lineage tracing approach to quantify the number of bulge stem cell progenitors that reside in the wound epicenter over time as a proxy for migration. To carry out these experiments, the RU486 inducible *keratin 15* (K15)-crePR1:*Vdr^flox^*^/^*^flox^*:Confetti reporter mouse line was created, and lineage traced for four days after wounding to capture the early stage of the process (**Fig. 3B**). Ablation of the *Vdr* within bulge stem cells increased the number of Confetti-labeled progenitors localized to the epicenter of wounds but not in individual hair follicles, resulting in enhanced wound closure (**Fig. 3C**, upper left and lower panels). Furthermore, bulge stem cell specific *Vdr* ablation resulted in thicker wound neoepidermis, an indication of increased EMT and fibrosis occurring following *Vdr* ablation. Mice with functional *Vdr* within bulge stem cells, on the other hand, had contiguous clonal streaks of Confetti-positive cells within individual hair follicles, indicating commitment of bulge stem cell progenitors down the hair follicle lineage (**Fig. 3C, upper left panel**). Furthermore, a subset of bulge stem cell progenitors was fate restricted to the epidermal lineage but only migrated to the wound periphery four days after injury. These findings suggest that the *Vdr* regulates the fate of bulge stem cells and inhibits the migration of progenitor cells in healthy animals, which are consistent with its role in blocking migration of cancer cells.

**Figure 3.**
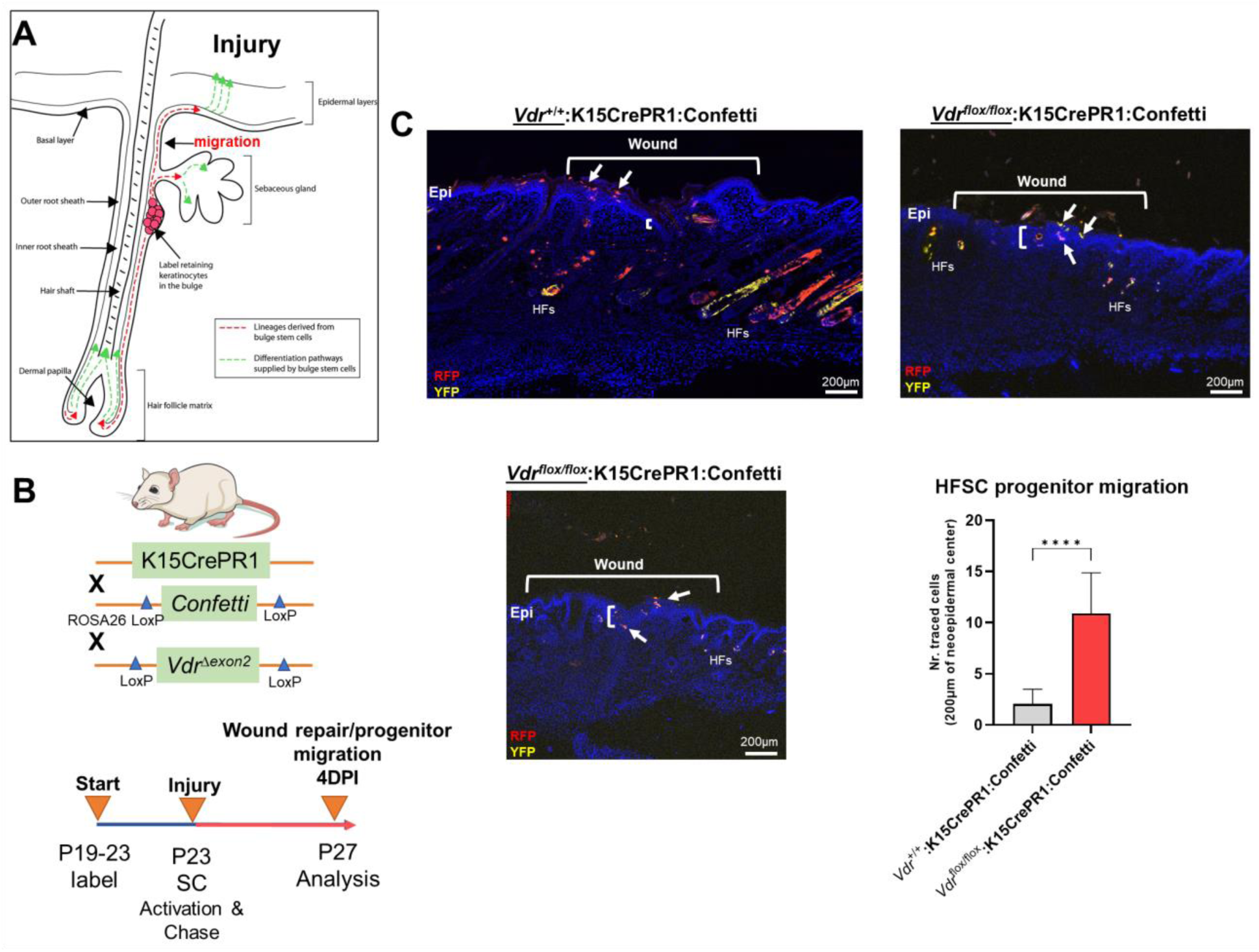
*Vdr* signaling inhibits cell migration *in vivo*. A) Hair follicles include oligopotent stem cells that, in response to environmental stimuli such as injury, can create hair follicle and/or epithelial progenitor cells. These cells can move to their final destinations to accomplish specific functions, such as fibrosis-based wound healing and hair follicle development. B) Top: In order to explore the involvement of the *Vdr* in injury-induced cell migration *in vivo*, a conditional *Vdr* ablation and hair stem cell Confetti reporter mice was developed. Bottom: Male experimental mice were treated topically with RU486 between the post-natal (P) ages of 19 and 23 days to activate Cre recombinase and label the initial pool of keratin 15 (K15)-positive hair follicle stem cells. Following 3-mm full-thickness skin lesions to activate stem cells (SCs), the animals were tracked for four days to assess the lineage and migration of progenitor cells (YFP, yellow fluorescent protein; RFP, red fluorescent protein). C) Left: At 4 days post-injury (DPI), the control animals’ wounds had not fully healed, as evidenced by the thin neo-epidermis, the neo-epidermal gap, and the migration of tagged progenitor cells to the wound margin. Right and Bottom: Experimental animals with *Vdr* ablation exhibit quicker wound closure, increased thickness (fibrosis) of the neo-epidermis, and the presence of tagged progenitor cells in the center of the neo-epidermis, indicating accelerated migration and improved tissue healing. In addition, *Vdr* deletion in hair stem cells hindered the development and contribution of progenitor cells to individual hair follicles, demonstrating that the *Vdr* plays an essential role in hair formation. D) Quantification of hair follicle stem cell progenitors in the wound’s epicenter. Student t test; *p* ≤ ***0.001 (n=4-5).

### Epigenomic and putative target gene and activity analysis reveals the integration of the VDR with neurotransmitter pathways in osteosarcoma cell models

Given the distinct features of the OS subtypes, we sought to better define molecular and epigenetic regulators within the models. We re-analyzed epigenomic data obtained from MG63 and LM7 cells, as well as data obtained from primary patient OS samples and those that had spread to lung tissues (GSE74230), by focusing on transcription factor (TF) motif binding, differential peaks, and gene ontology (GO) analysis of acetylated (ac) H3K27 histone modifications that reflect active transcription (**Fig. 4A**)^66^. TF factor motif analysis within H3K27ac peaks revealed 29 MG63-specific TF binding sites, including those for EOMES, PRDM1, EPAS1 (HIF2A), E2F6, and MYC/MAX, which reflect known early development, tumor-initiation, and stemness factors (**Fig. 4B & Worksheet 1**)^67^. GO analysis of motif-binding TFs confirmed their role in embryonic development, including brain development, in part due to the function of EOMES (also known as T box transcription factor 2, TBR2) in stem cell mediated neurogenesis (**Fig. 4C**)^68^. In addition, pathway analysis revealed the involvement of oxidative stress responses, a known regulator of cell migration^69^, and the PDGF signaling pathway, which integrates major signaling hubs to promote stem cell self-renewal^70^. Eighteen TF binding sites specific to LM7 were identified for TEAD, TR4, MYOG, SOX2, and E2F7, which support mesenchymal gene expression to disrupt ECM and cell-cell adhesions to promote cell migration (**Fig. 4B**)^71–73^. Of note, fibroblasts can transform into skeletal muscle cells via myogenin (MYOG)(45), indicating a potential regulatory link between metastatic OS and other musculoskeletal cancers such as rhabdomyosarcoma^74^, as revealed by the GO analysis (**Fig. 4C**). RUNX binding was observed in both OS cell lines, which is particularly important in OS development in its aberrant transcriptional control of osteoblast commitment^75^. Importantly, H3K27ac modifications were linked to VDR and RXR heterodimeric vitamin D response element (VDRE) binding in both cell models, indicating that 1,25(OH)_2_D regulates multiple functions of OS subtypes (**Fig. 4B**). Interestingly, GO Disease analysis revealed that subtype-specific TFs were strictly associated with cancers, whereas shared TFs were associated with a variety of cancer types as well as diseases such as type 2 diabetes mellitus, a risk factor for cancer development. In addition, by comparing H3K27ac with H3K4me1 (i.e., histone changes associated with enhancers) modifications and DNase I hypersensitive sites, the degree of active *versus* poised enhancer utilization within the OS subtypes was determined at the epigenome-wide level. In MG63 cells, there was a weaker correlation between H3K27ac and H3K4me1 than in LM7 cells (0.14 *versus* 0.44), indicating that enhancer elements are poised to help define a more undifferentiated cell state (**Fig. 4D left**)^76^. This was also shown by comparing overall H3K27ac marks between LM7 and MG63 cells, showing more gene activity in LM7 cells (**Fig. 4D right**).

**Figure 4.**
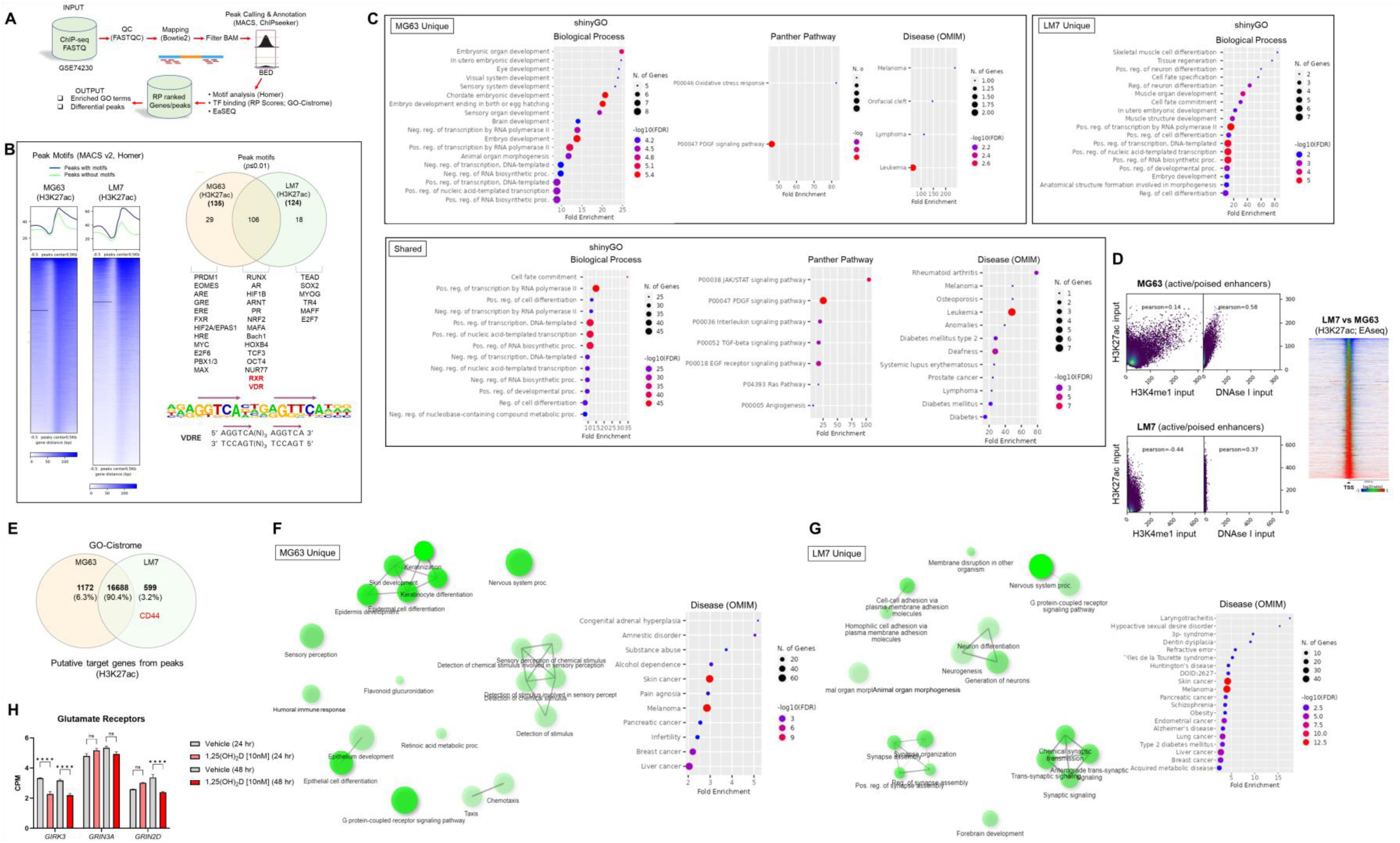
Epigenomic and putative target gene analysis of osteosarcoma cell models. A) Framework for ChIP-seq analysis and peak calling across MG63 and LM7 OS cell lines derived from GSE74230. Data quality specifications for FASTQ files processed with the Peak calling and Annotation pipeline. Using enriched GO keywords based on differential peaks, biological process, route, and disease functional annotations were generated. Using Homer and GO-Cistrome, motif analysis and transcription factor regulatory potential (RP) scores were obtained. Utilizing EaSeq, heatmaps and differential peaks were generated. B) Left: Homer was utilized to identify histone modification peaks (H3K27ac) and motif discovery between LM7 and MG63 cells. In HOMER, Summits.bed files from MACS2 (Model-based analysis of ChIP-Seq) were utilized to identify recognized motifs. The x-axis analysis utilized fixed-size peak discovery on histone markers. Right: A subset of peak motifs that are enriched in the LM7 and MG63 cell lines. Motifs having a *p*-value 0.01 are displayed. Below: Sequence of the consensus VDR direct repat 3 response element (VDRE) is shown. C) Gene ontology enrichment analysis of OS motifs with ShinyGo version 0.75. The x-axis represents fold enrichment, which indicates the extent to which genes of a certain pathway/biological process/disease are overrepresented. Fold enrichment is defined as the proportion of genes in the list that belong to a pathway divided by the proportion of genes in the background. The hues show the magnitude of the negative log10 of the false discovery rate (FDR), which indicates the likelihood that the enrichment occurred by coincidence (larger value means smaller FDR). D) Assessing epigenome-wide poised or active enhancers between MG63 and LM7 cells. Right: BED files were used to perform Pearson correlation analysis between H3K27ac/H3K4me1/DNase I peaks. Left: Heat map showing comparison of H3K27ac peaks between cell lines. E) Probable target genes generated from H3K27ac peaks identified using GO-Cistrome. F) Gene ontology enrichment analysis of MG63 OS H3K27ac mapped genes using ShinyGo version 0.75, with network analysis shown in green. G) Gene ontology enrichment analysis of identified LM7 OS H3K27ac genes utilizing ShinyGo version 0.75 Green depiction of network analysis. H) Glutamate receptor expression in MG63 cells as determined by RNA-seq. Two-way ANOVA Tukey’s multiple comparison test; *p* ≤ ****0.0001 (n=1).

Next, using GO-Cistrome, candidate target genes mediated by H3K27ac were identified and assessed (**Fig. 4E**)^77^. Approximately 13.3% and 9.3% of the H3K27ac peaks in LM7 and MG63 were located within 1KB of the promoter regions of genes (**Fig. S2**). There were 1172 and 599 H3K27ac-defining MG63 and LM7 specific gene signatures, with *CD44* identified as an active LM7 target gene (**Worksheets 2 & 3**). The GO Network and Disease analysis of MG63 putative genes was consistent with the motif analysis, which found correlations with epidermal differentiation, neuronal/sensory systems, and invasive malignancies (**Fig. 4F, S3 & File S1**). Compared to MG63 cells, LM7 H3K27ac-linked genes were significantly associated with neurogenesis and neurological disorders, as well as cell-cell adhesion molecules (**Fig. 4G, S3 & File S2**). Given the likelihood that osteosarcomas may act as neuroendocrine carcinoids, we examined transcription of ionotropic and metabotropic glutamate receptors following 1,25(OH)_2_D treatment of MG63 cells. Only three of the available 24 mammalian receptors and subunits were expressed in MG63 cells (i.e., *GIRK3*, *GRIN3A*, and *GRIN2D*), with two of the three receptors downregulated by 1,25(OH)_2_D (**Fig. 4H**) (GSE220948). The ionotropic glutamate [kainate] receptor 3 (*GIRK3*) was downregulated at both time points following 1,25(OH)_2_D treatment, whereas the ionotropic glutamate [NMDA] receptor subunit epsilon-4 (*GRIN2D*) was only downregulated after 48 hours and was identified as a putative H3K27ac-target gene in LM7 cells (**Worksheet 3**). These results imply that vitamin D_3_ may decreased tumor burden by inhibiting OS-nerve interactions.

### Vitamin D modulates MG63 tumorigenicity by inhibiting nonsense-mediated RNA decay in a SMG6-SMG7-dependent manner

To gain insights into the biological aspects of epigenetic modifications that are differentially represented in the OS subtypes, we utilized EaSeq (**Fig. 5A**). There were approximately 2,634 more H3K27ac gene-associated peaks than H3K4me1 peaks due to the fine-tuning regulatory effects of enhancers (**Worksheets 4 & 5**). By comparing 1,25(OH)_2_D resistant LM7 to MG63 cells, overrepresented target genes of H3K27 acetylation implicated the involvement of the mRNA surveillance pathway, but underrepresented genes engaged the HIF-1 and TNF signaling pathways, indicating a putative decrease in immunogenicity and immunosurveillance of LM7 metastatic cells (**Fig. 5B**). LM7 cells overexpressed NMD machinery components involved in the assembly of the surveillance complex at aberrant PTCs (e.g., *eRF1*, *UPF2*, *UPF3*) and cleavage of RNA templates involving exo/endonucleolytic pathways (*SMG5*, *SMG6*, *SMG7*) (**Fig. 5C**). Interestingly, the RNA-binding protein MUSASHI (encoded by *MSI1*), which lies upstream of the surveillance complex, was increased in LM7 cells indicating the regulation of translation of defective transcripts vulnerable to nonsense-mediated translation repression (NMTR)^78^.

**Figure 5.**
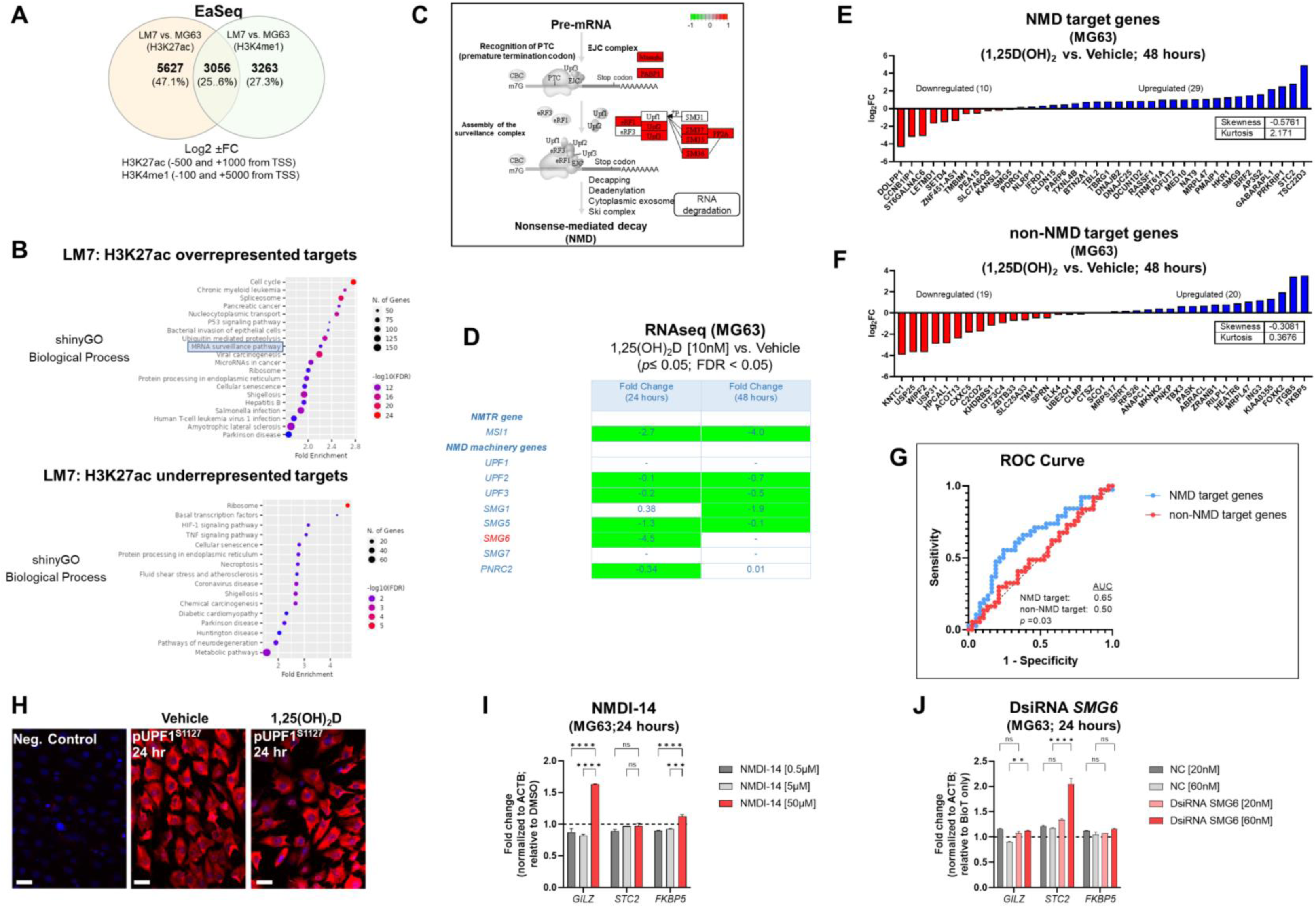
Differential peak analysis of osteosarcoma lines reveals overrepresentation of the NMD pathway. A) Predicting distinct H3K27ac and H3K4me1 peaks in OS cells using the EaSeq software. Greater differential peaks between cell lines and histone markers based on Log2 fold variations inside designated regions relative to the transcription start site (TSS). B) Gene ontology enrichment analysis of LM7 OS H3K27ac overrepresented and underrepresented putative target genes with ShinyGo version 0.75. The mRNA surveillance pathway is highlighted among shown biological processes. C) Pathview visualization of the nonsense-mediated RNA decay (NMD) pathway in relation to the H3K27ac target genes affected in LM7 cells. Targets that were overrepresented at the exon junction and surveillance and cleavage complexes are shown in red. D) RNA-Seq evaluation of nonsense-mediated translation suppression and NMD machinery genes in MG63 cells. Notable was SMG6, which is responsible for the cleavage of pre-mRNA, resulting in its degradation. E) In MG63 OS cells, 1,25(OH)_2_D selectively upregulates known NMD target genes. MG63 cells treated with 10nM 1,25(OH)_2_D for 48 hours exhibited an elevated 3:1 ratio of upregulated known NMD target genes relative to non-NMD target genes. The RNA-seq data set’s statistically significant genes (FDR ≤ 0.05) were compared. The symmetry of the data distribution was also characterized by descriptive statistics (skewness/kurtosis). F) In MG63 OS cells, 1,25(OH)_2_D does not selectively upregulate non-NMD target genes. After treatment with 1,25(OH)_2_D, the ratio of upregulated to downregulated genes in a random sample of non-NMD target genes is 1:1. G) Analysis of the receiver operating characteristic (ROC) curve for statistical comparisons and identification of NMD or non-NMD target gene classes. To compare the two classifiers, the area under the ROC curve (AUC) was computed, which quantifies the overall performance of the test to discriminate between genes in the presence of 1,25(OH)_2_D that are NMD-target or non-NMD target genes relative to the null hypothesis (0.5). H) Phosphorylated UPF1^S1127^ expression in MG63 cells after 24 hours of 10nM 1,25(OH)_2_D. Bar = 20µm. I) NDM inhibitor 14 (NMDI-14) treatment of MG63 for 24 hours. Two-way ANOVA test with Tukey’s multiple comparisons (n=3); *** *p ≤* 0.001, **** *p ≤* 0.0001. J) DsiRNA *SMG6* knockdown of MG63 cells for 24 hours. Negative control (NC) duplexes used for comparisons. All conditions were compared to transfection reagent (BioT) only treatments. Two-way ANOVA test with Tukey’s multiple comparisons (n=3); ** *p ≤* 0.01, **** *p ≤* 0.0001.

We next investigated vitamin D_3_ regulation of NMD in MG63 and LM7 cells. We observed a statistically significant downregulation of autoregulatory NMD machinery as well as NMTR genes following 10nM 1,25(OH)_2_D treatment of MG63 cells (**Fig. 5D**)^79^. Interestingly, there was a statistically significant downregulation of *SMG5* among a subset of detectable NMD genes in LM7 cells treated with 10nM 1,25(OH)_2_D for 48 hours as well (**Fig. S4A**). Next, we examined the impact of 1,25(OH)_2_D on known NMD target genes. On the basis of four investigations in which NMD effectors were inhibited in human cervical cancer cells^79–82^, we compiled a list of NMD-target and non-NMD-target genes (**Fig. S5**). Based on this analysis, more than 4,000 genes were identified as NMD nontarget genes, whereas more than 900 genes were recognized as NMD targets by at least one study; whereby, 50 genes were classified as NMD core targets by all studies. These curated NMD-target and nontarget genes were used to analyze the RNA-seq dataset generated from MG63 cells. Thirty nine of the 50 core NMD-target genes were mapped to MG63 cells, and overexpressed in a 3:1 ratio (−0.6 skewness and 2.2 kurtosis) relative to those that were downregulated in response to 1,25(OH)_2_D treatment (**Fig. 5E**). TSC22 Domain Family Member 3 (*TSC22D3*), also known as *GILZ*, which encodes a glucocorticoid-inducible anti-proliferation transcription factor, was the most elevated NMD target in MG63 OS cells^83^. Stanniocalcin-2 (*STC2*), which is involved in normal osteoblast differentiation^84^, was the second most elevated NMD-target gene in response to 1,25(OH)_2_D, as well as *CLDN15* and *BTN2A1*, which may promote cell-to-cell adhesion to limit migration^85^ and the expression of a phosphoantigen to enhance T cell activity^86^, respectively. In contrast, a multiple-comparison study of NMD nontarget genes demonstrated a 1:1 (−0.3 skewness and 0.4 kurtosis) association between overexpressed and downregulated genes (**Fig. 5F**). In addition to the descriptive statistics, a receiver operating characteristic (ROC) curve analysis was performed to investigate the capacity to distinguish between genes that are one of two classifiers (i.e., NMD target or non-target genes) in the presence of 1,25(OH)_2_D relative to the null hypothesis (area under the curve, 0.5). The C statistics revealed statistical significance (*p* = 0.03) between the two classifiers based on ROC curve analysis (**Fig. 5G**).

To gain a better understanding of how 1,25(OH)_2_D controls the NMD pathway in MG63 cells, we investigated UPF1, which is a key RNA helicase that assembles the upstream surveillance complex at the exon junction complex of aberrant mRNAs dictated by its phosphorylation status (**Fig. 5C**)^87, 88^. After 24 hours of 1,25(OH)_2_D treatment, there was no significant change in UPF phosphorylation compared to vehicle treated MG63 cells (**Fig. 5H**), suggesting that changes in *UPF2*/*UPF3* expression may be the key NMD upstream factors regulated by 1,25(OH)_2_D (**Fig. 5D**). To better understand the role of downstream NMD pathway factors on regulation of NMD-target (e.g., *GILZ*, *STC2*) and non-target (e.g., *FKBP5*) genes in MG63 cells, we pharmacologically and genetically ablated components of the RNA degradation complex. First, we utilized the small molecule NMD inhibitor, NDMI-14, which prevents SMG7-UPF1 interactions^89^, and observed a modest increase in *GILZ* after 24 hours of treatment (**Fig. 5I**). There was a small, yet significant increase in *FKBP5*, which may reflect stress responses of high NMDI-14 concentrations, given its function as a Hsp90-associated co-chaperone^90^. Next, we utilized dicer-substrate short interfering RNAs (DsiRNA) to knockdown (KD) *SMG6* in MG63 cells and observed a modest increase in *STC2*, as well as a statistically significant, yet small increase in *GILZ* (**Fig. 5J & S4B**). Among the conditions tested, the findings suggest that *GILZ* is dependent on both SMG7 and SMG6, while *STC2* is modulated by SMG6 only, and that 1,25(OH)_2_D exhibited a greater impact on NMD target genes likely reflecting its simultaneous effects on multiple upstream and downstream components of the NMD pathway to suppress OS.

### LM7 resistance to 1,25(OH)_2_D is dictated by high *SNAI2* and low *VDR* levels

To better understand the functional role of 1,25(OH)_2_D on metastasis, the transcriptional responses of EMT genes on highly metastatic LM7 cells were compared to those of MG63 cells. The decrease in expression of *SNAI2* in MG63 cells was less pronounced in LM7 cells following 1,25(OH)_2_D treatment, while *CD44* expression was unaffected (**Fig. 6A**). There was a concentration-dependent effect to 1,25(OH)_2_D on *MMP3* expression 24 hours after treatment, with 10nM increasing *MMP3* levels, suggestive of resistance. In addition, the dose and duration of 1,25(OH)_2_D in LM7 cells resulted in a maximal 2-fold overexpression of *SOD2*, a potent antioxidative enzyme that inhibits tumor progression via reduced free radical production, which was less pronounced compared to MG63 cells (**Fig. 6B**). Interestingly, there was significant overexpression of the mitochondrial monooxygenase, *CYP24A1*, which catabolizes and deactivates 1,25(OH)_2_D via 24-hydroxlation, in LM7 cells after 1,25(OH)_2_D treatment (**Fig. 6B**). Also, by evaluating the relative levels of *VDR* and *SNAI2* transcripts, MG63 cells were characterized as *VDR*^high^*SNAI2^l^*^ow^ whereas LM7 cells were *VDR*^low^*SNAI2*^high^, providing more evidence for 1,25(OH)_2_D resistance in LM7 cells (**Fig. 6C**). By increasing the duration of 1,25(OH)_2_D from 48 to 96 hours, LM7 cells overcame the resistance as depicted by a significant decrease in CD44 expression levels (**Fig. 6D**).

**Figure 6.**
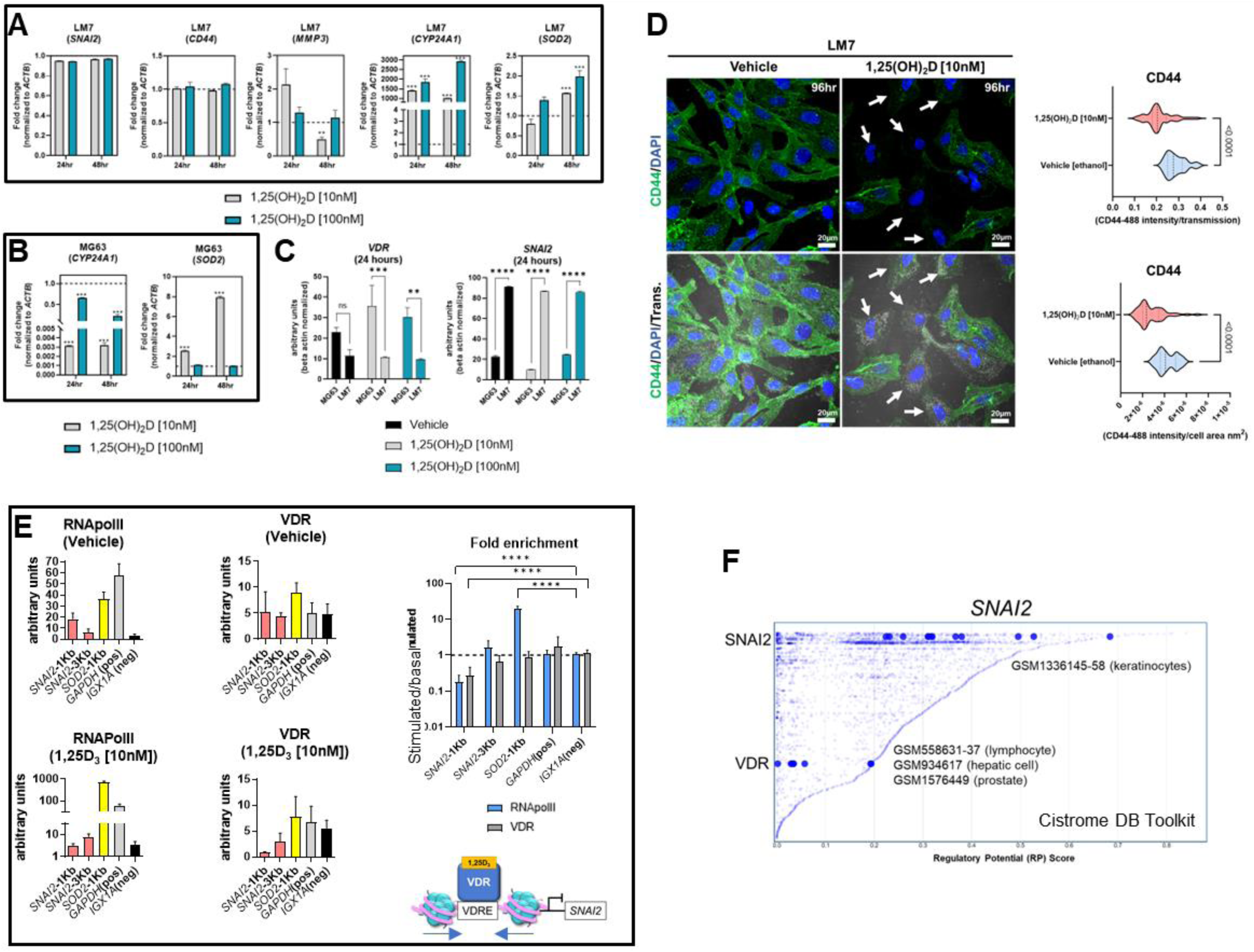
LM7 metastatic cells are resistant to 1,25(OH)_2_D and *SNAI2* is a VDR-target gene. A) qPCR study of EMT, vitamin D, and anti-oxidative transcripts in LM7 cells following treatment with 1,25(OH)_2_D. Two-way ANOVA Multiple comparisons using Tukey’s test; *p* ≤ **0.01, ***0.001 (n=3). B) qPCR study of vitamin D and anti-oxidative transcripts in MG63 cells following treatment with 1,25(OH)_2_D. Two-way ANOVA Tukey’s test for multiple comparisons; *p* ≤ 0.001 (n=3). C) Normalized *VDR* and *SNAI2* gene expression in LM7 and MG63 cells. Two-way ANOVA Test for multiple comparisons using Tukey’s method; *p* ≤ **0.01, ***0.001, ****0.0001 (n=3). D) 96 hours of 10nM 1,25(OH)_2_D treatment decreases CD44 expression levels in LM7 cells. Arrows depict cells with largest differences in CD44 expression. Unpaired T test (n=4). E) ChIP study of VDR-RNApol2 chromatin interactions in 1,25(OH)_2_D-treated MG63 cells. A two-way ANOVA was performed on the fold-change between stimulated and baseline states. Dunnett’s multiple comparison test relative to *IGX1A*; *p* ≤**0.01, ***0.001, ****0.0001 (n=3). F) Analysis of transcription factors that regulate *SNAI2* using the Cistrome DB Toolkit. The static plot with RP scores on the X-and Y-axes depicts transcription factors, whereas the blue dots represent individual public ChIP-seq data sets. Listed next to each transcription factor are public GEO data series.

### VDR-mediated suppression of the EMT transcriptional repressor *SNAI2* occurs via chromatin condensation

Based on the ATACseq and qPCR data we hypothesized that *SNAI2* is a direct VDR target gene. Using NUBIscan^91^, a putative VDRE direct repeat (DR3) binding site was found within 1KB upstream of the transcriptional start site (TSS) exclusively in the *SNAI2* locus, but not in the *SOD2* locus (**Fig. S6**). MG63 cells treated with 1,25(OH)_2_D or vehicle were subjected to conventional chromatin immunoprecipitation quantitative PCR (ChIP-qPCR) to measure VDR binding and transcriptional activity mediated by RNAPolII at the *SNAI2* distal and proximal promoter regions (**Fig. 6E**). After 1,25(OH)_2_D treatment, there was decreased enrichment of the VDR and RNAPolII in the proximal promoter region of *SNAI2* compared to the intergenic control (*IGX1A*). After 1,25(OH)_2_D treatment, *SOD2* transcriptional activity increased, although this was not VDR-dependent. Given the likelihood that VDR regulates *SNAI2* transcriptional activity directly, the Cistrome DB toolset was used to identify additional VDR:SNAI2 chromatin interactions^92^. Three GEO datasets were identified when the 1KB domain of the *SNAI2* TSS was investigated for potential VDR binding in lymphocytes, hepatocytes, and prostate cancer cells (**Fig. 6F**). The identification of putative SNAI2 binding sites in keratinocytes, which are highly migratory cells of the body (**Fig. 6F**)^69^, also suggests the possibility of intracrine feedback effects that maintain high *SNAI2* levels in LM7 cells. Overall, LM7 cells are more resistant to 1,25(OH)_2_D than MG63 cells, which may be a direct result of CYP24A1 catabolic effects on ligand and decrease VDR responses at the *SNAI2* locus.

### H3K27 is differentially acetylated at the *SNAI2* and *VDR* loci in highly metastatic and low metastatic osteosarcomas

To gain a better understanding of *SNAI2* and *VDR* regulation in osteosarcomas, we evaluated H3K27ac and H3K4me1 epigenetic alterations in a panel of highly metastatic and low metastatic human OS cell lines (GSE74230)^66^. Compared to low metastatic cell lines MG63, HOS, and Hu09, the highly metastatic OS cell lines LM7, MNNG, and M112 had higher average H3K27ac peak values at the *SNAI2* TSS (**Fig. 7A,C**), which corresponded to the degree of DNase hypersensitivity (lower tracks). Intriguingly, there were no differences in *SNAI2* H3K27ac peak values when comparing two OS patient primary bone and metastasized lung tumor samples (**Fig. 7B,C**), possibly due to reduced migration in the lung tumor microenvironments compared to *in vitro* conditions of cell lines. In contrast, the low metastatic cell lines revealed greater H3K27ac peaks at the *VDR* TSS than the highly metastatic cell lines (**Fig. 7D**). Similarly, there were no differences in H3K27ac and H3K4me1 epigenetic modifications at the *VDR* TSS and distal 5kb regions between OS patient primary bone or lung tumors (**Fig. 7E**). These findings show that metastasizing OS cells exhibit increased *SNAI2* H3K27ac, but low metastatic lines exhibit increased *VDR* H3K27ac, potentially making them more responsive to 1,25(OH)_2_D, a pattern that was abolished in metastasized patient lung tumor samples.

**Figure 7.**
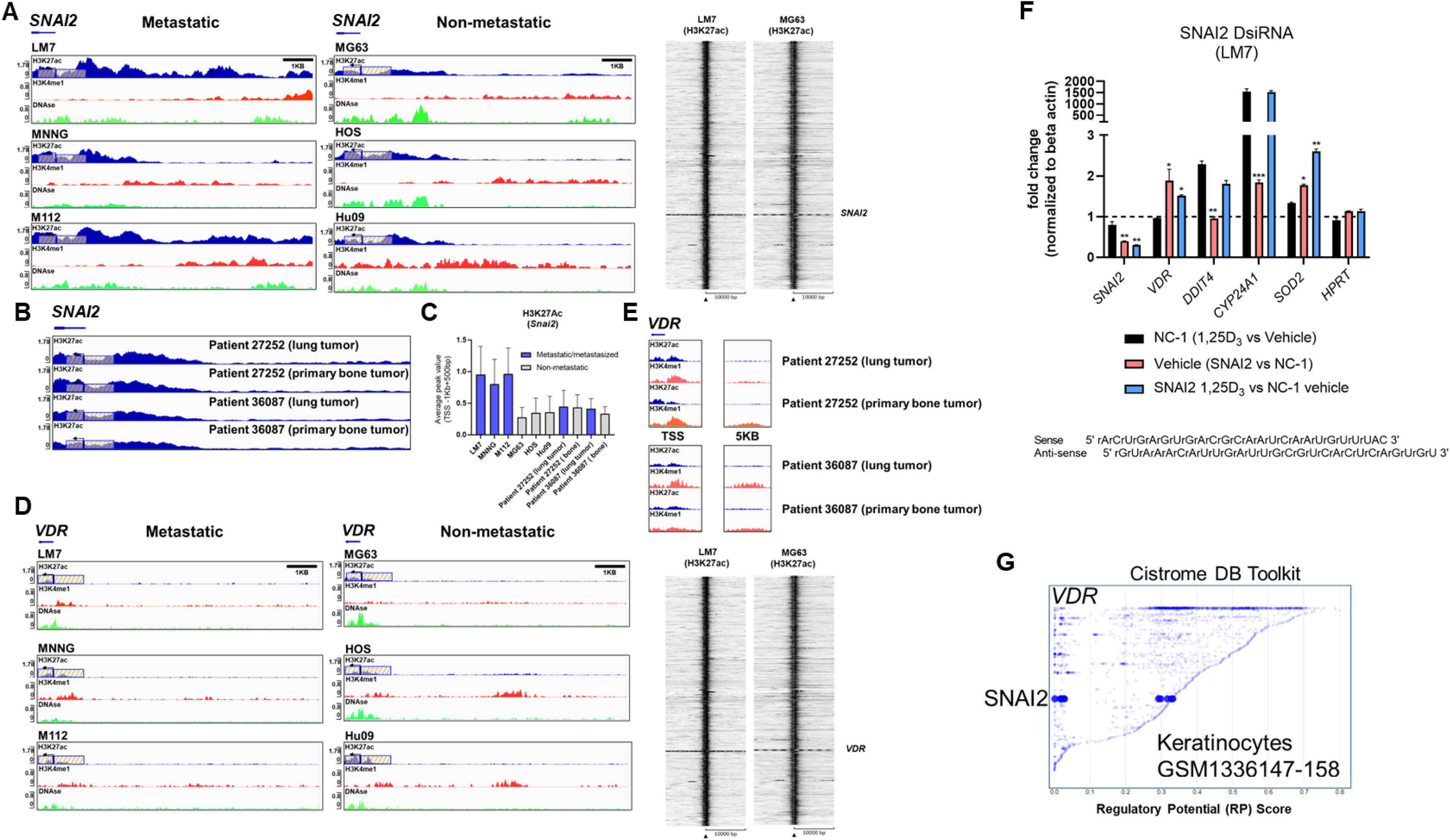
Differential H3K27 acetylation at the *SNAI2* and *VDR* loci between OS subtypes and mediation of antioxidative pathways by vitamin D and SNAI2. A) ChIP-seq studies and peak calling for the *SNAI2* gene in metastatic and non-metastatic OS lines from GSE74230. Evaluation of H3K27ac, H3K4me1, and DNase I hypersensitive sites 500bp downstream and 1000bp upstream of the transcription start site (TSS). Quantification of peak values in H. Right: Using EaSeq, a heat map of H3K27ac was created at the TSS. Using IGV’s auto-scale of BED files, all scales reflect the minimum and maximum reads in that window. B) ChIP-seq study of H3K27ac changes at the *SNAI2* gene in primary bone and lung tumor tissues from OS patients. Samples generated from GSE74230. C) Quantification of the average peak values of OS cell lines that are poorly metastatic, metastatic, and patient samples. D) ChIP-seq studies and peak calling for the *VDR* gene in metastatic and non-metastatic OS lines from GSE74230. Evaluation of H3K27ac, H3K4me1, and DNase I hypersensitive sites 500bp downstream and 1000bp upstream of the transcription start site (TSS). Using IGV’s auto-scale of BED files, all scales reflect the minimum and maximum reads in that window. Using EaSeq, a heat map of H3K27ac was created at the TSS. E) ChIP-seq study of H3K27ac changes at the *VDR* gene in primary bone and lung tumor tissues from OS patients. Derived samples from GSE74230. F) Gene responses to DsiRNA-mediated SNAI2 knockdown in LM7 cells. The sense and antisense DsiRNA SNAI2 sequences are displayed below. Tukey’s multiple comparison test for two-way ANOVA; *p* ≤ *0.05, **0.01, ***0.001 (n=4). G) Cistrome DB Toolkit study of *VDR*-regulating transcription factors such as SNAI2. The static plot with RP scores on the X-and Y-axes depicts transcription factors, whereas the blue dots represent individual public ChIP-seq data sets. Listed next to each transcription factor are the public GEO data series.

### DsiRNA knockdown of *SNAI2* reveals antioxidative responses mediated by vitamin D_3_

To better comprehend the plausibility of VDR-SNAI2 reciprocal regulation, we utilized DsiRNA to KD *SNAI2* in LM7 cells (**Fig. S7,S8**). In order to explore the transcriptional consequences of DsiRNA KD of *SNAI2* in LM7 cells, a comparative qPCR analysis was conducted. Negative control (NC-1) DsiRNA treated samples exhibited the predicted responses to 1,25(OH)_2_D, notably the downregulation of *SNAI2* and the overexpression of *DDIT4* (a potent VDR-dependent inhibitor of mTOR^51^), *CYP24A1*, and *SOD2* (**Fig. 7F, black bars**). Comparing vehicle treatment across SNAI2 and NC-1 DsiRNA KD samples, the expected >60% KD of *SNAI2* and an intriguing elevation of *VDR, CYP24A1* and *SOD2* were detected, indicating direct dependence and negative regulation by SNAI2 (**Fig. 7F, red bar**). To provide additional evidence of potential SNAI2 interactions at the *VDR* promoter site, we re-evaluated the Cistrome DB Toolkit and identified the datasets GSM1336147-158, which illustrate putative interactions in highly motile skin keratinocytes (**Fig. 7G**)^93^. In contrast, *SNAI2* DsiRNA KD had no effect on *DDIT4*, suggesting SNAI2-independent signaling. The effect of *SNAI2* DsiRNA KD on 1,25(OH)_2_D responses revealed further *SNAI2* downregulation, providing additional evidence that *SNAI2* is VDR-dependent (**Fig. 7F, blue bar**). Due to an absence of autocrine VDR signaling, there was no significant difference in *VDR* expression between vehicle (**Fig. 7F, pink bar**) and 1,25(OH)_2_D *SNAI2* DsiRNA KD (**Fig. 7F, blue bar**) responses. After 1,25(OH)_2_D treatment, both *DDIT4* and *CYP24A1* levels returned to NC-1 levels when *SNAI2* was inhibited, demonstrating once again that *DDIT4* is independent of SNAI2, but *CYP24A1* was found to be co-dependent on VDR and SNAI2. Intriguingly, given the maximal effect of most genes following *SNAI2* DsiRNA KD and 1,25(OH)_2_D treatment relative to the control, *SOD2* levels continued to increase (**Fig. 7F, blue bar**), indicating a repressive effect of SNAI2 on *VDR* expression. Overall, SNAI2 can modulate 1,25(OH)_2_D responses, thereby linking EMT, ROS, and vitamin D_3_ signaling into a novel OS network^94, 95^.

### Vitamin D_3_ modulates SOD2 intracellular localization to inhibit osteosarcoma mitochondrial oxidative stress

Given that ROS regulates cell migration and vitamin D_3_ induces *SOD2* expression^69^, the epigenomic regulation of *SOD2* in OS cell lines was evaluated. At rest, LM7 cells displayed increased H3K27 acetylation at the *SOD2* TSS compared to MG63 cells, with augmentation of intronic and intergenic H3K4me1 modification (**Fig. 8A**), indicating a pronounced antioxidative response mediated by active enhancer elements. ATACseq analysis indicated similar degree of chromatin accessibility at the *SOD2* TSS in MG63 cells treated with 1,25(OH)_2_D or vehicle (**Fig. 8B**), suggesting that *SOD2* mRNA levels are likely regulated by post-transcriptional mechanisms^48^. In addition, we analyzed SOD2 protein and observed that it was mainly located in the nucleus of MG63 cells under basal conditions and was seldom associated with VDAC1-positive mitochondria (**Fig. 8C**). In contrast, 1,25(OH)_2_D directed the mitochondrial localization of SOD2, possibly allowing it to perform its primary ROS detoxifying functions (**Fig. 8C,D**). Because of these OS subtype differences, we investigated ROS responses to 1,25(OH)_2_D treatment. Pentafluorobenzenesulfonyl fluorescein, a hydrogen peroxide (H_2_O_2_) reporter dye, revealed that 10nM 1,25(OH)_2_D for 24 hours promoted efficient scavenging of H_2_O_2_ in MG63 cells (**Fig. 8E,F**)^69^. Interestingly, in agreement with the epigenetics of *SOD2*, LM7 metastatic cells had an 18-fold increase in H_2_O_2_ compared to MG63 cells (**Fig. 8G**), whereas 1,25(OH)_2_D was also able to suppress H_2_O_2_ levels, implying that anti-oxidative pathways regulated by vitamin D_3_ are shared in both OS subtypes (**Fig. 8H**).

**Figure 8.**
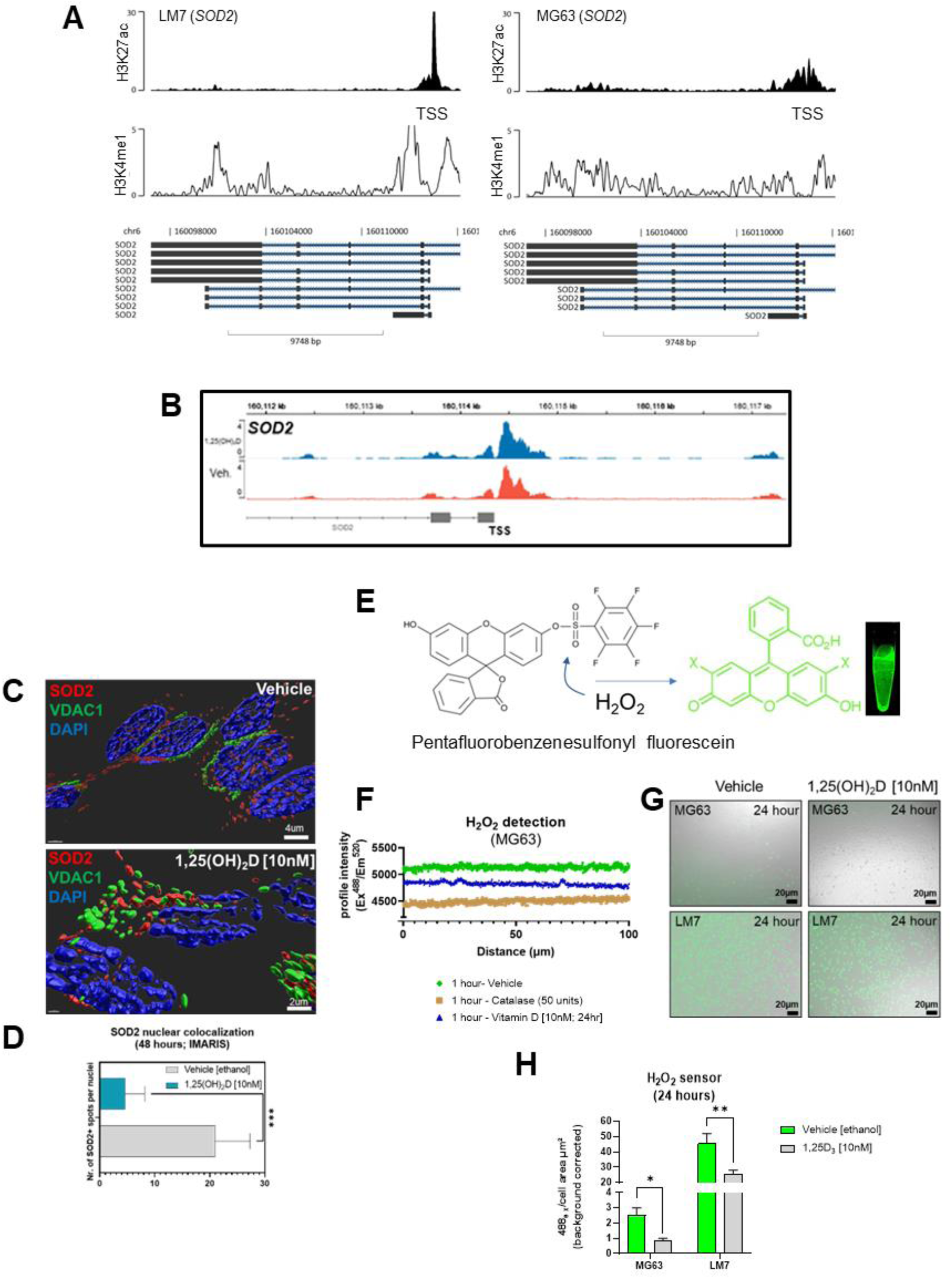
Vitamin D modulates nuclear-localized SOD2-to-*SOX2* axis to antagonizes osteosarcoma oxidative stress, iron transport and pluripotency to dictate migratory status. A) ChIP-seq study of the *SOD2* region in LM7 and MG63 cells. B) ATAC-seq study of MG63 cells at the *SOD2* gene in the presence or absence of 1,25(OH)_2_D (10nM, 24 hours). C) Confocal imaging of MG63 OS cells with SOD2 and mitochondrial VCAC1. Imaris surfaces for rendering 3D surfaces. D) Quantitative analysis of nuclear SOD2. Student T test; *p* ≤ ***0.001 (n=5). E) Schematic of the sensor pentafluorobenzenesulfonyl fluorescein for hydrogen peroxide (CAS 728912-45-6). F) Intensity of the pentafluorobenzenesulfonyl fluorescein confocal profile over MG63 cells. Vehicle, catalase (to breakdown hydrogen peroxide), and 1,25(OH)_2_D were used to treat cells. G) Representative confocal images demonstrating the formation of hydrogen peroxide by MG63 and LM7 cells under various circumstances. H) Quantification of the presence of hydrogen peroxide in MG63 and LM7 cells. Tukey’s multiple comparison test, two-way ANOVA; *p* ≤ *0.05, **0.01 (n=4).

### Vitamin D suppresses free radical generation and desmoplasia in low metastatic osteosarcoma and promotes MET of highly metastatic osteosarcoma in soft agar spheroids

Next, we used soft agar spheroids to determine if the antioxidative responses of 1,25(OH)_2_D are retained in the 3D environment. MG63 cells grew as round, well-circumscribed spheroids in soft agar for up to 14 days in culture (**Fig. 9A**). When spheroids were treated with 10-50nM 1,25(OH)_2_D, there was a significant dose-dependent shrinkage (**Fig. 9B**). Vehicle-treated MG63 spheroids exhibited a necrotic core (i.e. Zombie Aqua-positive), commonly seen in intact tumors, and surrounding cells with high 4-hydroxynonenal (4-HNE) expression, a marker for lipid peroxidation and free radicals (**Fig. 9C**, **bottom orthogonal views**). In contrast, 1,25(OH)_2_D reduced the expression of 4-HNE in spheroids as well as cells expressing Zombie Aqua, indicating the presence of viable cells (**Fig. 9D**). Given that vitamin D_3_ can elicit anti-fibrotic responses and that cancer is a fibrotic disease known to obstruct, in part, immune cell infiltration^12, 96^, we examined collagen protein and mRNA expression in MG63 spheroids and cells. There are 28 types of human collagens encoded by 43 genes, and 22 collagen genes were expressed in MG63 cells. 1,25(OH)_2_D inhibited the transcription of half of the collagen types detected (11 out of 22), and suppressed mature type 1 collagen within spheroids (**Fig. 9E,F**). Unlike MG63 cells, LM7 metastatic cells grown in soft agar did not form spheroids but instead remained as single viable cells with minimal growth after 14 days (**Fig. 9G**). In contrast to MG63 cells, 1,25(OH)_2_D promoted the clustering of Zombie Aqua-negative LM7 cells in soft agar, implying the induction of mesenchyme-to-epithelial transition (MET) as a means of inhibiting metastasis (**Fig. 9H**). Overall, the findings indicate that vitamin D_3_ inhibits free radical and collagen production in a cell intrinsic manner, and initiates cell-cell adhesion in a highly metastatic OS model.

**Figure 9.**
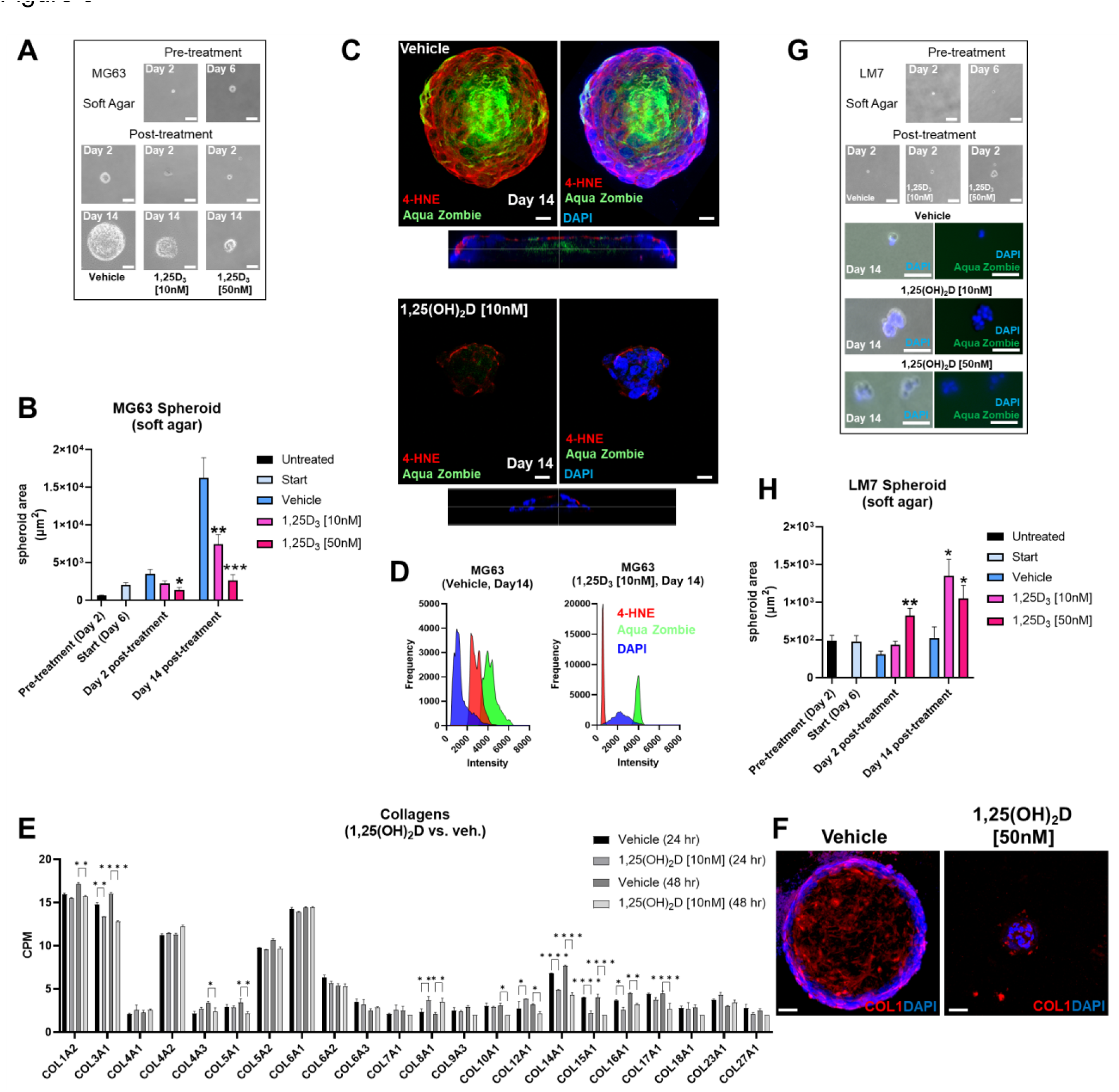
Vitamin D reduces free radical production, desmoplasia, and lipid peroxidation in non-metastatic osteosarcoma spheroids, while promoting MET in metastatic osteosarcoma spheroids in soft agar. A) MG63 cell spheroid assay with soft agar. Bar = 100µm. B) Measurement of A. Tukey’s multiple comparison test for two-way ANOVA; *p* ≤ *0.05, **0.01, ***0.001 (n=4). C) Confocal imaging of MG63 spheroids with the lipid peroxidation indicator 4-HNE and the dead cell dye Zombie Aqua. Orthogonal perspective on the base of each composite. Bar = 20µm D) Quantification of C by evaluating the frequency and intensity of every marker. E) RNA-seq study of collagen expression before and after 1,25(OH)_2_D treatment in MG63 cells. Tukey’s multiple comparison test; *p* ≤ *0.05, **0.01, ***0.001, ****0.0001 (n=2). F) Type 1 collagen protein expression in MG63 spheroids. Bar = 40µm G) Spheroid assay with soft agar for LM7 cells. 2-day bar = 100µm; 14-day bar = 20µm H) Measurement of F. Tukey’s multiple comparison test, two-way ANOVA; *p* ≤ *0.05, **0.01 (n=4).

### Calcipotriol, a clinically relevant vitamin D_3_ analogue, inhibits osteosarcoma migration, desmoplasia and growth

Because of its lower calcemic activity in regulating calcium metabolism for clinical application, we investigated the effects of calcipotriol, a synthetic analogue of vitamin D_3_ (**Fig. 10A**)^59^. The effects on cell proliferation were first assessed using the MTT assay, which revealed that calcipotriol inhibited MG63 cell proliferation starting at 1nM lasting 48 hours (**Fig. 10B**). On the other hand, calcipotriol had no effect on LM7 proliferation during the treatment period. Instead, calcipotriol significantly inhibited LM7 migration compared to 1,25(OH)_2_D treatment (i.e., a ∼50 vs 26% suppression; **Fig. 10C & 2E**). Moreover, when compared to 1,25(OH)_2_D, calcipotriol significantly inhibited the growth of MG63 spheroids in soft agar (**Fig. 10D**), with concomitant suppression of desmoplasia through decreased type 1 and 3 collagen production (**Fig. 10E**). Interestingly, 1,25(OH)_2_D and calcipotriol both inhibited the laminar deposition of both types of fibrillar collagens, removing the outer surface of collagen “tracks” known to aid in EMT, metastasis, and the blocking of immune cells (**Fig. 10E, arrows**)^97^. The magnitude of calcipotriol effects on MG63 was not as pronounced as for 1,25(OH)_2_D, according to qPCR analysis (**Fig. 10F, upper**). For example, after 48 hours, the *SOD2* level induced by 1,25(OH)_2_D was close to 8-fold in MG63 cells (**Fig. 6B**), but never exceeded 1.8-fold after calcipotriol treatment. Furthermore, calcipotriol treatment had no effect on *SNAI2* in the study’s concentration range or duration when compared to 1,25(OH)_2_D treatment of MG63 cells (**Fig. 1B**), reflecting its function as an unique VDR agonist. In contrast, qPCR analysis of LM7 cells revealed that the transcriptional readout was significantly enhanced after calcipotriol treatment compared to 1,25(OH)_2_D-treated cells (**Fig. 10F, lower**). Calcipotriol, for example, reduced *SNAI2* expression by half, a significant reduction compared to 1,25(OH)_2_D responses (**Fig. 6A**). Similarly, calcipotriol stimulated a large transcriptional increase of *GILZ* 24 hours after treatment, indicating that the NMD pathway may have dynamic and rapid effects on aberrant gene expression. Overall, these findings show that calcipotriol can enhance the EMT and growth suppression of OS subtypes.

**Figure 10.**
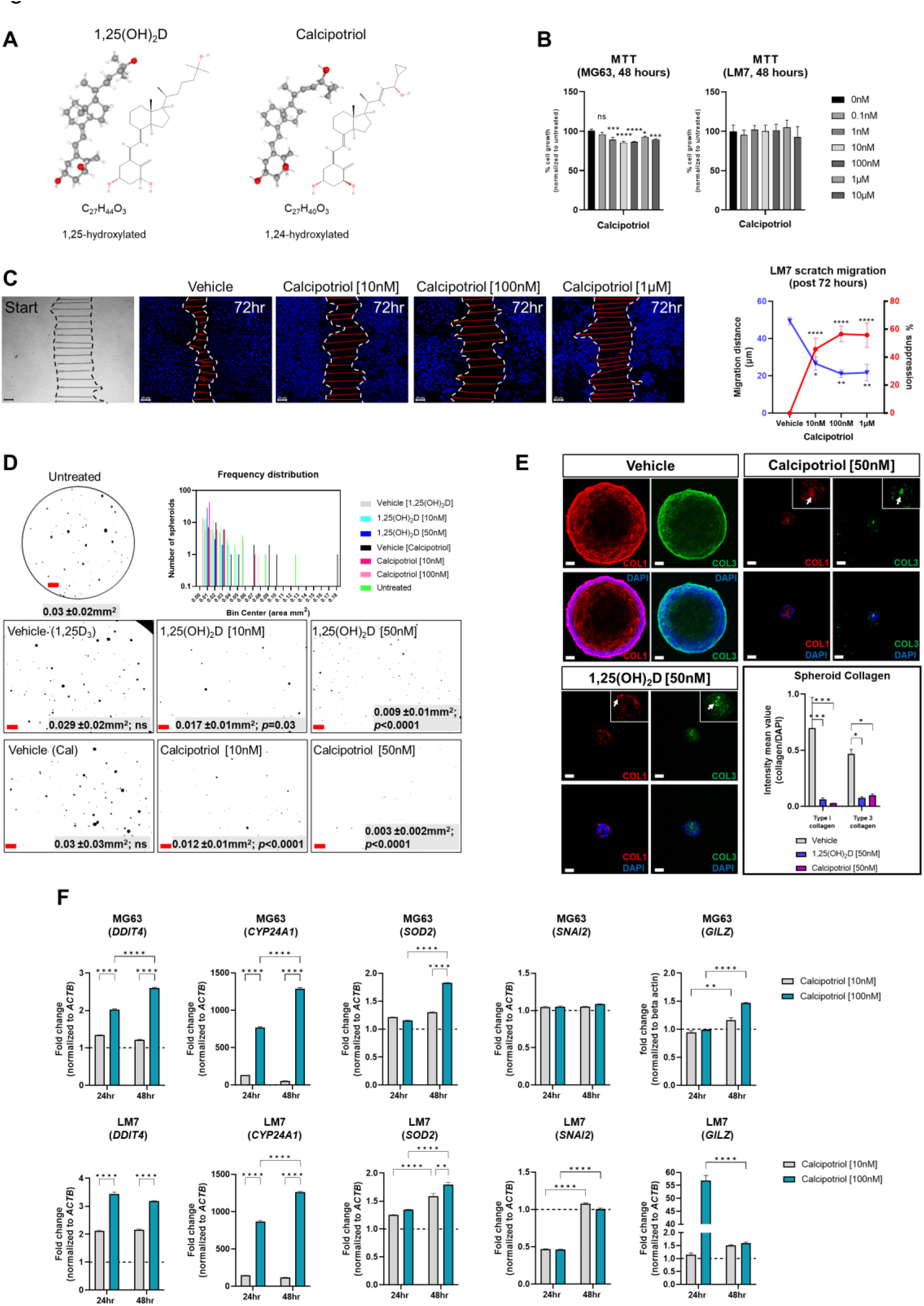
Effects of vitamin D3 synthetic analogue calcipotriol on MG63 and LM7 cells. A) The structural formula of calcipotriol, a synthetic vitamin D_3_ analogue and VDR agonist. B) MTT (3-[4,5-dimethylthiazol-2-yl]-2,5 diphenyl tetrazolium bromide) assay to measure cell proliferation 48 hours post-treatment. One-way ANOVA combined with Dunnett’s multiple comparison analysis; *p* ≤ *0.05, ***0.001, ****0.0001 (n=2). Bar = 20µm C) Scratch migration assay employing LM7 cells treated with calcipotriol. For each condition, the distance of the scratch was measured across the scratch. Tukey’s multiple comparison test for two-way ANOVA; *p* ≤ *0.05, **0.01, ****0.0001 (n=4). Bar = 20µm D) Frequency distribution of spheroids in soft agar treated with calcipotriol. Composite represent the background corrected, threshold-applied ImageJ post-processed images. One-way ANOVA; Šídák’s multiple comparisons test versus untreated (n=3). Bar = 1mm E) 1,25(OH)_2_D and calcipotriol decrease collagen production and laminar deposition within MG63 spheroids. MG63 cells in soft agar for 14 days were immunostained for types 1 and 3 collagen. Arrows depict collagen aggregations in the magnified insets. Two-way ANOVA Multiple comparisons using Tukey’s test; *p* ≤ *0.05, ***0.001 (n=3). Bar = 40µm F) qPCR analysis of MG63 and LM7 osteosarcomas treated with calcipotriol. Two-way ANOVA Multiple comparisons using Tukey’s test; *p* ≤ **0.01, ***0.001 (n=4) G) LM7 xenograft metastatic model in Nu/J (nude) mice. The images are from 5-week-old Nu/J mice that had LM7 cells delivered to their right flank. After xenograft, mice were immediately given either vehicle or calcipotriol (60 µg/kg b.w.) treatment until tissue harvest.

### Calcipotriol inhibits the spread and tumorigenicity of LM7 osteosarcoma *in vivo*

Given the pronounced effects of calcipotriol on LM7 gene expression and migration, an experimental LM7 xenograft mouse model was created to evaluate the efficacy of calcipotriol against metastasis and tumorigenicity (**Fig. 11A**). First, we ectopically engrafted LM7 cells into the right flank of athymic nude (*Foxn1^nu^*, also known as Nu/J) mice without matrigel to investigate metastatic potential of LM7 cells. Five weeks after LM7 transplantation, cancerous lesions in the lower mandibular/cervical neck region were observed in vehicle treated animals (**Fig. 11B, upper**). In contrast, animals given calcipotriol (60 µg/kg body weight three times per week) for five weeks showed no visible signs of cancerous lesions throughout their bodies (**Fig. 11B, lower**). Nine weeks after treatment, the animals were subjected to PET imaging as well as histological and anatomical analysis. Animals on vehicle developed large hematomas around the sub-mandible region after euthanasia, indicating an excess of broken blood vessels from localized tumors (**Fig. 11C,E**). Vehicle-treated mice developed numerous skin polyps around the sub-mandible region, while the dorsal skin displayed numerous cyst-like structures and wounds near the richly vascularized ears (**Fig. 11C**). Tumors also formed in the periprostatic adipose (pad) tissue of vehicle-treated mice, indicating a favorable niche (**Fig. 11C**). Histological examination of skin tissue of vehicle-treated mice confirmed scab formation and hyper dermal and epidermal responses (**Fig. 11D**), while lung tissues exhibited nodules in the outer respiratory tracks within the interlobular septum (**Fig. 11E**), indicating the presence more favorable niches. PET analysis confirmed calcipotriol’s anti-metastatic and anti-tumorigenic effects (**Fig. 11F**). Vehicle-treated animals had excessive ^18^F-flurodeoxyglucose (^18^F-FDG) uptake and distribution in the sub-mandibular, brachial plexus, and urogenital regions compared to calcipotriol-treated animals, which were nearly tumor-free (**Fig. 11F right, Movies S1 & S2**). Comparisons of maximal image projections (MIPs) revealed a statistically significant decrease in tumor size and spread after calcipotriol treatment (**Fig. 11G**). Finally, we used a CD44 human-specific antibody to track LM7 cells in order to better understand the cutaneous pathologies associated with LM7 xenografts (**Fig. S9**). Immunofluorescence analysis revealed LM7 localization at the polyp’s periphery but not within the centroid, implying paracrine effects of exogenous LM7 cells in polyp transformation (**Fig. 11H upper**). Furthermore, we observed both peri and intravascular localization of CD44-positive LM7 cells in vehicle-treated animals (**Fig. 11H lower**), but not in calcipotriol-treated animals (**Fig. S9**), indicating that endothelial transmigration was suppressed by calcipotriol. Finally, cyst formation in vehicle-treated animals appeared to be induced by paracrine effects of surrounding CD44-positive LM7 cells, as these cells were not integrated within the cysts themselves (**Fig. 11I**). Overall, calcipotriol significantly suppressed LM7 metastasis and tumorigenicity in humanized mice.

**Figure 11.**
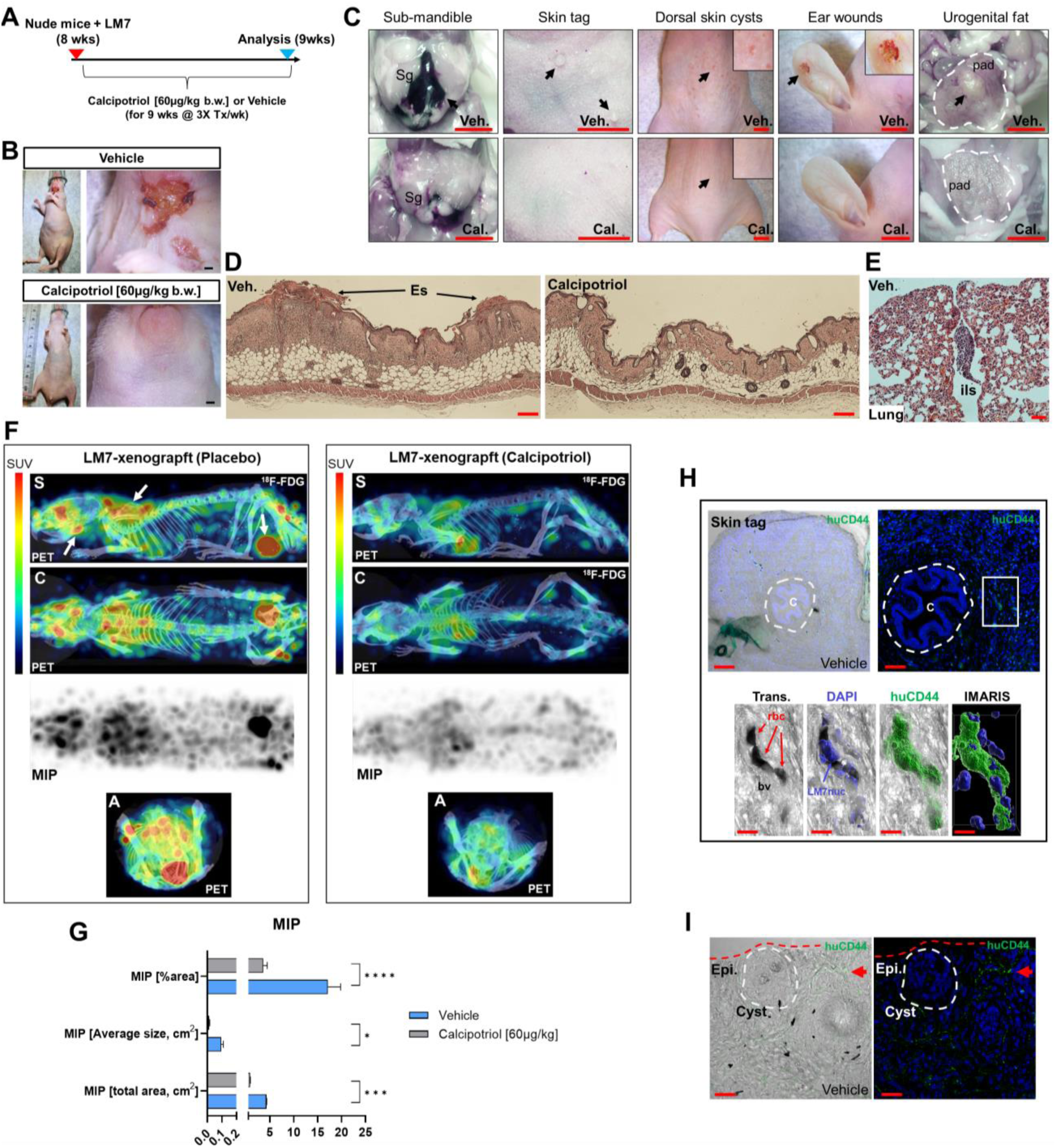
Calcipotriol suppresses LM7 spread and tumor growth in a xenograft mouse model. A) Overview of LM7-bearing xenograft calcipotriol treatment strategy. B) Sub-mandible lesions apparent by 5-weeks after LM7 transplantation only in vehicle-treated animals. Bar = 5mm. C) Gross anatomical and necroscopic assessment of vehicle and calcipotriol-treated animals after 9 weeks of treatment. Leftmost panel: Arrow depicts large hematoma. Submandibular gland (Sg). Rightmost panel: Outlined is the periprostatic adipose (pad) tissue with an arrow depicting tumor mass. All bars = 5mm D) Hyperdermal and -epidermal responses to LM7 cells. Hematoxylin and eosin staining of dorsal skin sections. Eschar (Es) Bar = 20µm E) H&E staining of outer lung of vehicle-treated LM7-bearing nude animal. Interlobular septum (ils) harbors a tumor nodule. Bar = 50µm F) PET imaging of treated LM7-bearing nude mice. Left arrows depict tumors localized to the sub-mandible, brachia plexus, and urogenital regions. Standardized tracer uptake values (SUV) are depicted in the color bar. Maximal intensity projection (MIP). Different views are represented as sagittal (S), coronal (C) and axial (A) image sections. G) MIP statistics. Unpaired T test; *p* ≤ *0.05, ***0.001, ****0.0001 (n=4) H) Immunofluorescence analysis of vehicle-treated skin tags using a human-specific CD44 antibody to detect LM7 cells. Center (C) of the skin epithelial polyp is encircled. The boxed region is highlighted below showing the localization of LM7 cells transmigrating within blood vessels (bv). Red blood cells (rbc). Bar (upper left) = 100µm, Bar (upper right) = 50µm, Bar (lower panel) = 10µm I) Immunofluorescence analysis of vehicle-treated skin using the huCD44 antsibody. The cyst is encircled and a cluster of LM7 cells (red arrow) is depicted in the periphery. Bar = 20µm

### Discussion

Despite recent advances in the field, OS remains a fatal disease. Because approximately 80% of patients are thought to have metastases at the time of initial presentation, understanding the cellular and molecular mechanisms that control OS-specific tumorigenicity and metastasis is critical for developing novel therapies^98^. Several studies have linked low vitamin D_3_ status to an increased risk of common cancers^25, 26, 33–38^. In contrast, the largest clinical trial of vitamin D_3_ supplementation ever conducted, and its *post-hoc* re-analysis, suggest that vitamin D_3_ benefits patients with advanced or lethal cancers associated with metastasis^25^. Despite these findings, the molecular and cellular mechanisms by which vitamin D_3_ influences cancer outcomes remain elusive. The current study sheds light on how 1,25(OH)_2_D and calcipotriol can influence tumor dormancy and outgrowth by regulating OS self-renewal and metastatic initiation potential (**Fig. 12**). In part, we show that 1,25(OH)_2_D inhibits OS self-renewal and EMT to block migration, while also suppressing the NMD machinery and re-expressing putative immunogenic antigens and pro-apoptotic proteins (e.g., BTN2A1, GILZ) from defined NMD-target genes. Furthermore, our findings reveal additional novel regulatory effects of 1,25(OH)_2_D, such as inhibition of desmoplasia and suppression of SOD2 non-canonical and neurogenic pathways, all of which have novel therapeutic implications. Depending on the metastatic state, 1,25(OH)_2_D also appeared to promote MET and will necessitate additional research to better understand the nature of the adhesion factors that halt OS migration *in vivo*.

**Figure 12.**
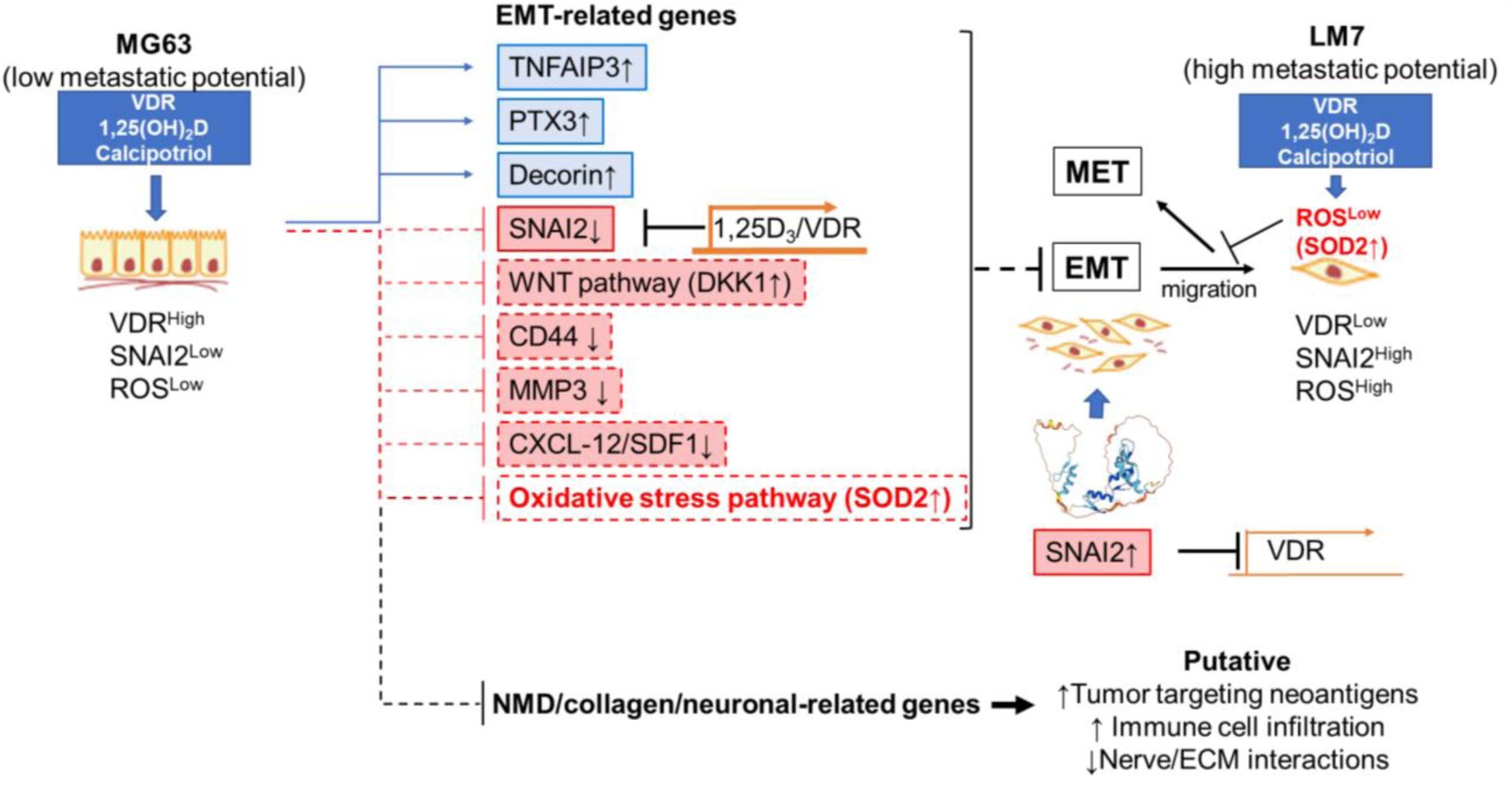
Novel anti-cancer properties of 1,25(OH)_2_D in osteosarcoma Schematic demonstrating the likely mechanism underlying the suppression of EMT, NMD, desmoplasia, and neural connections between OS by 1,25(OH)_2_D. The reciprocal regulation of VDR and SNAI2 constitutes one mechanism.

#### Vitamin D_3_ regulation of osteosarcoma EMT and desmoplasia

Several studies have shown that 1,25(OH)_2_D regulates EMT in multiple cancers (e.g., breast and colon)^93, 99–101^ and fibrosis-related tissue model systems^12^. For example, 1,25(OH)_2_D suppressed EMT by modulating a number of genes that encode EMT transcription factors (SNAI1/2, ZEB1, TWIST1), adherens junction proteins (E/N-cadherin), focal adhesion members (paxillin, integrins), tight junction components (claudins), cytoskeletal proteins (vimentin), ECM proteins (collagens, MMPs), and polarity proteins^100^. Despite these findings, the direct effects, and underlying mechanisms of 1,25(OH)_2_D in OS and cutaneous wound EMT have not been determined to date. Through chromatin interactions, we identified *SNAI2* as a direct VDR target gene, highlighting a central EMT inductive node for regulating OS metastasis. Other EMT transducers, such as ZEB1, ZEB2, E47, or TWIST1, were not influenced by 1,25(OH)_2_D^93^, implying disease-specific responses. Importantly, our findings have wider implications against other cancers with aberrant upstream receptor activation, as SNAI2 can work together with oncogenes such as RAS and ERBB2 to enhance tumorigenicity^102^. Our SNAI2 DsiRNA KD studies indicate that it may also repress *VDR* expression, most notably in the highly metastatic LM7 OS cell line. This finding is consistent with previous research in colon cancer cells, which found that SNAI2 repressed *VDR* expression, inhibiting the induction of E-cadherin to promote metastasis^93^. This newly discovered OS relationship points to decreased *VDR* expression in metastatic OS progression, which explains the reduced responsiveness to 1,25(OH)_2_D. However, an interesting finding was that the calcipotriol analogue enhanced anti-migratory effects in LM7 cells in both culture and in mice, suggesting clinical usage to overcome resistance in the future. Our analysis of the Cistrome DB Toolkit also suggests that SNAI2 may repress *VDR*, as observed in motile skin keratinocytes, which may be a common feature. Thus, combined approaches that target both the SNAI2 and the VDR signaling systems may be more effective in treating high grade malignant cancers and possibly skin disorders involving fibrosis such as scleroderma.

EMT entails the cellular deposition and remodeling of ECM, as well as the loss of the epithelial cell-to-basement membrane interactions, to generate a fibroblastic phenotype. Both 1,25(OH)_2_D and calcipotriol decreased collagen proteins and mRNA expression in OS spheroids and culture, a critical feature of EMT and tumor cell immunorecognition. These findings are consistent with recent observations in mammary epithelial cells^103^, and in different intestinal fibrosis mouse models, where both vitamin D_3_ dietary intervention and epithelial-specific *Vdr* ablation promoted fibrosis and pro-fibrotic factors^12^. Recent research indicates that hair follicle bulge stem cells can give rise to progenitor cells with fibroblast-like characteristics via EMT^104, 105^. Our findings show that ablation of the *Vdr* within hair follicle bugle stem cells results in increased migration and putative fibrotic responses within thickened neo-epidermal tissue after injury. Overall, the findings of this study suggest that modulating VDR signaling during key stages of cancer and tissue injury may be important from a therapeutic perspective.

#### Vitamin D_3_ regulation of the NMD pathway

Vitamin D_3_ modulation of the NMD pathway has yet to be reported. We found that vitamin D_3_ disproportionally induced the upregulation of NMD-target genes in OS cells, with the potential to sensitize immunotherapy in patients with OS. One unanswered question is how vitamin D_3_ regulates NMD machinery factors. According to recent research, upstream regulators of NMD include proteins involved in the integrated stress response (ISR) and the unfolded protein response (UPR)^106^. That is, in the presence of cellular stress, ISR and UPR are activated, resulting in overall suppression of protein synthesis and NMD factors. Indeed, our previous research has shown that 1,25(OH)_2_D treatment of MG63 induces mitohormesis, which involves UPR activation and a decrease in protein translation^107^, which may be linked to NMD inhibition as part of an adaptation response to restore mRNA integrity and full-length protein in the presence of PTC. However, it is unclear how vitamin D_3_ can induce selective suppression of both upstream and downstream components of the NMD pathway, regardless of ISR/UPR, which may entail, direct or indirect, gene-targeting elements induced by the VDR.

Most vitamin D-related immune research focuses on how it boosts the innate immune system^40, 47, 108–110^, but the biological/therapeutic effects of vitamin D’s cell-autonomous immunomodulatory responses in tumor cells are poorly understood. Recent research has demonstrated that combining vitamin D_3_ with immunotherapy can have sensitizing effects^111, 112^. In patients who were resistant to anti-PD1 alone, vitamin D_3_ supplementation in conjunction with anti-PD1 sensitized oral tumors and activated T cells^113, 114^. Also, Rituximab and Trastuzumab can detect surface antigens on B-cell lymphomas and breast tumor cells, but vitamin D deficient patients do not respond well to these therapies^115, 116^. Although the mechanism of action is unknown, our research indicates that vitamin D_3_ may make tumors more immunogenic by regulating novel effector systems such as NMD to sensitize tumor cell killing^17, 86^. Cancer’s NMD pathway is complex, with the ability to elicit both pro-tumor and tumor suppressor functions corresponding to the tumor’s genetic landscape and microenvironment^20^. On the one hand, NMD can be advantageous for cancer cells by inhibiting the expression of antigens with immunogenic activity. In contrast, targeted suppression of NMD in cancer cells has been used in preclinical studies to promote the production of neoantigens that induce an anti-tumor immune response^21, 117, 118^. Importantly, NMD is particularly pronounced in osteosarcomas, and the absence of adequate cell-to-cell adhesions (e.g., *CLDN1*, *CLDN15*) and a potent host immune response, makes vitamin D_3_ a likely candidate for combined immunotherapy^14–17^. Future research will seek to identify and validate novel functional neoantigens that may aid in the killing of OS tumor cells *in vivo*. In addition, it appears that calcipotriol may function similarly to 1,25(OH)_2_D, yet with enhanced effects on cell migration and gene expression across multiple pathways, such as EMT and NMD, depending on the subtype of OS cells^119^.

#### Vitamin D_3_ regulation of neuronal markers of osteosarcoma

Previous genome-wide association studies identified two susceptibility loci for osteosarcoma, one of which that achieved genome-wide significance was a single nucleotide intronic variant, rs1906953, at 6p21.3, in the glutamate receptor metabotropic 4 (*GRM4*) gene^19^. Although not formally and biochemically confirmed, it is believed that rs1906953, which is located at a DNase I hypersensitive site, enhances transcription of *GRM4* given that it is overexpressed in the majority of osteosarcomas^120^. Furthermore, the direct role of glutamate receptors in OS was recently investigated, whereby pharmacological treatment with the Riluzole glutamate receptor antagonist inhibited cancer cell proliferation and migration^121^. Despite these studies, how neurotransmitter receptors are regulated in osteosarcomas is unclear. Interestingly, an increasing body of evidence suggests that tumors, including gastrointestinal tumors, interact with neurons that innervate the tumor microenvironment, resulting in tumor progression and metastasis^122^. Importantly, our epigenetic and transcriptomic analysis indicates that OS may be classified as neuroendocrine tumors or carcinoids. Along these lines, a clinicopathological correlation was established in gastric cancer patients for overall survival in prespecified subgroups based on the level of innervation, which revealed higher levels of VDR expression in aneural samples and higher VDR levels correlated with lower grade gastric cancers^123^. Overall, our studies provide further evidence that impaired ionotropic glutamate receptors are part of the pathogenesis of osteosarcomas that can be regulated by vitamin D for potential clinical benefits.

#### Vitamin D_3_ modulation of intracellular iron and SOD2 functional localization in osteosarcoma

SOD2 is a mitochondrial superoxide dismutase that catalyzes the conversion of superoxide radicals (O_2_^−^) into molecular oxygen, which is a byproduct of oxygen metabolism that causes cell damage. Except for *SOD2*, most of the 1,25(OH)_2_D-modulated EMT pathway genes were not affected in highly metastatic LM7 OS cells. That is, it appears that reducing ROS is a common mechanism for facilitating 1,25(OH)_2_D-induced EMT suppression of migration. In advanced breast cancer cells, it also appears that ROS is able to activate SNAIL by recruiting NF-kB subunits to promoter regions to drive *SNAIL* expression^94^, thereby extending the significance of ROS and its regulation of EMT^124^. Our findings also indicate that the VDR regulates *SNAIL2* expression directly, in conjunction with its effects on ROS via *SOD2* upregulation, thereby providing a mechanism how SNAIL transcription factors can regulate ROS production directly^94^. Not only did 1,25(OH)_2_D increase *SOD2* levels, but it also promoted SOD2 mitochondrial localization to potentially act on mitochondrial ROS. In support of this, our previous research had shown that 1,25(OH)_2_D mediates antioxidant functions in OS cells by specifically lowering mitochondrial O_2_^−^ levels^107^. Elsewhere, breast cancer cells exhibit nuclear SOD2 as well as chromatin decondensation at genes involved in self renewal, dedifferentiation, and stemness reprogramming, resulting in increased metastatic potential^125^. Interestingly, new research has also revealed that SOD2 incorporated with iron (FeSOD2) functions as a nuclear pro-oxidant peroxidase, increasing oxidative stress^126^. Excess cellular iron has been linked to cancer progression, with increased cellular iron leading to metabolic misadaptations to oxidative stress^127^. Interestingly, FeSOD2 uses H_2_O_2_ as a substrate to promote tumorigenic and metastatic cancer cell phenotypes, whereas nuclear FeSOD2 promotes the induction of genes associated with EMT and stemness reprogramming^126^. In this context, we discovered that 1,25(OH)_2_D can suppress H_2_O_2_ across OS lines, as well as EMT and stemness/self-renewal genes, potentially compromising FeSOD2 nuclear pro-oxidant activity after treatment.

#### Novel vitamin D_3_-EMT interactions and model systems to establish and study in the future

We discovered that assessing several human OS lines for metastasis potential was more informative for their clinical relevance in epigenetic regulation studies of EMT and metastasis. Indeed, re-analysis of cDNA array data of metastatic lung tumors from primary osteosarcomas show that they in fact express higher levels of E cadherin compared to normal bone tissues and even primary OS samples themselves (**Fig. S10 & S11**)^128, 129^. This suggests that “metastatic” tumors re-expressed epithelial adhesion markers, and represent post-migratory tumors within newly established tumor microenvironments. Nevertheless, despite these limitations, based on the reanalysis of cDNA array data, SOD2 levels were consistently enhanced in metastatic OS samples, yet decreased relative to normal bone tissue, once again suggesting that SOD2 plays an upstream adaptive role in osteosarcomas and stimulation of its anti-oxidative organelle-specific functions may be key to therapy (**Fig. S10 & S11**).

Lastly, while this study focused on several common EMT factors, we did discover other novel 1,25(OH)_2_D interactions worth mentioning. For example, 1,25(OH)_2_D treatment increased the expression of pentraxin 3 (*PTX3*), a known pattern recognition molecule and oncosuppressor that inhibits FGF-dependent tumor growth and metastasis (**Fig. 1 & 12**)^130^. Furthermore, we discovered that 1,25(OH)_2_D treatment increased the expression of tumor necrosis factor alpha-induced protein 3 (*TNFAIP3*), which is required for the ubiquitination and degradation of EMT transcription factors like SNAIL1/2 and ZEB1 in gastric cancers^131^. Surprisingly, 1,25(OH)_2_D also inhibited C-X-C motif chemokine Ligand 12 (*CXCL12*) expression, which is a key pre-metastatic niche factor that recruits tumor cells (i.e., oncogenic “seeds”), thus escalating tumor progression and metastatic potential. Our findings also revealed that *SOX2* was enriched in highly metastatic LM7 cells, and that vitamin D_3_ can suppress its expression (GSE220948). EMT is often activated during cancer invasion and metastasis, generating cells with properties of stem cells^132^. Although *SOX2* is a well-known marker of (neuronal) stem cell pluripotency, recent research has shown that it also promotes cell migration and invasion in ovarian^133^, breast^134^, retinal^135^, and laryngeal^136^ cancers; thus, vitamin D_3_ in this context may have a broader impact. To date, it is unknown whether vitamin D_3_ can inhibit OS in patients or PDX animal models; therefore, future research is needed to create murine models of metastatic OS to provide *in vivo* tools to investigate vitamin D_3_ and OS more thoroughly.

## Author contributions

T.S.L. conceived, designed, and performed the experiments, analyzed the data, and wrote the manuscript. E.C. and V.M. performed the experiments. V.M. provided excellent technical support of the project. J.H. and V.M. edited the manuscript. E.C., G.S. and S.R. analyzed the data and edited the manuscript. All authors gave final approval to the manuscript.

## Acknowledgment

T.S.L. was supported by Grant # IRG-17-183-16 from the American Cancer Society, and from the Sylvester Comprehensive Cancer Center at the Miller School of Medicine, University of Miami. S.R. was funded by grant # 1R01CA215973. E.C. was supported by the grant NSF 19-500, DMS 1918925/1922843, JAX Computational Sciences, JAX Cancer Center (JAXCC) and NCI CCSG (P30CA034196). We would like to thank Ava King (U. Miami) for technical support, and Dr. Jonathan Trent from the Sarcoma Research Group at the Sylvester Comprehensive Cancer Center (SCCC) for providing the LM7 cell line. We would also like to thank Dr. Daniel Bilbao and Dr. Christian Mason (Cancer Modeling Shared Resource/SCCC) for excellent technical support and advice for the PET imaging studies.

## Data availability statement

All data generated during and/or analyzed during the current study are available from the corresponding author on reasonable request. RNAseq data: Gene Expression Omnibus GSE220948 (https://www.ncbi.nlm.nih.gov/geo/query/acc.cgi?acc=GSE220948)

## Materials and methods

### Reagents and human osteosarcoma cell lines

Crystalline 1,25(OH)_2_D (679101; MilliporeSigma, Burlington, MA, USA) was reconstituted in ethanol and kept at −80°C. Calcipotriol (hydrate) was reconstituted in ethanol before treatment (10009599; Cayman Chemical). Human MG63 osteosarcoma (CRL-1427; American Type Culture Collection, Manassas, VA, USA) and SaOS-LM7 (RRID:CVCL_0515; kind gift from Dr. Johnathan Trent, U. Miami) cells were cultured in complete media containing Eagle’s minimum essential medium (ATCC, 30–2003), 10% heat-inactivated fetal bovine serum (Gibco, Thermo Fisher Scientific, Waltham, MA, USA), and 100 U/mL penicillin, 100 mg/mL streptomycin (Life Technologies, Carlsbad, CA, USA). For assays, cells were treated with 0 (vehicle; equal-volume ethanol; 0.0001%), 10nM, and 100nM 1,25(OH)_2_D incubated in tissue culture plates (CytoOne, USA Scientific, Ocala, FL, USA) at 37°C in a humidified atmosphere of 5% CO_2_, 95% air.

### Lineage tracing and migration of wound induced K15 bulge stem cell progenitors in the absence of Vdr

The Tg(Krt1-15-cre/PGR)22Cot line (referred to as K15-CrePR1; JAX stock no. 005249) was on a mixed C57BL/6 and SJL background and was utilized to generate RU486-inducible, Cre-mediated gene deletions in BSCs of mouse hair follicles. The Gt(ROSA)26Sortm1(CAG-Brainbow2.1)Cle strain (also known as R26R-Confetti; JAX stock number 013731) was maintained on the C57BL/6J strain. Professor Geert Carmeliet (KU Leuven) kindly provided mice with a floxed *Vdr* gene that were generated as previously described ^137^. Prior to conducting tests, the conditional alleles and deletion efficiencies were validated via PCR analysis. Floxed *Vdr* mice were bred with *K15*-CrePR1 and R26R-Confetti mice expressing Cre-recombinase under the control of the K15 promoter in order to trace the lineage of subsequent progenitor cells following RU486 treatment and wounding. Each experimental mouse received a 3-mm-diameter, full-thickness truncal wound with a sterile biopsy punch (Sklar Instruments, PA). As an analgesic, slow-release buprenorphine was administered, and the animals were monitored daily for signs of distress. After sacrifice, the degree and cellular composition of re-epithelialization were evaluated by obtaining tissue in PBS containing 4% PFA (pH 7.4). Wounds were sectioned and imaged with a Zeiss LSM 880 confocal laser scanning microscope after being cut along the midlines and placed in cryofreezing liquid. Using serial wound sections for Confetti-labeled cell counting, 4-5 separate animal experiments were conducted.

### Scratch migration and Boyden chamber invasion assays

The scratch assay was used to examine cell migration following scratch-induced cell loss. Cells were grown to confluence, then glass Pasteur pipettes were used to scratch the surface of the vessels in a vertical direction. To prevent re-adhesion of dissociated cells, the wells were immediately rinsed in PBS. Cells were cultured under a variety of conditions, with the gap distance being monitored every day for up to 72 hours. After 72 hours, cells were fixed with 4% PFA and stained with a crystal violet solution containing 0.05% crystal violet in 30% methanol. In certain instances, staining with 4-HNE was performed after scratching. For impedance-based detection of migrating cells, Boyden chamber invasion tests were conducted using Millicell with hanging Transwell polyethylene terephthalate inserts (8 µm pore size, Millipore, PIEP12R48). 200 µl of serum-free medium containing 5104 unstimulated MG63 or LM7 cells was added to the upper compartment, while 750 µl of complete EMEM medium with or without 1,25(OH)_2_D, was added to the lower compartment. Cotton swabs were used to remove non-migratory cells from the upper chamber after 24 hours. On the lower side of the Transwell membrane, migratory cells were fixed with 4% PFA and stained with DAPI. Using ImageJ, cells were counted using a Zeiss 880 LSM confocal microscope (Olympus).

### ATACseq and analysis

ATACseq was used to evaluate the chromatin of MG63 cells (Active Motif, Carlsbad, CA, USA; 53150). At the Sylvester Comprehensive Cancer Center’s Oncogenomic Core Facility, DNAseq library preparation was performed. Samples were sequenced on an Illumina NovaSeq 6000 utilizing 100-bp paired ends. Using Cutadapt, sequencing reads (40 million per sample) were trimmed of Nextera adaptor sequences and filtered. Reads were mapped to the reference genome (hg38 Canonical) utilizing Bowtie2 with the following parameters: “extremely sensitive end-to-end (–very-sensitive)”, “set the maximum fragment length for viable paired-end alignments: 1000”, and “enable mate dovetailing to produce BAM files.” BAM data sets on a range of variables were used to filter out uninformative reads with poor mapping quality and improper pairing (Galaxy Version 2.4.1). ≥30 was applied to the read mapping quality (phred scale) filter. ATACseq TF motifs were called using HOMER (http://www.homer.ucsd.edu).

### RNA-seq and analysis

RNA-seq data has been submitted to GEO with accession GSE220948. Total RNA was isolated from MG63 lysates (*n* = 2 per treatment condition) using the PureLink RNA Mini kit (12183018A, Thermo Fisher Scientific) with DNase set (12185010, Thermo Fisher Scientific). The Agilent Bioanalyzer confirmed that the RIN counts of the RNA samples were greater than >8.5. The Oncogenomic Core Facility of the Sylvester Comprehensive Cancer Center was responsible for library preparation and RNA-sequencing (University of Miami). Samples were sequenced using 75 bp paired ends on an Illumina NextSeq 500 (San Diego, CA, USA), resulting in ∼30 million trimmed and filtered reads per sample using Cutadapt. Compressed Fastq.gz files were uploaded to a Galaxy account (https://usegalaxy.org) and concatenated data sets tail-to-head (Galaxy Version 0.1.0). HTseq-count was executed to generate gene counts that were not standardized for each sample. iDEP (http://bioinformatics.sdstate.edu/idep93) was utilized to perform normalization of RNA-seq gene counts (counts per million [CPM]) as well as differential gene expression and visualization analyses. Individual gene expression graphs produced from RNA-seq were subjected to a two-way ANOVA with Bonferroni’s multiple comparisons test using the CPM values. The *p* value summaries were represented as \*\*\*\**p* ≤ 0.0001, \*\*\**p* ≤ 0.001, \*\**p* ≤ 0.01 and \**p* ≤ 0.05. Using statistically significant differentially expressed genes, GSEA (v 4.3.2) was used to evaluate whether an a priori-defined collection of genes differed between the two biological states. GSEA used a normalized enrichment score (NES) > +/− 2, nominal *P* value (NOM *P*-Val) < 0.05 and false discovery rate (FDR) < 0.05 for statistical significance. The molecular Signatures Database v7.3 with signature gene sets was utilized for GSEA. The “clusterProfiler” R package was used to analyze GO and KEGG enrichment^138^. ShinyGO enrichment tool ^139^ was also utilized for data analysis and graphical representation.

### Analysis of Osteosarcoma patient samples using microarray data

Raw probe-level data (.CEL files) from Affymetrix HG U133A microarray experiments were collected from the Gene Expression Omnibus (GEO) and reanalyzed in order to evaluate human osteoblast patient samples. The series GSE16088^128^ included array data for 14 OS tissue samples, whereas the series GSE14359^129^ included array data for 10 OS tissue samples, four metastatic lung OS tissue samples, and a normal bone tissue sample. Normal osteoblast states were examined utilizing the GSE39262 series. The CEL files were read using the R package “oligo”^140^. Averages of the probe set of values were normalized using the robust multichip algorithm (RMA) method of “oligo” R package, as well as heatmap and PCA plots. The “limma” R package was used to do differential expression analysis^141^. Genes with adjusted *p*-values ≤ 0.05 and log2 foldchanges ≥2 were determined to be differentially expressed.

### Comprehensive analysis and processing of epigenomic data sets for osteosarcoma

We reanalyzed raw genome-wide histone modification data from series GSE74230, which includes ChIP-seq of H3K4me1, H3K27ac, and DNase I Hypersensitivity data in metastatic and non-metastatic OS cell line pairings, as well as primary and metastatic tumors in human patients. Bowtie2 was used to index the sequences to the reference hg38 genome^142^. BowtieQC (v 0.11.9) was used to assess the quality of the raw FASTQ files. Reads that were not uniquely mapped or redundant were removed. The “callpeak” module of Model-based Analysis for ChIP Sequencing (MACS v 2.1.3) was utilized to identify regions with ChIP-seq enrichment over the background (*q* < 0.01)^143^. The R Bioconductor program ChIPseeker^144^ was used to generate Brower Extensible Data (BED) format files depicting genomic areas with each histone modification and DNase I hypersensitive site peak for the annotation of enriched peaks acquired from the MACS2 callpeak function. The 500bp sequences surrounding the peak summits were recovered from the promoter region, and transcription factor motif finding was performed using HOMER version 4.11 and ranked by adjusted *p*. CISTROME-GO was implemented to evaluate gene-level regulatory potential (RP) scores and enrichment of Kyoto Encyclopedia of Genes and Genomes (KEGG) pathways between H3K27ac changes in MG63 and LM7 OS cells ^77^. A half-decay distance of 10.0 kb was established for all differential peaks found by DESeq2 (*P*_adjusted_ < 0.05, log2-fold change > 1). The epigenetic changes were also annotated independently using the EaSeq platform^145^. Peak annotations for H3K27ac (−500 and +1000 from TSS) and H3K4ME1 (−100 and +5000 from TSS) were determined using a log2 FC threshold for LM7 and MG63 samples. Overrepresented peaks were defined as log2 (0.2 to 11.6), while underrepresented peaks were defined as log2 (−0.1 to -2). The Cistrome DB toolkit was used to extract *SNAI2* and *VDR* TFs based on >250,000 epigenomic data sets taken from human and mouse samples [13]. EaSeq and Integrative Genome Browser (IGV) was used to visualize peaks, correlation plots (poised and active enhancer elements), and heatmaps.

### Target gene prediction for nonsense-mediated RNA decay and statistical comparison

On the basis of four studies in which NMD effectors were inhibited in human HeLa cells, a list of core NMD-target and non-NMD-target genes was compiled ^79–82^. NMD targets were filtered if they exhibited a ≥2-fold upregulation in response to UPF1/SMG6 knockdown and expressed FPKM ≥ 5 ^81^. Non-target genes of NMD were categorized as showing no or marginal upregulation (i.e., < 1.5-fold upregulation after knockdown^79–82^). Based on this analysis, more than >4,000 genes were identified as NMD nontargets, while more than >900 genes were identified as NMD targets by at least one study; however, 50 genes were identified as NMD targets by all studies, and we used these 50 genes as our core NMD-target gene set (**Fig. S4**). We compared our RNA-seq data from MG63 cells to NMD-target and nontarget genes. GraphPad was used to calculate descriptive statistics, such as skewness and kurtosis, for the NMD target and non-target genes. Using GraphPad, a ROC curve was generated by comparing the true positive rate to the false positive rate for NMD target and non-target genes. Multiple comparisons were conducted for non-NMD target genes by randomly selecting 39 genes from the pool of non-NMD target genes. GraphPad generated C statistics based on the area under the curve. Anti-phosphorylated UPF1 antibody (Millipore Sigma 07-1016) was used on MG63 cells.

### Disruption of the NMD SMG7-UPF1 complex to using NMDI-14

NMDI-14 inhibitor (Ethyl 2-(((6,7-dimethyl-3-oxo-1,2,3,4-tetrahydro-2-quinoxalinyl)acetyl)amino)-4,5-dimethyl-3-thiophenecarboxylate) was purchased from Calbiochem (Cat. Nr. 530838) and reconstituted in DMSO (10mg/ml; 24mM). For MTT assay, MG63 cells were treated with NMDI-14 at 0.05nM-50µM (10-fold series) with corresponding matching DMSO series performed independently in 96-welll plates. A same batch of MG63 cells were cultured in 6-well plates for total RNA collection and qPCR analysis. Cell toxicity and stress was observed at 50µM NMDI-14 at both 24 and 48 hours of treatment, which was reported not to occur with other cell lines^89^.

### DsiRNA SMG6 knockdown to evaluate NMD target gene expression

Dicer-substrate short interfering RNAs (DsiRNAs) and TriFECTa® kits (Integrated DNA Technologies) were utilized to knockdown *SMG6* more effectively than conventional siRNA. All RNA strands for DsiRNAs were synthesized as single-strand RNAs (ssRNAs) and resuspended in RNase-free water following purification by high-performance liquid chromatography. ssRNAs were annealed at 95°C for 5 minutes to form DsiRNA duplexes, then incubated at room temperature for 4 hours before being aliquoted and stored at 80°C. 27-mer hs.Ri.SMG6.13.3 duplex DsiRNA targeting the coding sequence of *SMG6* was sufficient for >60% knockdown (**Fig. S5**). For optimization, endogenous gene positive controls and qPCR assays (HPRT-S1 DsiRNAs and qPCR assays) were also employed. As a reference, a universal negative control (non-targeting DsiRNA) was utilized in all experiments as a reference. MG63 cells were transfected with BioT (Biolands), and after optimization, 20-60nM of targeting DsiRNAs were used. After 24 hours of treatment, the effects of *SMG6* knockdown on NMD-target and non-target genes were assessed using qPCR. The two-way ANOVA test with Tukey’s multiple comparisons was used to conduct statistical analysis. All data represent the mean standard deviation (S.D.) of at least three separate experiments. **** *p ≤* 0.0001.

### Analysis of soft agar osteosarcoma spheroids

MG63 and LM7 cells were seeded into 0.4% low-melting-point agarose (Lonza, Basel, Switzerland; 50101) on top of a 1% agarose layer (1000 cells per well, 24-well plate). For approximately 14 days, cells were maintained in an incubator at 37°C and 5% CO_2_ with vehicle or 1,25(OH)_2_D. The Live-or-Dye^TM^ Zombie Aqua fixable viability staining kit was used for live cell analysis (32004-T; Biotium). For quantification, colonies were fixed in 4% PFA and counted using a dissecting scope (Zeiss Stemi 508, Carl Zeiss, Jena, Germany) and ImageJ software. Images were converted to 8-bit, background subtracted using rolling ball radius of 1000 pixels and light background, and then applied threshold using Internodes correction. The spheroids were enumerated using the particle analyzer with the settings: size = 0.01/0.001-10mm^2^; circularity = 0-1.0. All experiments were designed with three replicates per condition. An ordinary one-way ANOVA with Sidak’s multiple comparisons test was conducted. A number of spheroids were extracted from soft agar and immunostained. Unconjugated rabbit polyclonal antibodies were used to detect 4-hydroxynonenal, type 1 and type 3 collagens (bs-6313R, bs-10423R, and bs-0948R; Bioss). Because of their small sizes, LM7 colonies were not extracted.

### DsiRNA SNAI2 knockdown to evaluate the effects of 1,25(OH)_2_D on gene expression and migration

Dicer-substrate short interfering RNAs (DsiRNAs) and TriFECTa® kits (Integrated DNA Technologies) were utilized to knockdown *SNAI2* more effectively than conventional siRNA. All RNA strands for DsiRNAs were synthesized as single-strand RNAs (ssRNAs) and resuspended in RNase-free water following purification by high-performance liquid chromatography. ssRNAs were annealed at 95°C for 5 minutes to form DsiRNA duplexes, then incubated at room temperature for 4 hours before being aliquoted and stored at 80°C. 27-mer hs.Ri.SNAI2.13.2, but not hs.Ri.SNAI2.13.1, duplex DsiRNA targeting the coding sequence of *SNAI2* was sufficient for knockdown (**Fig. S7**). Transfection efficiency control, TYE 563-labeled DsiRNAs, was used for optimization studies (**Fig. S6**). For optimization, endogenous gene positive controls and qPCR assays (HPRT-S1 DsiRNAs and qPCR assays) were also employed. As a reference, a universal negative control (non-targeting DsiRNA) was utilized. LM7 cells were transfected with BioT (Biolands), and after optimization, 80nM of targeting DsiRNAs were used. After 24 hours of treatment, the sensitizing effect of *SNAI2* DsiRNAs on 10nM 1,25(OH)_2_D-mediated gene expression in LM7 cells was determined using qPCR. The Student’s *t*-test was used to conduct statistical analysis. All data represent the mean standard deviation (S.D.) of at least three-four separate experiments. * *p ≤* 0.05, ** *p ≤* 0.01, *** *p ≤* 0.001, and **** *p ≤* 0.0001.; Student’s t-test.

### Western Blotting

MG63 cells were lysed in RIPA buffer (50mM Tris-Cl pH 7.4, 150mM NaCl, 0.1% SDS, 0.1% sodium azide, 0.5% Na-deoxycholate, 1mM EDTA, 1% NP-40, 1X Protease inhibitor cocktail) and centrifuged for 10 minutes at 13,000 g and 4 C. The amount of total protein was determined using a BCA kit. Protein samples (20 µg) were denatured at 95 C for 5 minutes in 1X Laemmle buffer (Tris-HCl pH 6.8, 2% SDS, 20% glycerol, 0.2% bromophenol blue, 0.025% beta-mercaptoethanol). Proteins were separated by SDS-PAGE, transferred to an activated polyvinylidene fluoride membrane, and then blocked for 1 hour at room temperature with 5% non-fat dried skim milk/1X TBST (Tris-buffer saline, 0.1% Tween 20). The primary antibodies were diluted in a buffer of 0.5% milk/1X TBST (CD44 antibody; 15675-1-AP, Proteintech). The Li-COR system was used to detect antibody binding.

### Chromatin Immunoprecipitation (ChIP)-qPCR

After 24 hours of treatment with vehicle or 10nM 1,25(OH)_2_D, MG63 cells (∼1.0 × 10^7^) were cross-linked with 1% formaldehyde at room temperature for 10 minutes. Cross-linking was inhibited with 100 mM glycine, and cells were lysed in 1 ml expansion buffer (25 mM HEPES pH 8.0, 1.5 mM MgCl2, 10 mM KCl, 0.1% NP-40, 1X PIC) and centrifuged at 2000 rpm to recover nuclei. Fixed nuclei were re-suspended in 1 ml sonication buffer (50 mM HEPES pH 8.0, 140 mM NaCl, 1 mM EDTA, 1% Triton X-100, 0.1% Na-deoxycholate and 0.1% SDS) containing a protease inhibitor cocktail and sonicated with a Q125 sonicator (QSonica) to generate fragment sizes of 200-800 bp. The supernatant was pre-cleared for 1 hour at 4°C with protein A Dynabeads (Invitrogen). Chromatin samples were incubated overnight at a low temperature with 10 µg ChIP-grade anti-RNA polymerase II (ab26721; Abcam, Boston, MA, USA) and anti-VDR (sc-1008x; Santa Cruz Biotechnology, Santa Cruz, CA, USA) antibodies. IP complexes were captured using protein-A Dynabeads (pre-blocked with 0.5 % BSA/1X PBS) and washed three times each with wash buffer-1 (50 mM HEPES pH 8.0, 500 mM NaCl, 1 mM EDTA, 1% Triton X-100, 0.1% Na-deoxycholate and 0.1% SDS), wash buffer-2 (20 mM Tris HCL pH 8.0, 1 mM EDTA, 0.5% NP-40, 250 mM LiCl, 0.5% Na-deoxycholate and 1X PIC) and TE buffer. Immunoprecipitated chromatin was eluted at 65°C for 10 minutes in 200 l of elution buffer (50 mM Tris pH 8.0, 1 mM EDTA, 1% SDS, 50 mM NaHCO3). Input and IP fractions were treated with 20 µg RNase A and 40 µg Proteinase K for 1 hour at 42 C and 65°C, respectively. Overnight, reverse crosslinking was performed at 65°C. DNA was extracted with phenol: chloroform: isoamyl alcohol (25:24:1), and ethanol precipitation was performed with 3M sodium acetate (pH 5.2) and 2 µg of glycogen. In 30 µl of nuclease-free water, DNA samples were resuspended after being washed with 70% ethanol. qPCR was conducted using the standard SYBR-green method.

### LM7 xenograft animal model and Positron Emission Tomography (PET), data acquisition and human cell tracking

Xenografts were performed on female homozygous Nu/J mice (strain #: 002019) aged 8-9 weeks and weighing 24-26 grams that were procured from The Jackson Laboratory (Bar Harbor, ME, USA) and maintained in the animal care facility of the Neuroscience Annex Building at the University of Miami (Miami, FL, USA). The Institutional Animal Care and Use Committee of the University of Miami Miller School of Medicine approved all animal procedures (protocol 21-107). All animals were kept in accordance with the circadian cycle (12hr light/12hr dark cycle) and were provided with sterilized food and drink *ad libitum*. 1X10^6^ LM7 cells/100 µl dilutant (1% ethanol in PBS cat#: 21-031-CV, Corning, Corning) were injected subcutaneously into the right flanks of Nu/J mice without matrigel. The animals were subsequently treated with either PBS (ethanol vehicle) or calcipotriol (60 μg/kg b.w., catalog number: NC1022293, Fisher Scientific; reconstituted in ethanol) three times each week for nine weeks. This dose is known not to induce hypercalcemia or any deleterious side effects^146^. Mice were humanely euthanized, and tissues harvest for qPCR analysis of human OS markers against mouse beta actin mRNA expression. Prior to sacrifice, PET imaging studies were conducted at the Cancer Modeling Shared Resource part of the Sylvester Comprehensive Cancer Center. A continuous body weight loss of >20% was also a predetermined survival endpoint, which did not occur during the 9 weeks of treatment. During imaging, experimental animals were anesthetized using isoflurane, and the isoflurane inhalation will be maintained throughout scanning. Animals were fasted overnight prior to ^18^F-fluorodeoxyglucose (FDG) positron-emitting radiotracer administration and start of imaging to monitor tumor response. A maximum of 10 µl per gram of body weight of ^18^F-FDG was injected intravenously via the tail vein per mouse (24 MBq of ^18^F-FDG, for 60 min). The whole-body PET scanning was done on a MILabs VECTor instrument with animals maintained under anesthesia and normothermia while monitoring vital signs in dorsal recumbency on a heating pad. For tracking human LM7 cells in nude mice, we utilized the anti-CD44 mouse monoclonal antibody (CF488A) [clone: HCAM/918] (Biotium, BNC880918-100) with affinipure Fab fragments goat anti-mouse IgG (H+L) (Jackson ImmunoResearch Laboratories, 115-007) blocking (**Fig. S9**).

### MTT Assay

All experiments were performed using 2.4 × 10^4^ cells/well in 96-well plates. Cells were seeded and then left to incubate at 37°C (5%CO2) overnight. The following day, cells were carefully washed with 1X PBS, and then treated with 1,25(OH)_2_D and the cellular status was assessed at 48 hours post treatment. The MTT (3-[4,5-dimethylthiazol-2-yl]-2,5 diphenyl tetrazolium bromide) assay was performed according to the manufactureŕs recommendation (10009365; Cayman Chemical). After incubation with the MTT reagent, the cells and salt were solubilized using the provided sodium dodecyl sulfate-based lysis buffer. The optical density absorbance was determined using a Molecular Devices EMax microplate spectrophotometer at 550 nm absorbance. Data is represented as the mean of six replicate wells ± SD, with analysis using the two-way ANOVA with Bonferroni post-hoc test (GraphPad Prism).

### Image processing

Imaris was used to process and analyze Zeiss LSM 880 LS microscope-generated confocal images (Bitplane). Imaris (Bitplane) was used to perform 3D reconstructions using the semi-automated Surfaces feature. Using the multiple-channel masking tool, colocalization analysis was conducted. Using regions of interest on maximally projected images, the average mean fluorescence intensities were obtained. Adobe Illustrator and Microsoft PowerPoint were utilized to develop the diagrams.

### Data availability

All data that support the findings of this study have been included in this manuscript.

## Supplemental Figures

**Figure S1.**
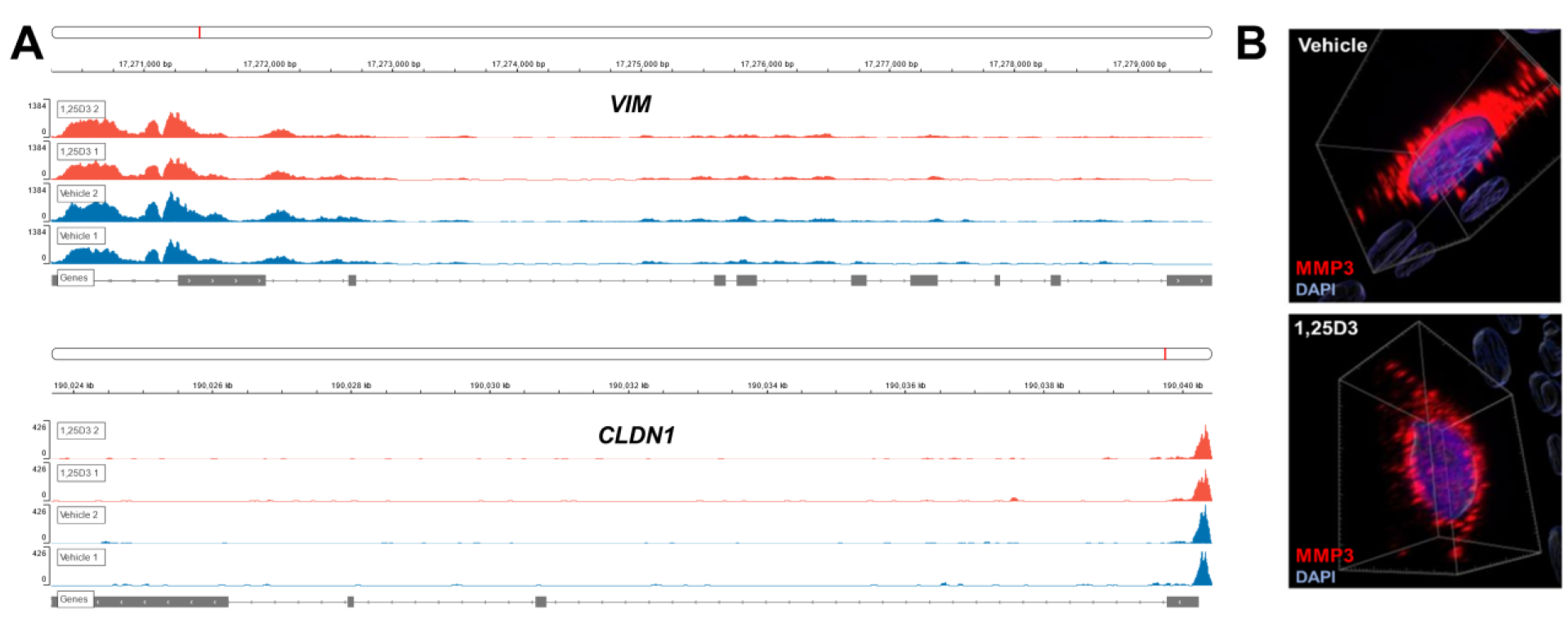
MMP13 expression and ATAC-seq genome browser tracks from *VIM* and *CLDN1*. A) 24 hours of 1,25(OH)_2_D treatment of MG63 cells at 10nM concentration. B) MG63 and MMP3 expression. Visualized using IMARIS.

**Figure S2.**
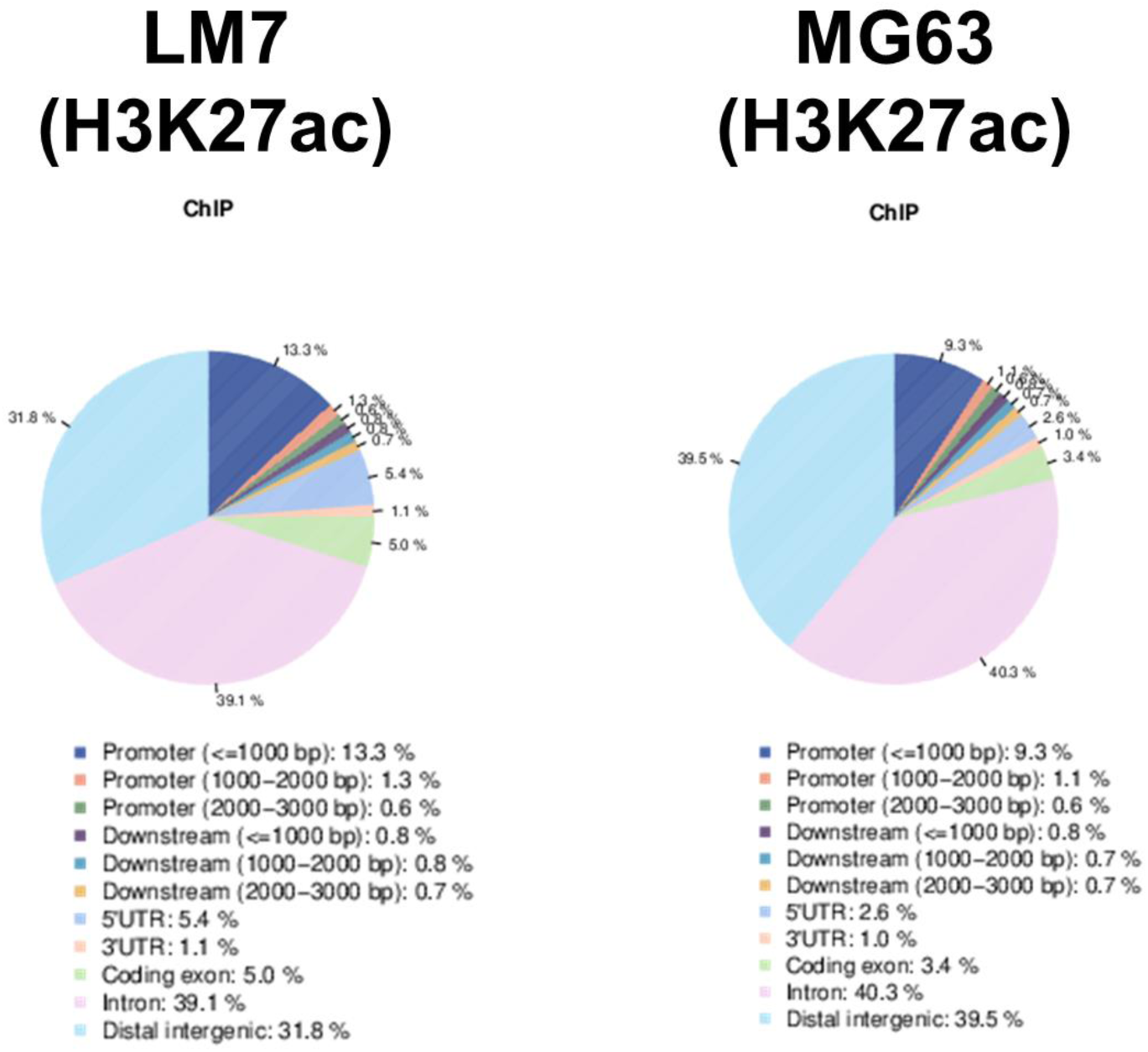
Pie chart distribution of ChIP regions (H3K27ac) over chromosomes for LM7 and MG63 osteosarcoma lines. The percentages indicate the total ChIP, with the proportion of ChIP regions indicating the enrichment and significance of genomic structures such as promoters, gene bodies, etc.

**Figure S3.**
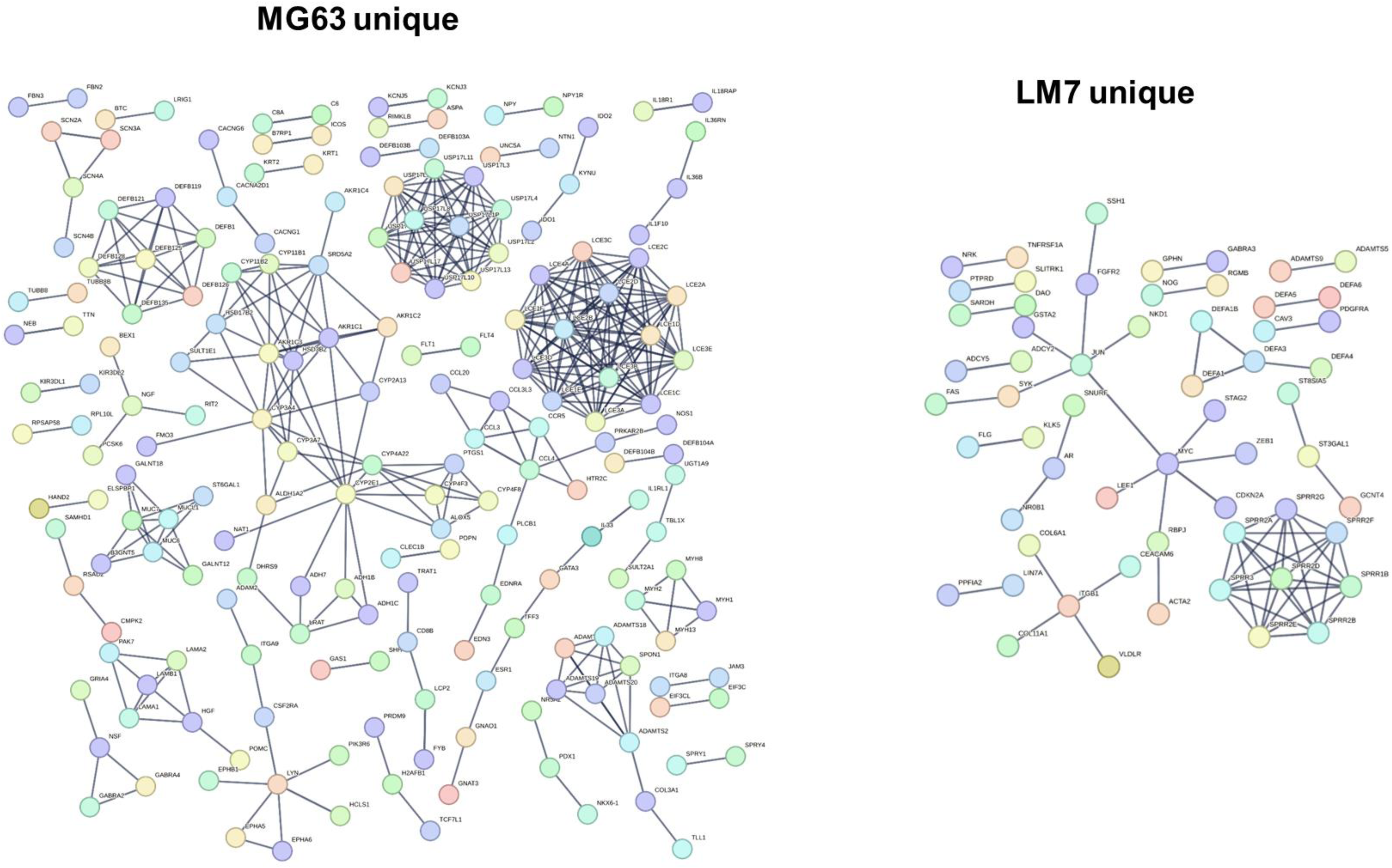
Network associate of putative target genes from H3K27ac peaks in MG63 and LM7 osteosarcoma cells. The network corresponds to Figures 4E and F in the main text. Additional functional enrichment analysis of these networks is provided as Supplementary Material.

**Figure S4.**
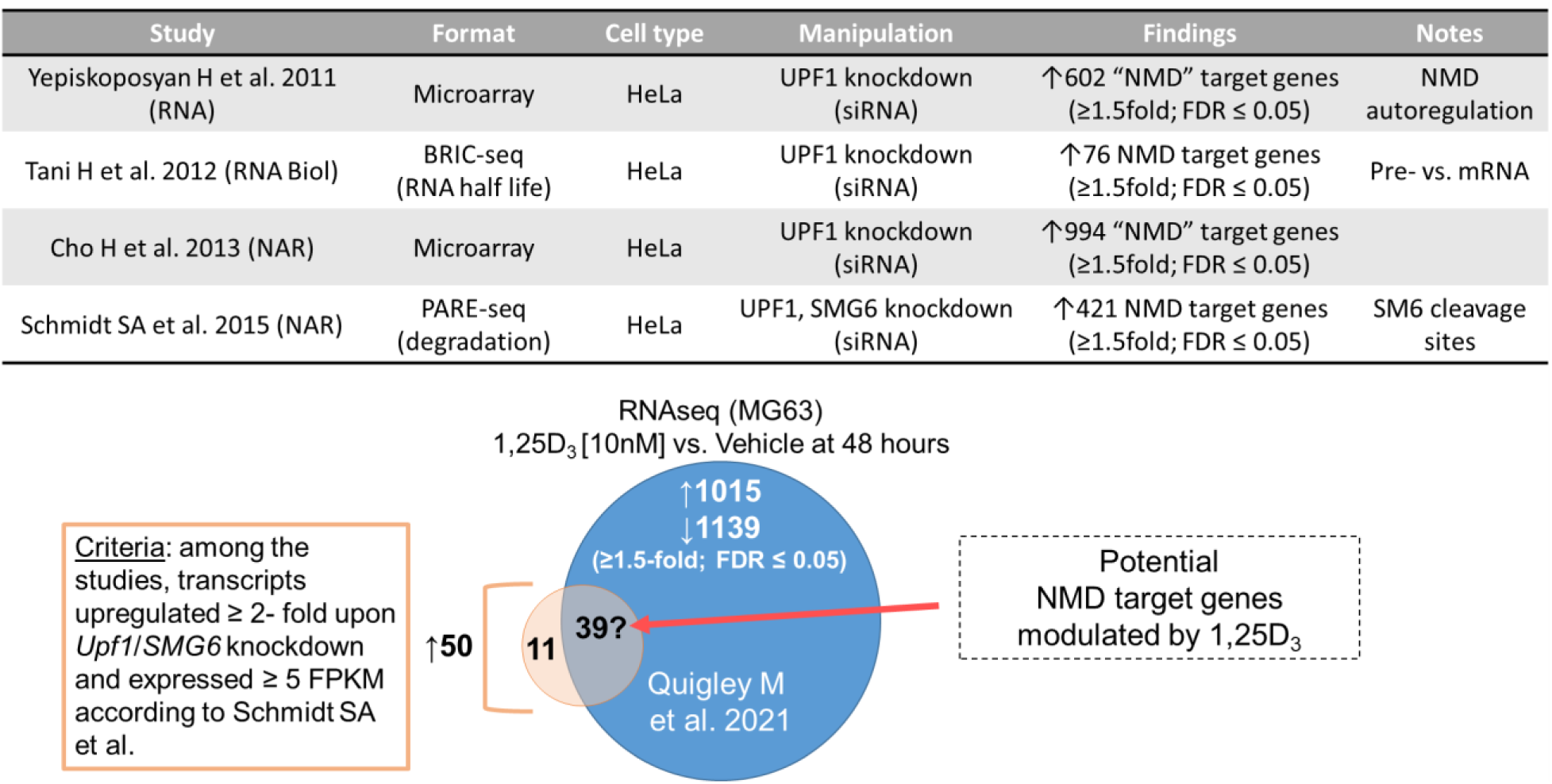
Potential NMD target genes modulated by 1,25(OH)_2_D in MG63 cells. Four major studies that manipulated the NMD machinery were screened for common NMD target and non-target genes. Those genes were then compared to differentially regulated gene sets modulated by 1,25(OH)_2_D. The overlapping gene set expression pattern was analyzed in the main text.

**Figure S5.**
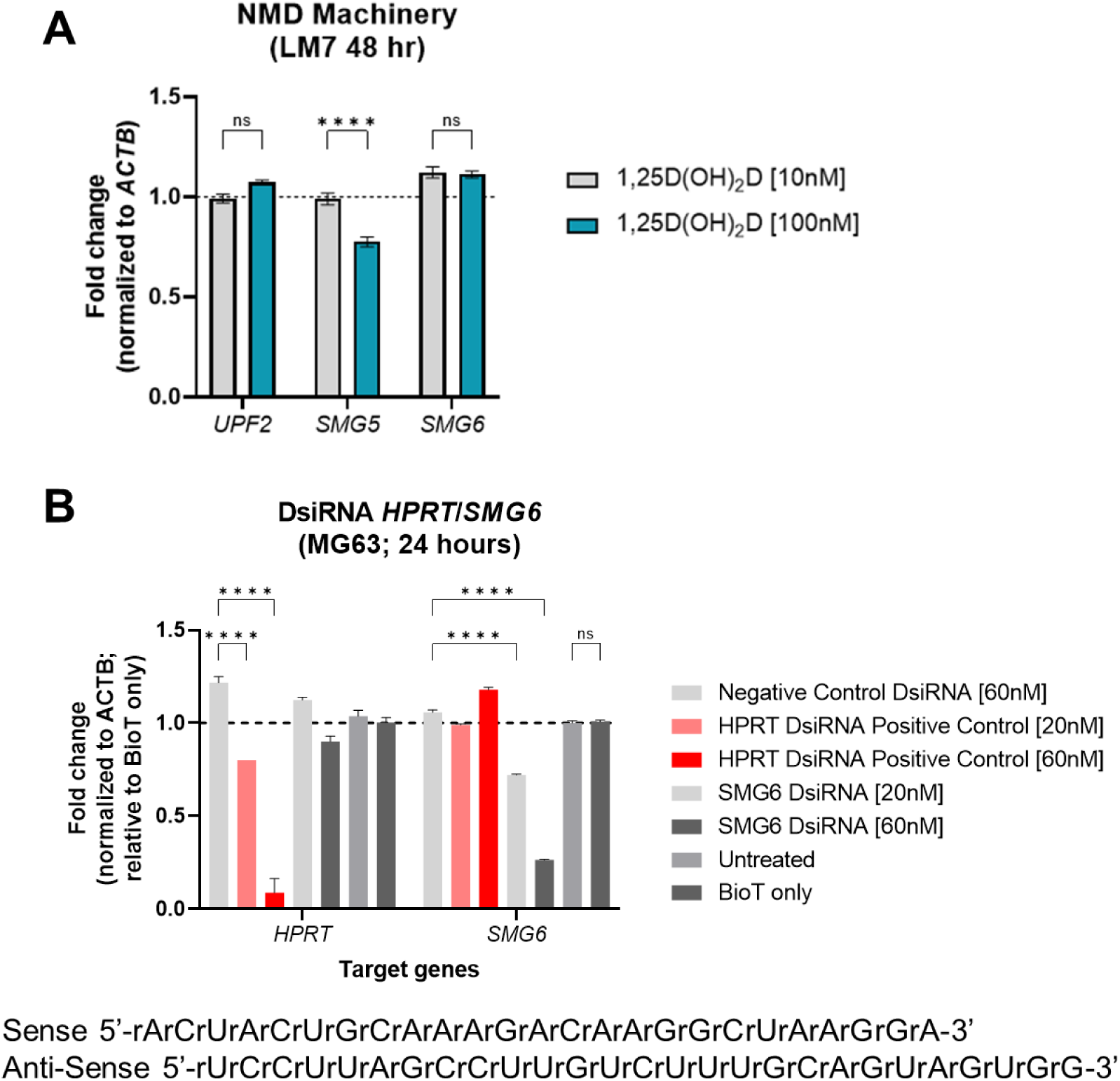
DsiRNA knockdown of control *HPRT* and *SMG6* in MG63 cells. A) Partial suppression by 1,25(OH)_2_D of the NMD machinery genes in LM7 cells. LM7 cells treated for 48 hours with 100nM 1,25(OH)_2_D exhibit a downregulation of SMG5. Two-way ANOVA Tukey’s test for multiple comparisons; *p* ≤ ****0.0001 (n=3). B) Specific knockdown of *SMG6* using the SMG6.13.3 duplex. Duplex sequences shown below the graph. Two-way ANOVA test with Tukey’s multiple comparisons. All data represent the mean standard deviation (S.D.) of at least three separate experiments. **** *p* < 0.0001.

**Figure S6.**
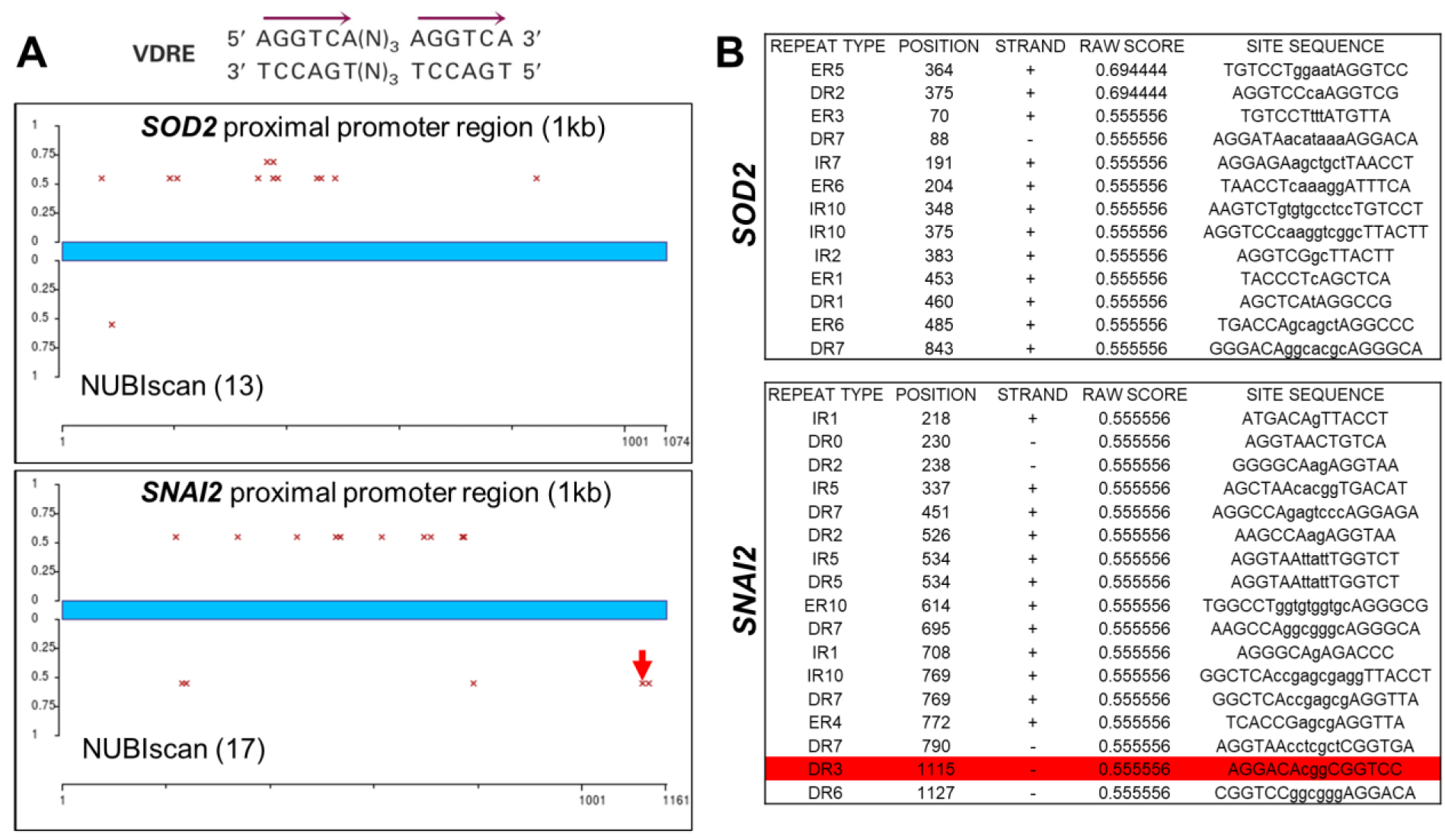
VDR directly enhances the transcriptional activity of *SNAI2* but not *SOD2*. A) Using NUBIScan (http://nubiscan.unibas.ch), potential vitamin D receptor response elements (VDREs) in the *SOD2* and *SNAI2* promoter regions were anticipated. As shown by the blue bar, the starting point is the transcription start sites (TSS, +1000bp). B) *SOD2* has 13 nuclear receptor binding sites, while *SNAI2* had 17. *SNAI2* was the only gene that featured a putative VDRE with a direct repeat 3 (DR3) intervals for RXR binding (highlighted by the red arrow and bar).

**Figure S7.**
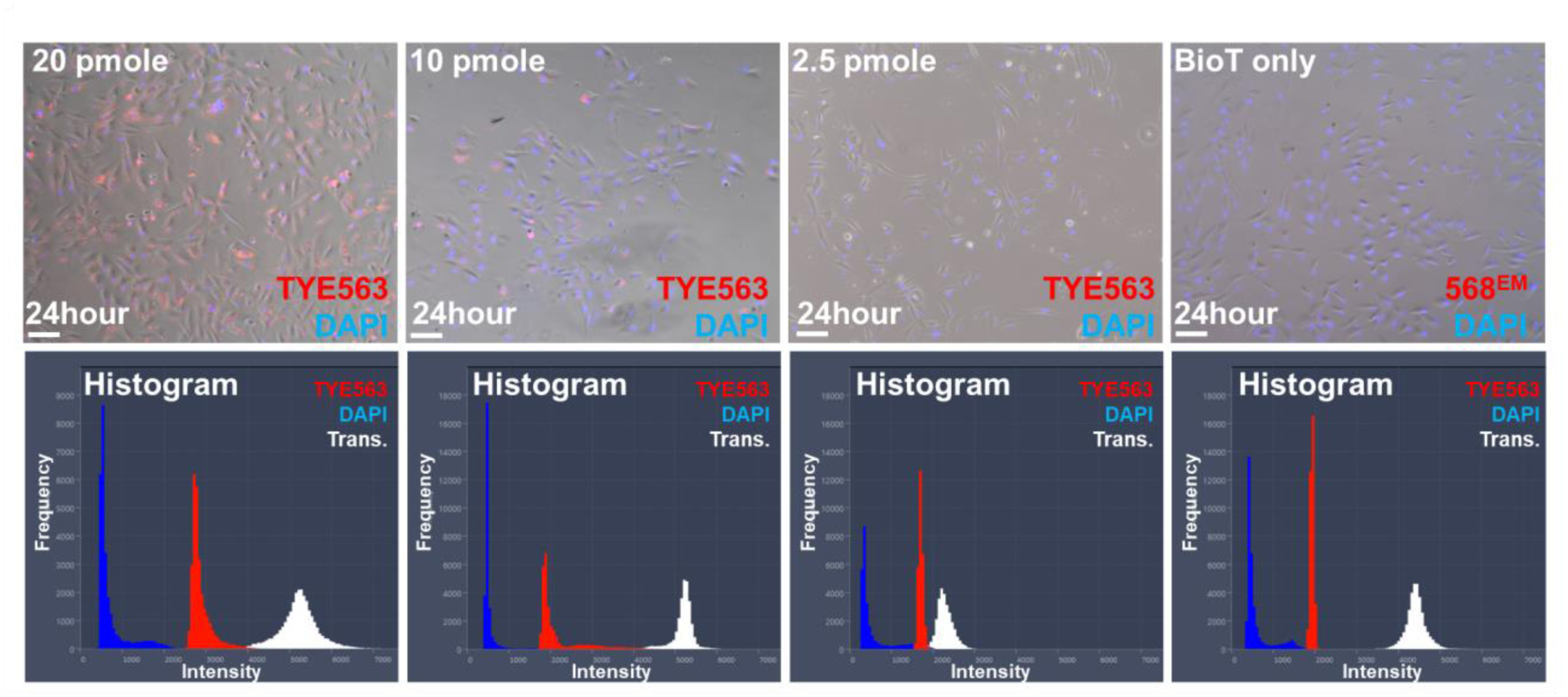
Transfection efficiency of DsiRNA Trifecta reagents in LM7 cells. After 24 hours of incubation, the transfection efficiency of 20pmole of TYE563-labeled DsiRNA was greater than 90%. Bar = 100µm Below: Histograms revealed that the 20pmole concentration comprised a bigger fraction of cells with a higher intensity.

**Figure S8.**
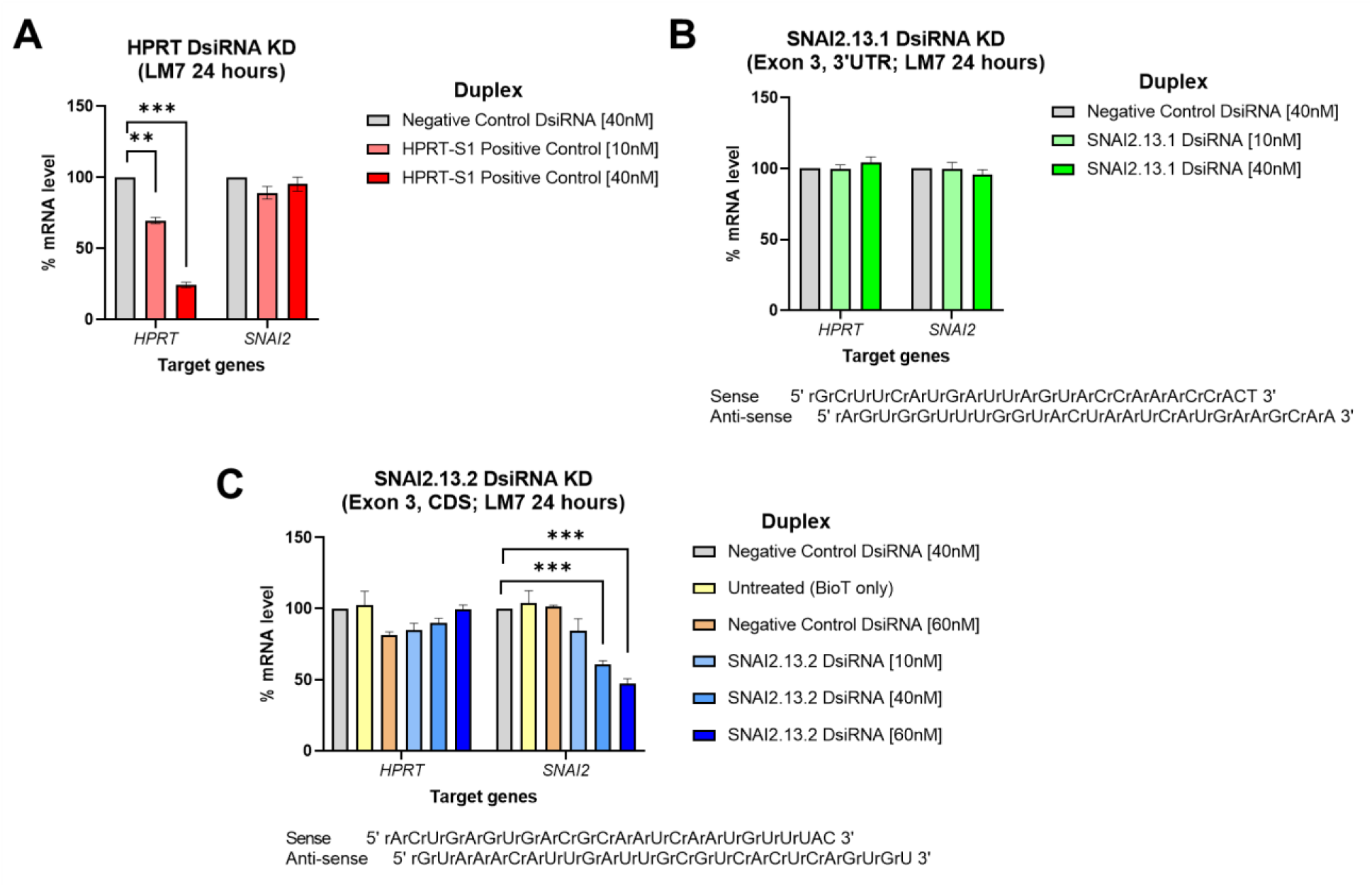
DsiRNA knockdown of *SNAI2* and control *HPRT* in LM7 cells. A) *HPRT* cleavage by DsiRNA. B) Using the SNAI2.13.1 duplex, DsiRNA aimed to inhibit *SNAI2* expression. C) Successful *SNAI2* DsiRNA knockdown with the SNAI2.13.2 duplex.

**Figure S9.**
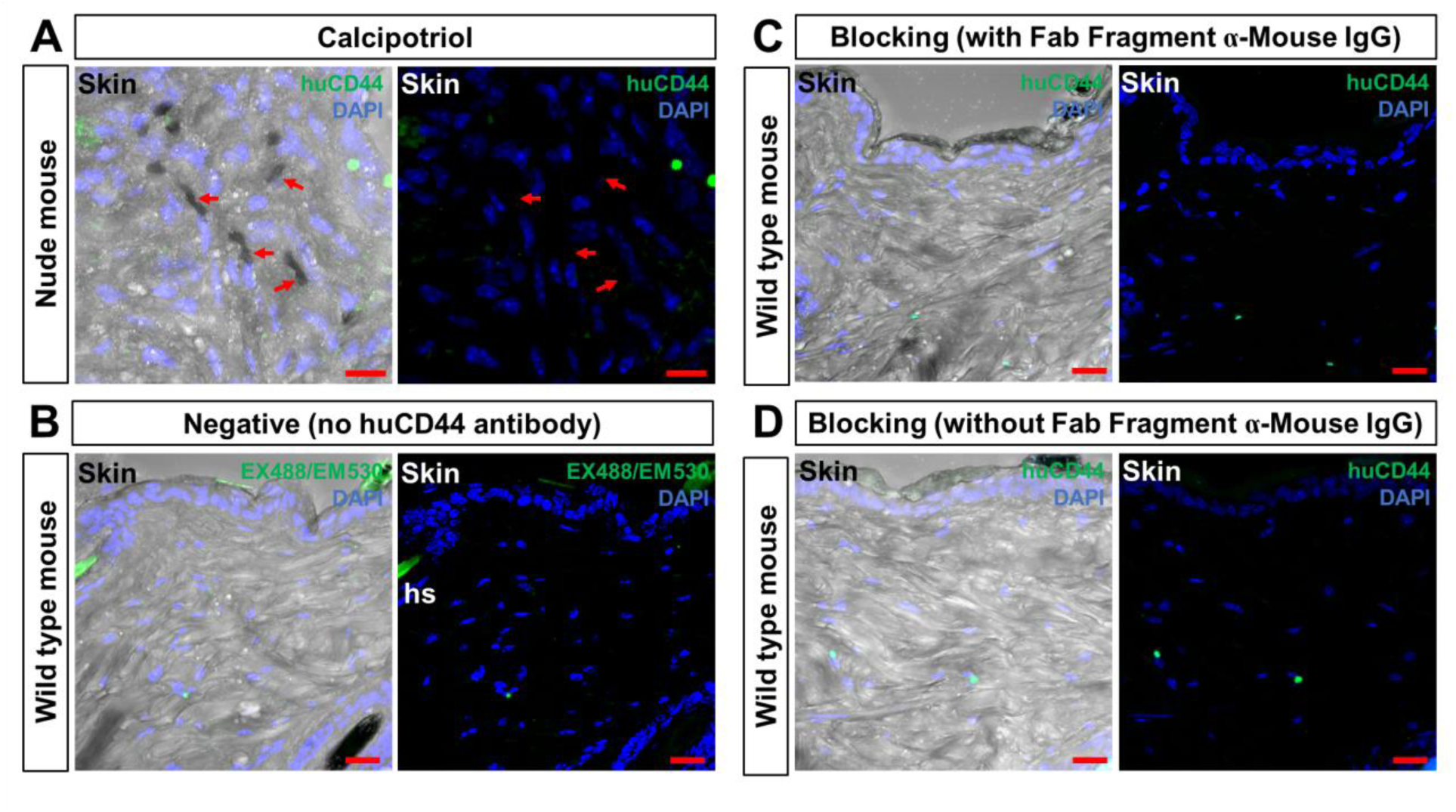
huCD44 antibody testing. A) Application of huCD44 antibody against calcipotriol-treated LM7 xenograft mouse skin. Red arrows depict red blood cells that are negative for huCD44. Bar = 10µm B) Negative control. Wild type mice skin sections were treated similarly but without huCD44 antibody. Typical autofluorescence of hair shafts (hs) are seen. Bar = 20µm C) huCD44 background analysis with mouse Fab fragment blocking. Bar = 20µm D) huCD44 background analysis without mouse Fab fragment blocking. Bar = 20µm

**Figure S10.**
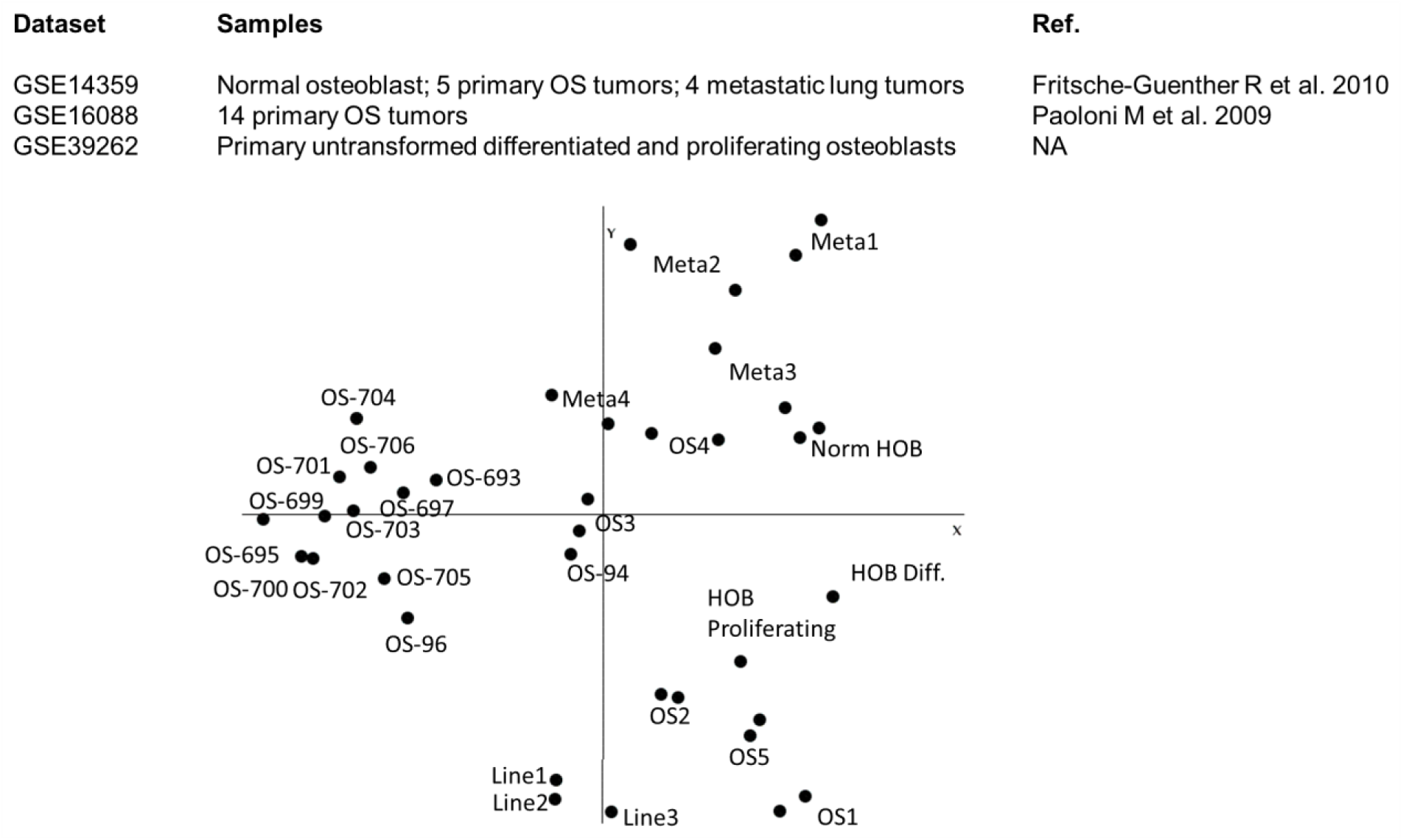
Principal component analysis of osteosarcoma and normal samples used for comparative gene expression analysis. The GEO series accession numbers are referenced for each sample set.

**Figure S11.**
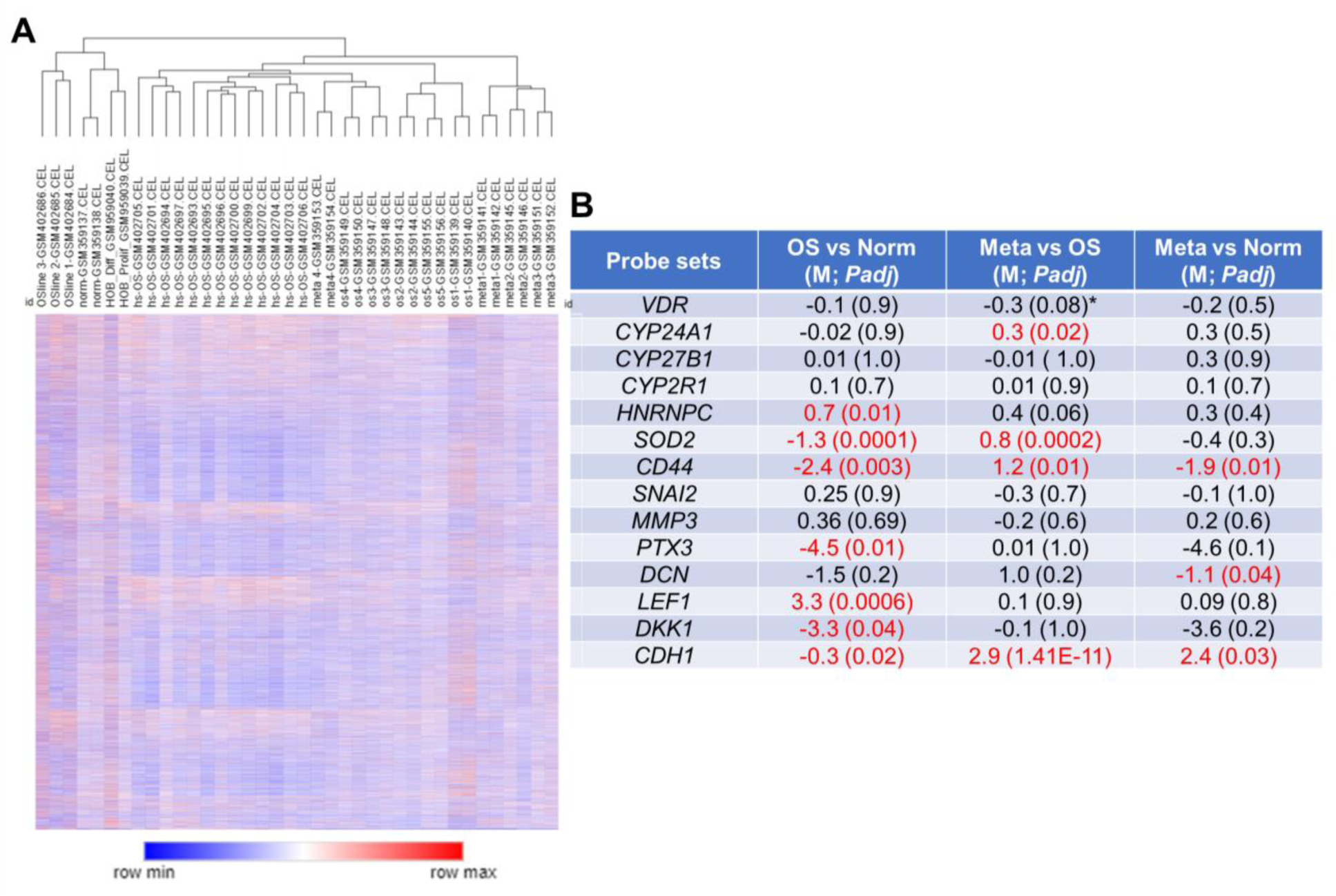
Comparative analysis between normal bone tissue, and primary osteosarcoma and metastasized lung tumors. A) Hierarchical clustering of cDNA arrays of human tumor samples and OS cell lines using Euclidean distance and full linkage. The GEO sample accession number is assigned to each sample (Corresponds to Figure). BioConductor software for traditional 3’ arrays were used for preprocessing and analysis of Affymetrix GeneChip data. RMA (Robust Multiarray Analysis) was utilized for array preparation. Moderated t-statistics (limma) were utilized to identify genes with differential regulation. B) The chart displays the M value, which is the log-ratio of the probeset intensities, as well as the corrected *p* values of numerous comparisons. Normal has normal osteoblast cells and normal tissue. Red indicates signatures that statistically differed from the others.

## Supplementary Material

File S1. Functional enrichment annotation analysis of MG63 unique H3K27ac target genes

File S2. Functional enrichment annotation analysis of LM7 unique H3K27ac target genes

Movie S1. Vehicle-treated LM7-bearing nude mice PET scan.

Movie S2. Calcipotriol-treated LM7-bearding nude mice PET scan.

---

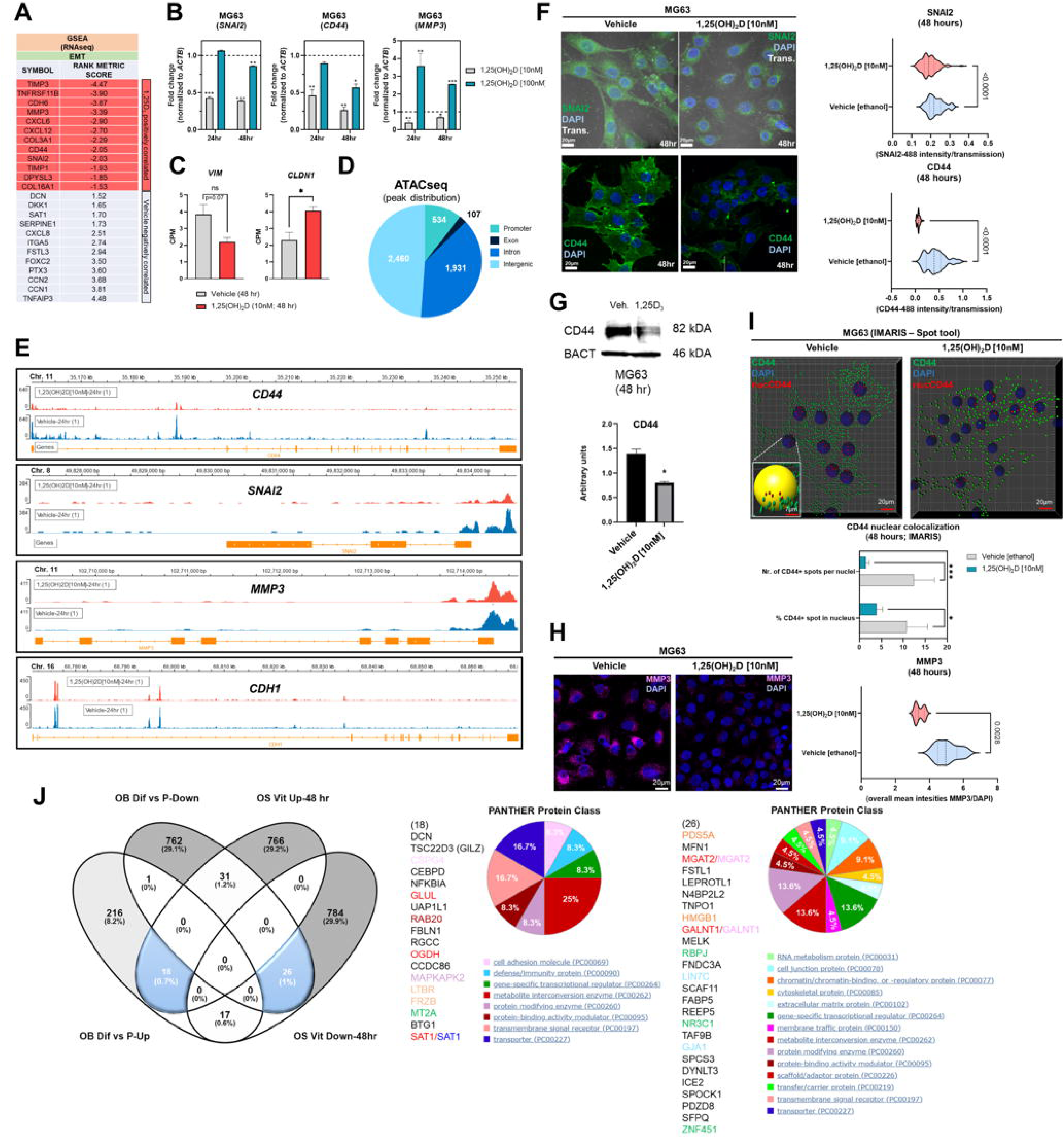

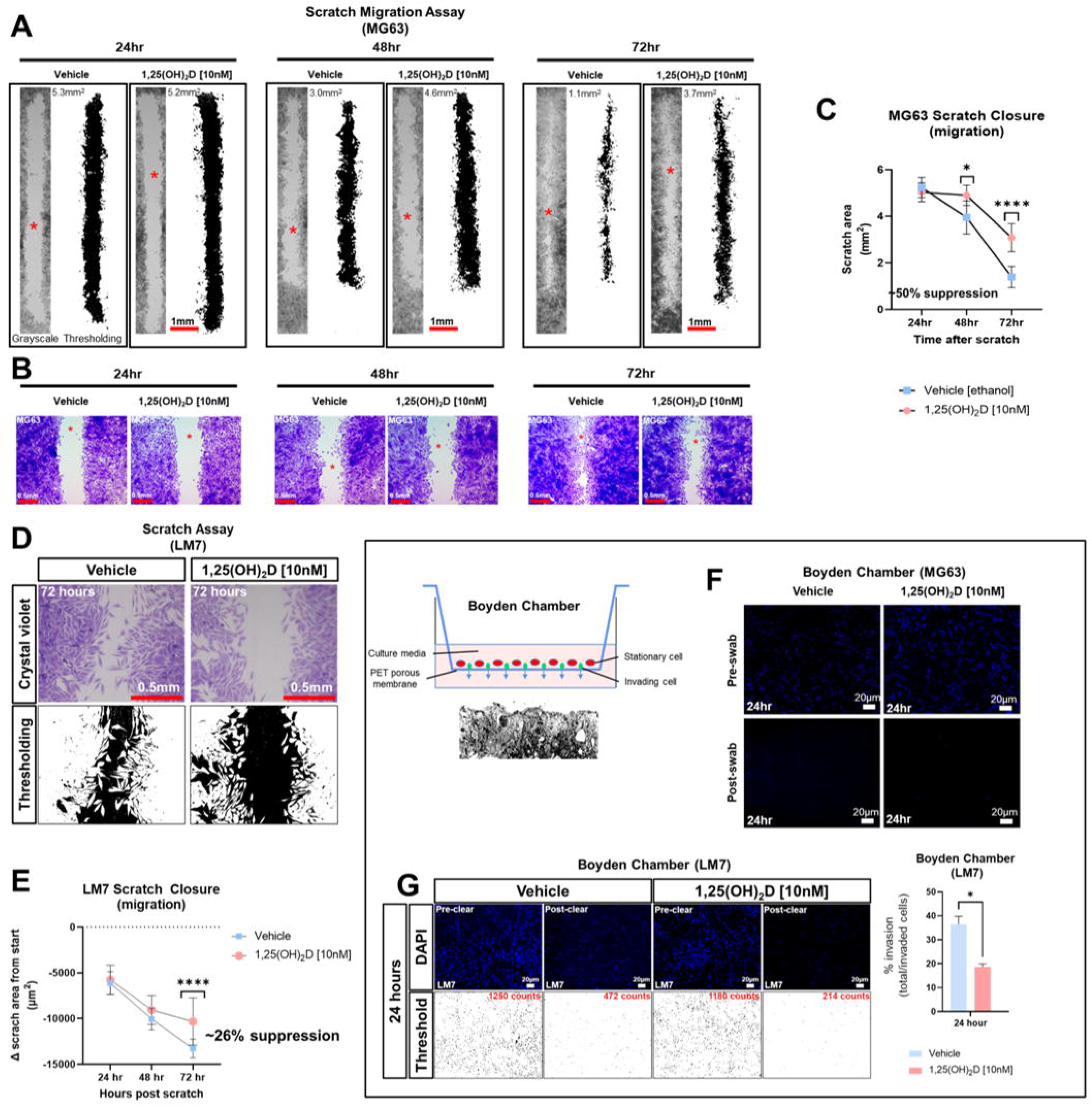

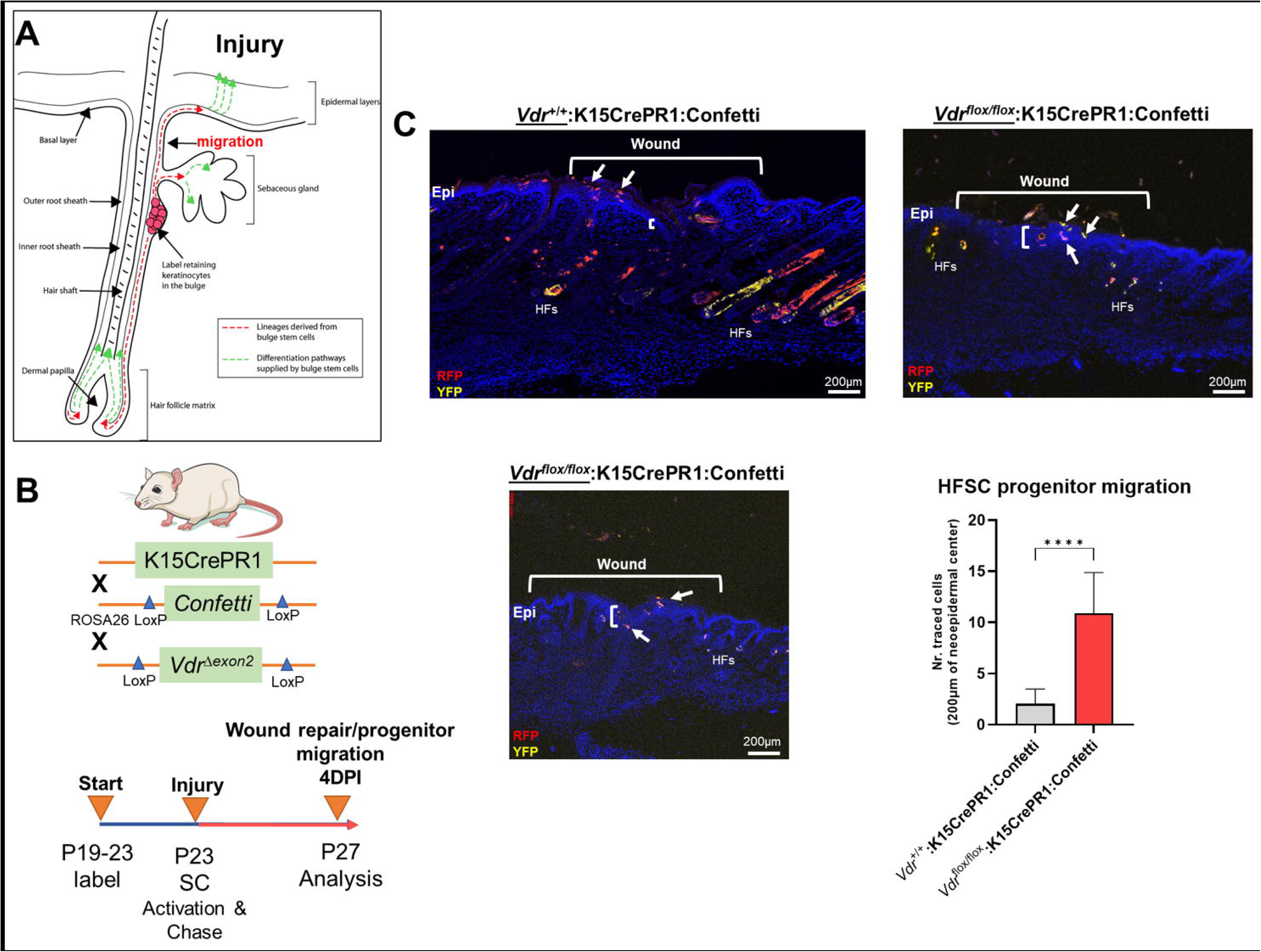

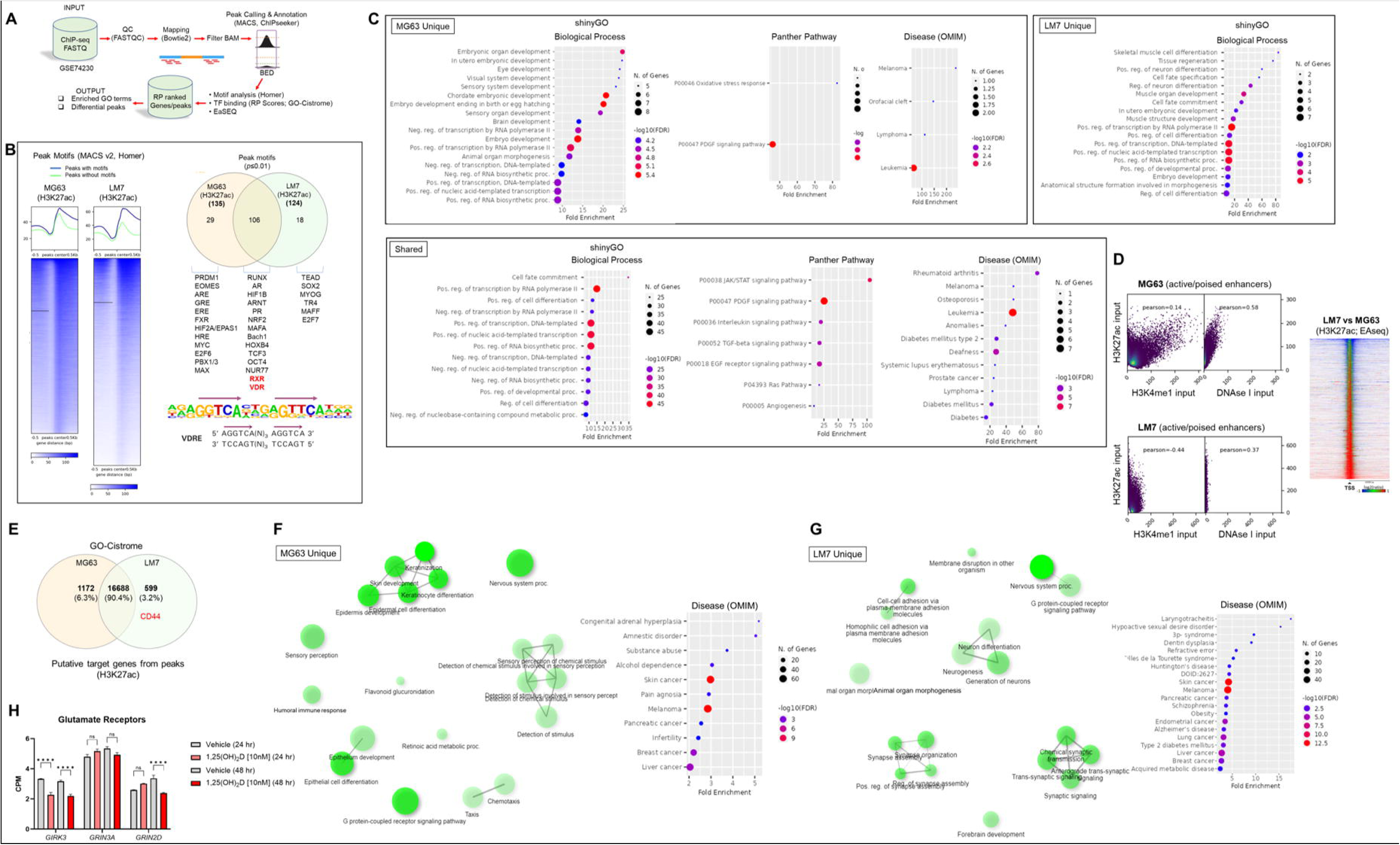

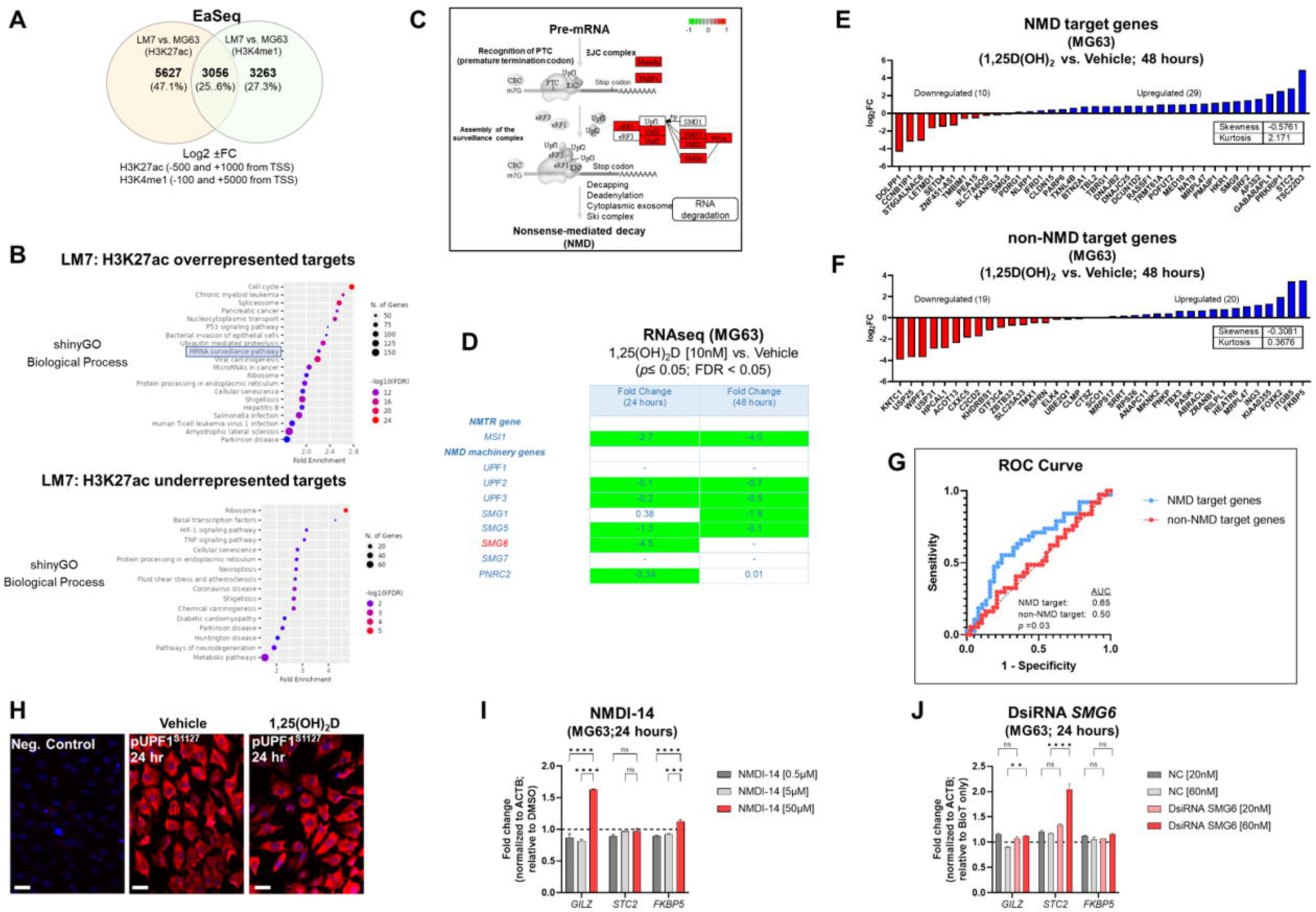

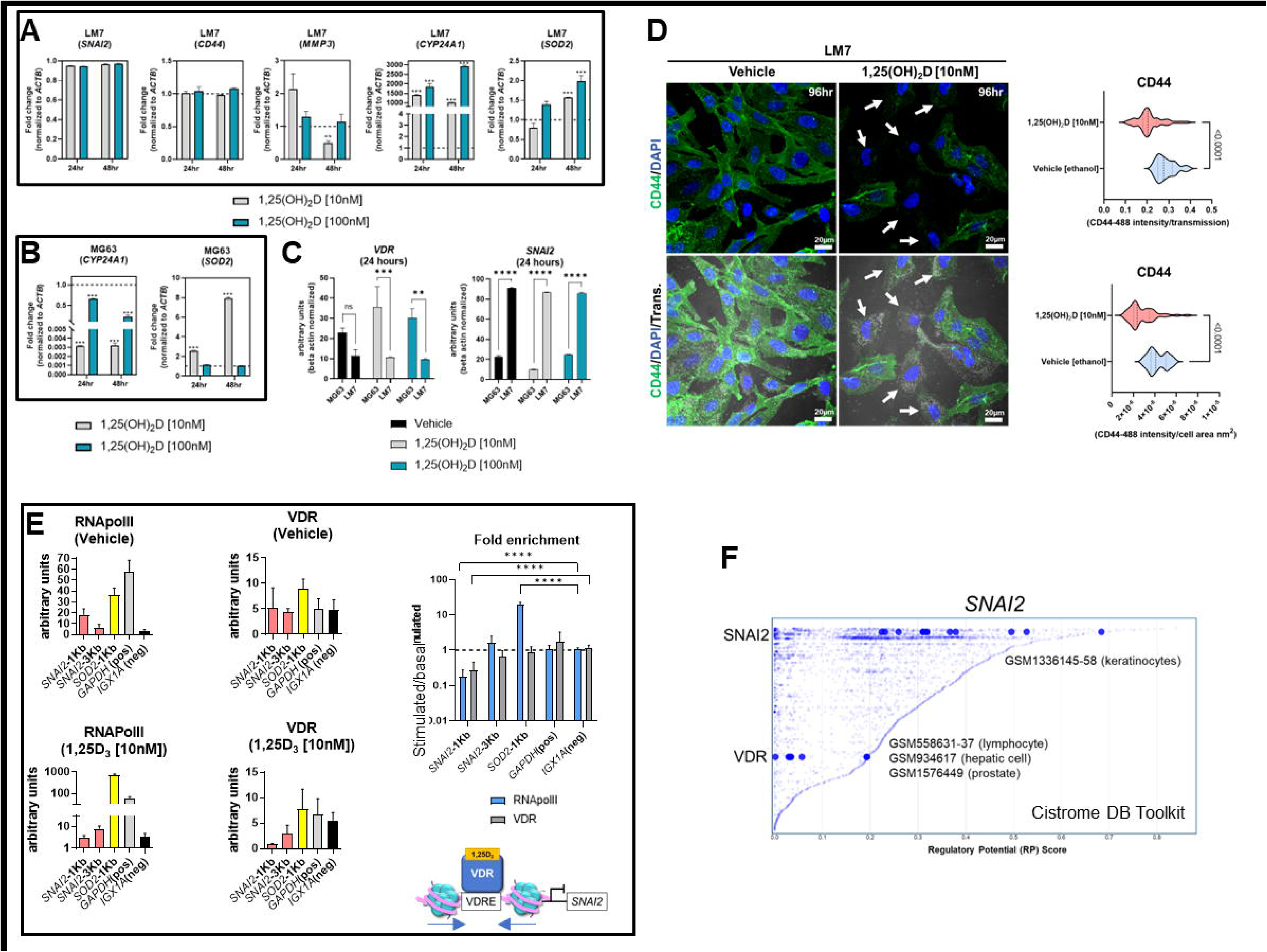

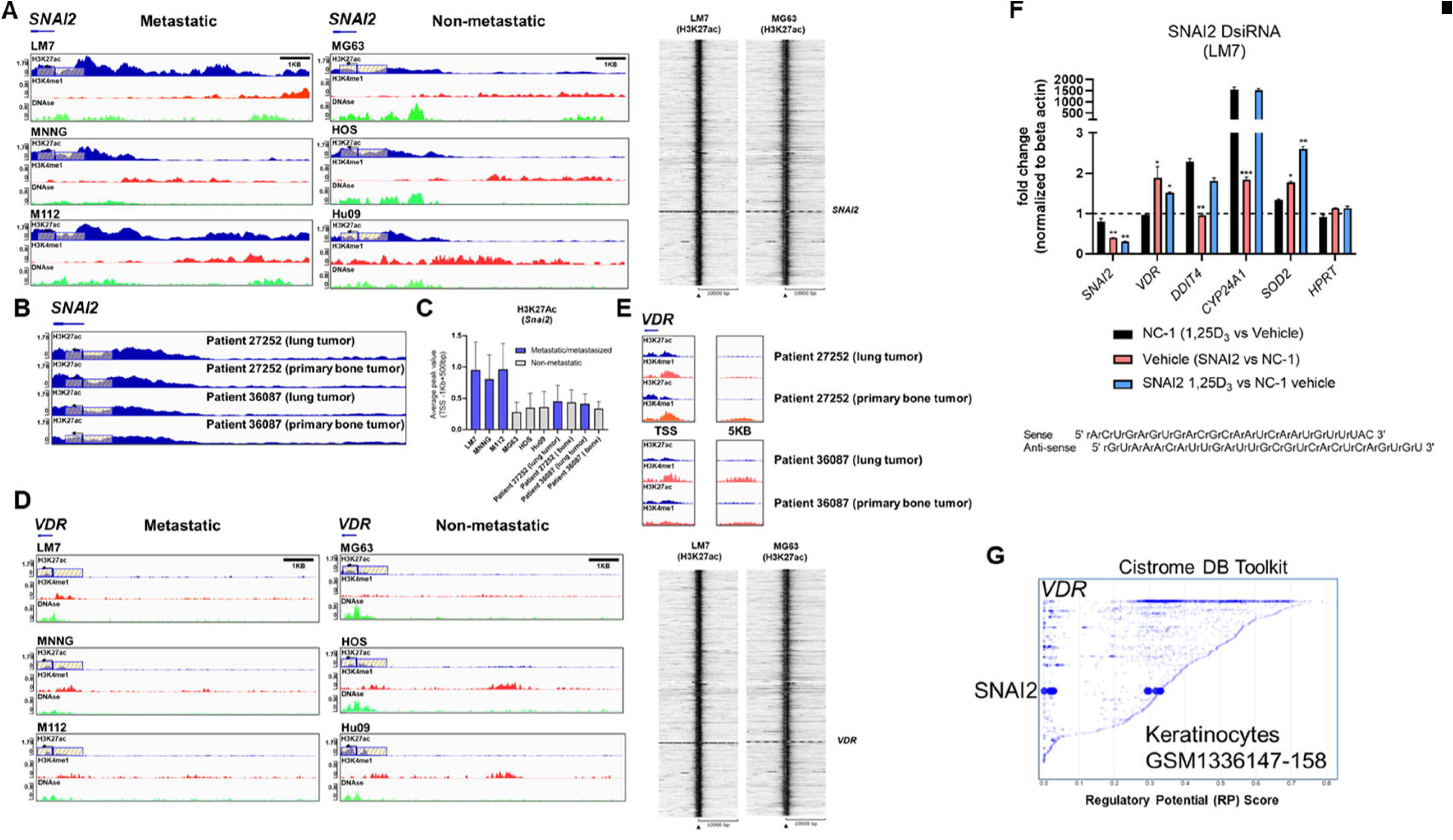

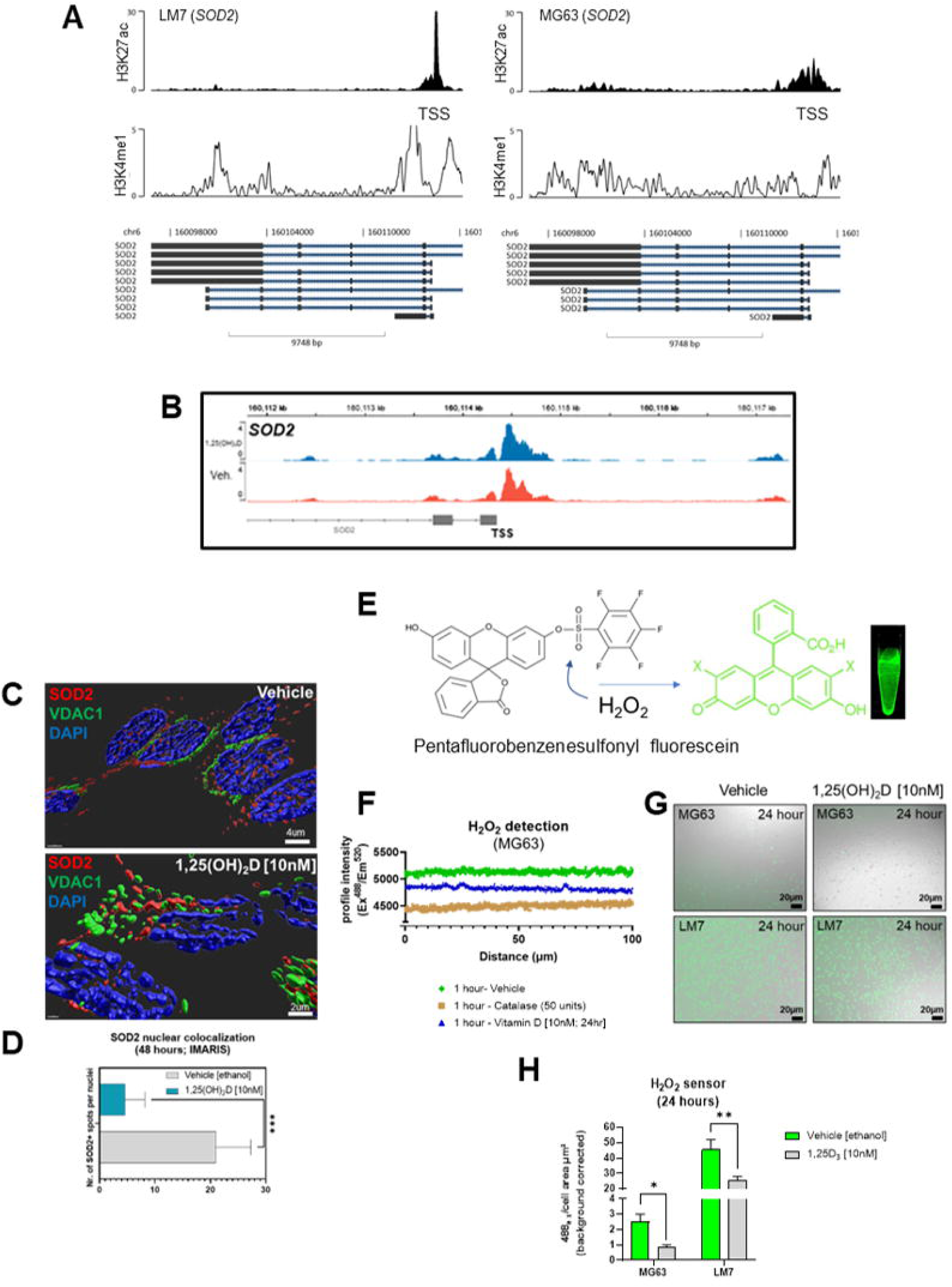

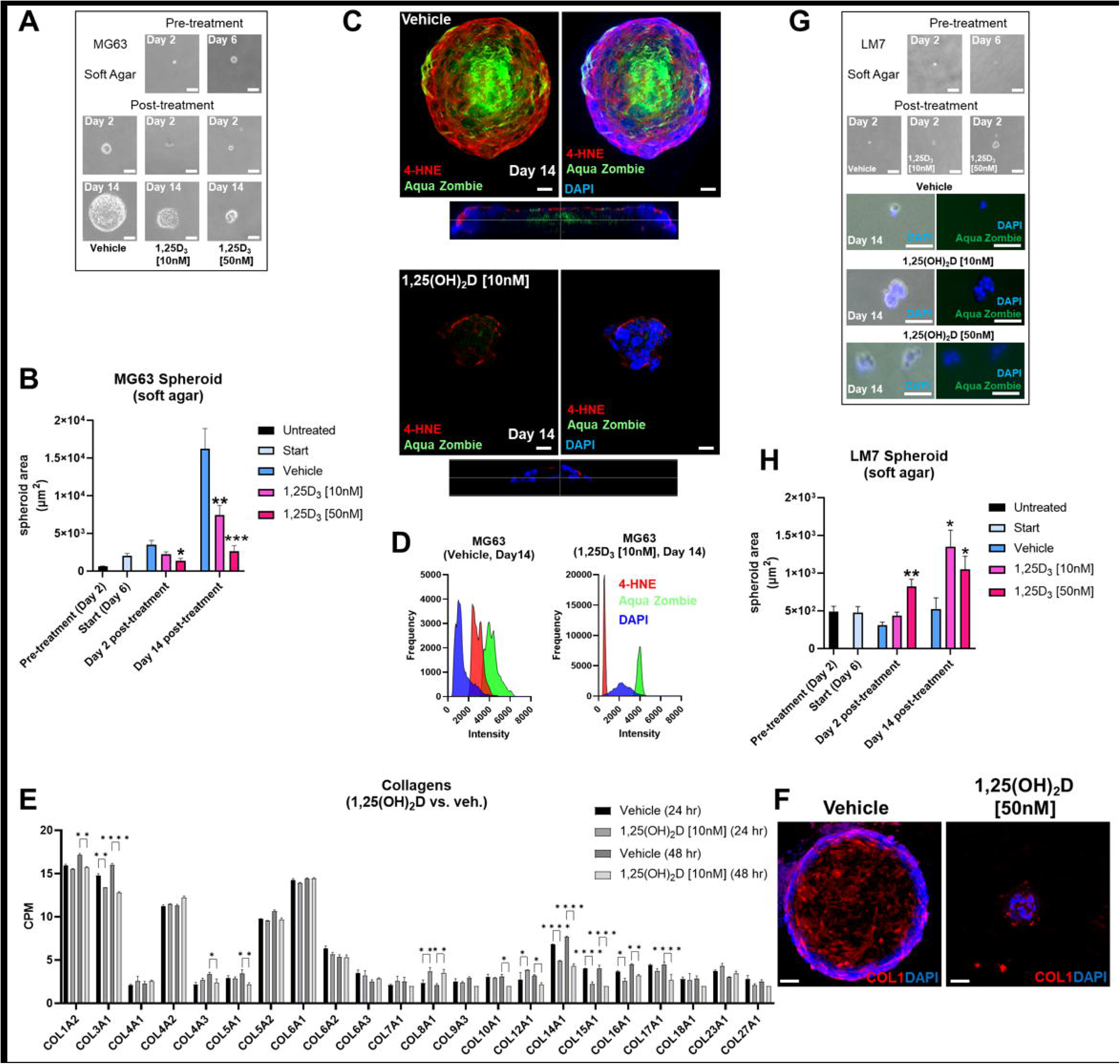

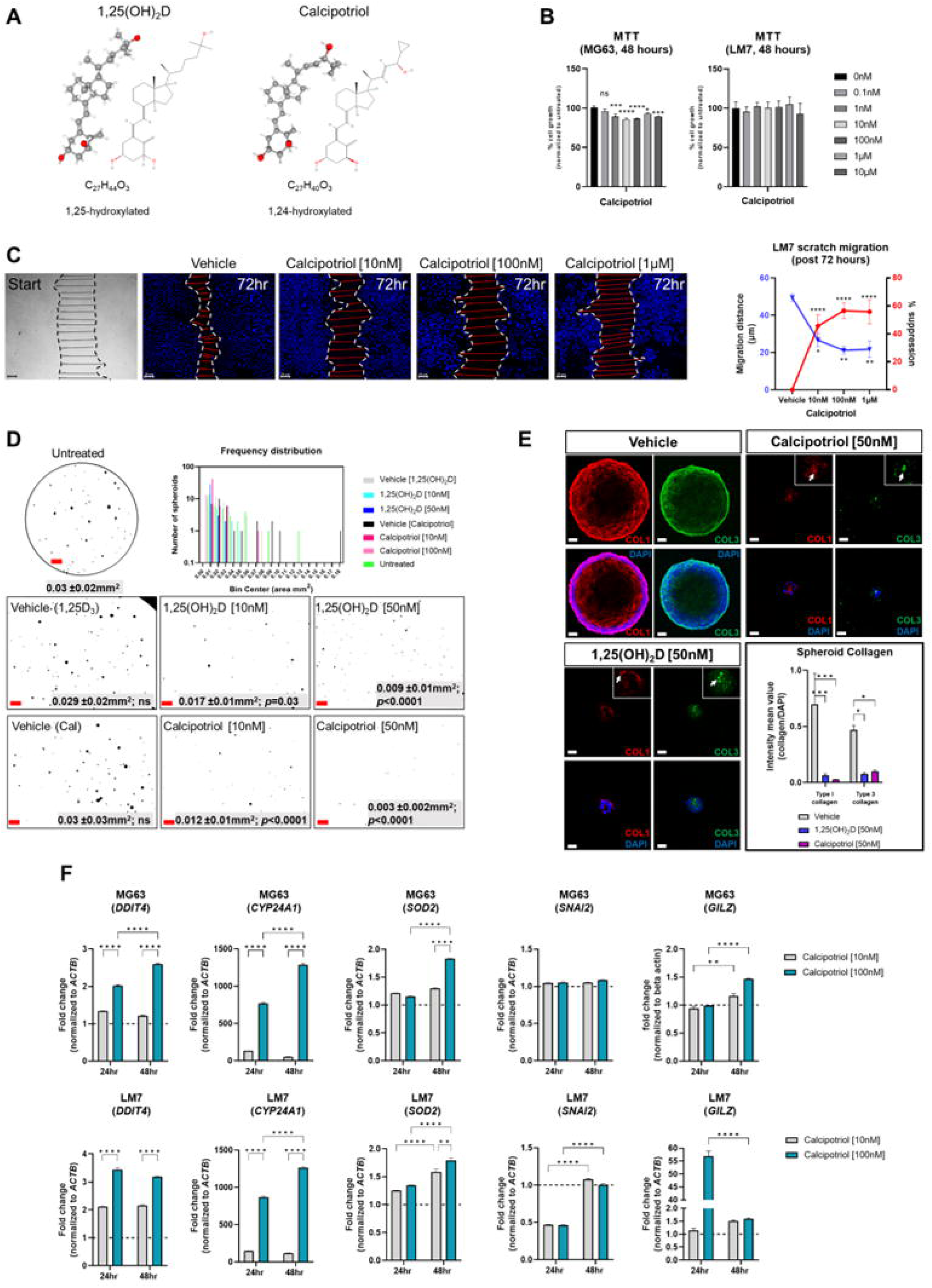

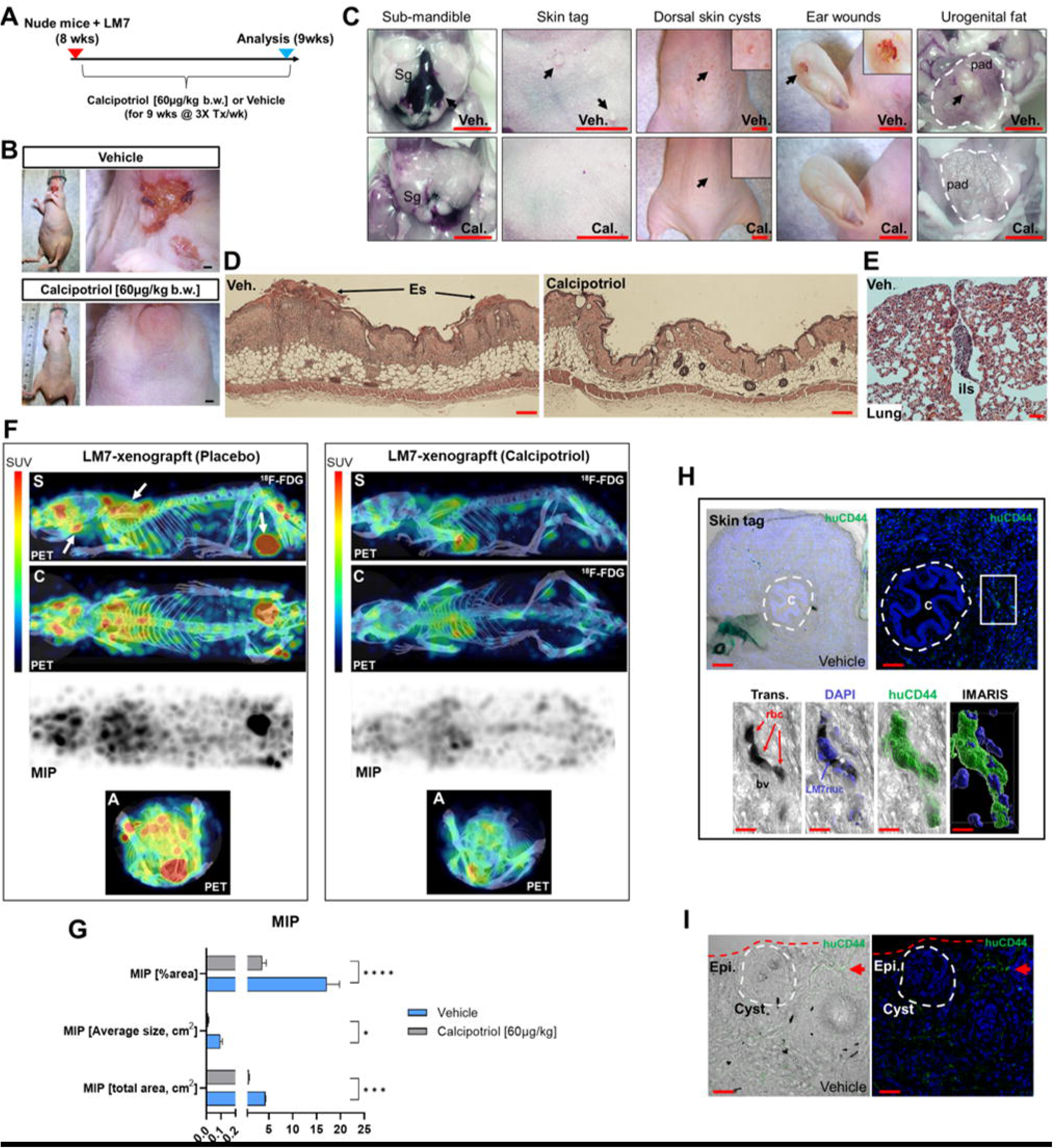

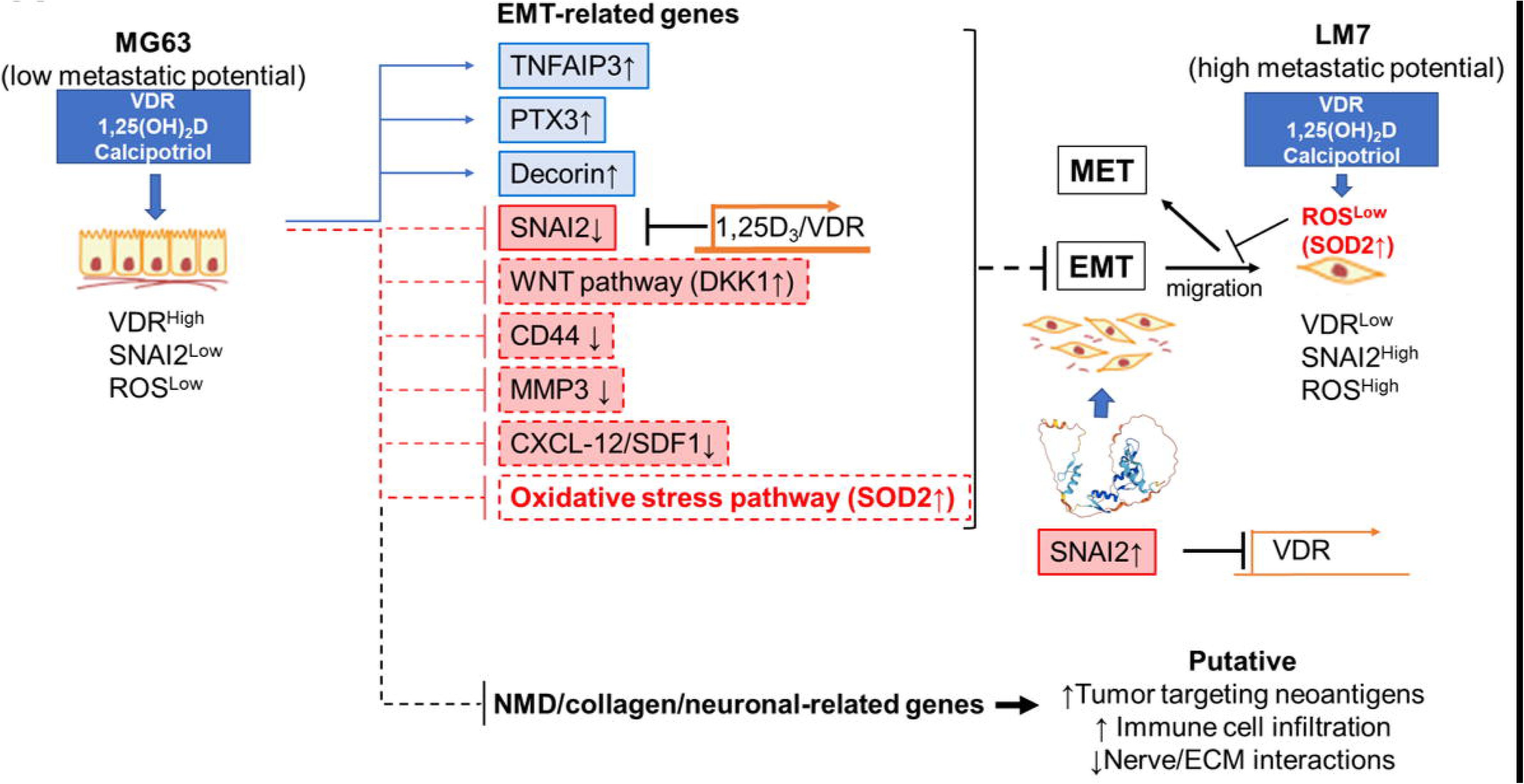

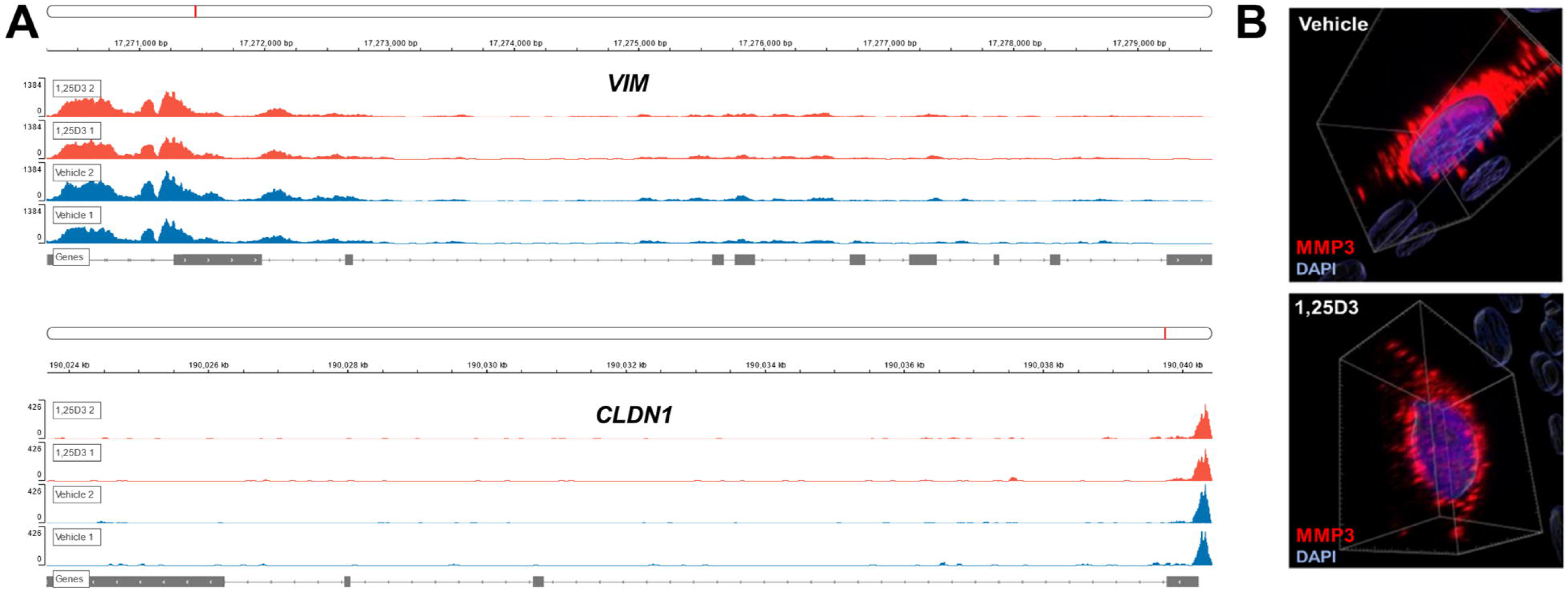

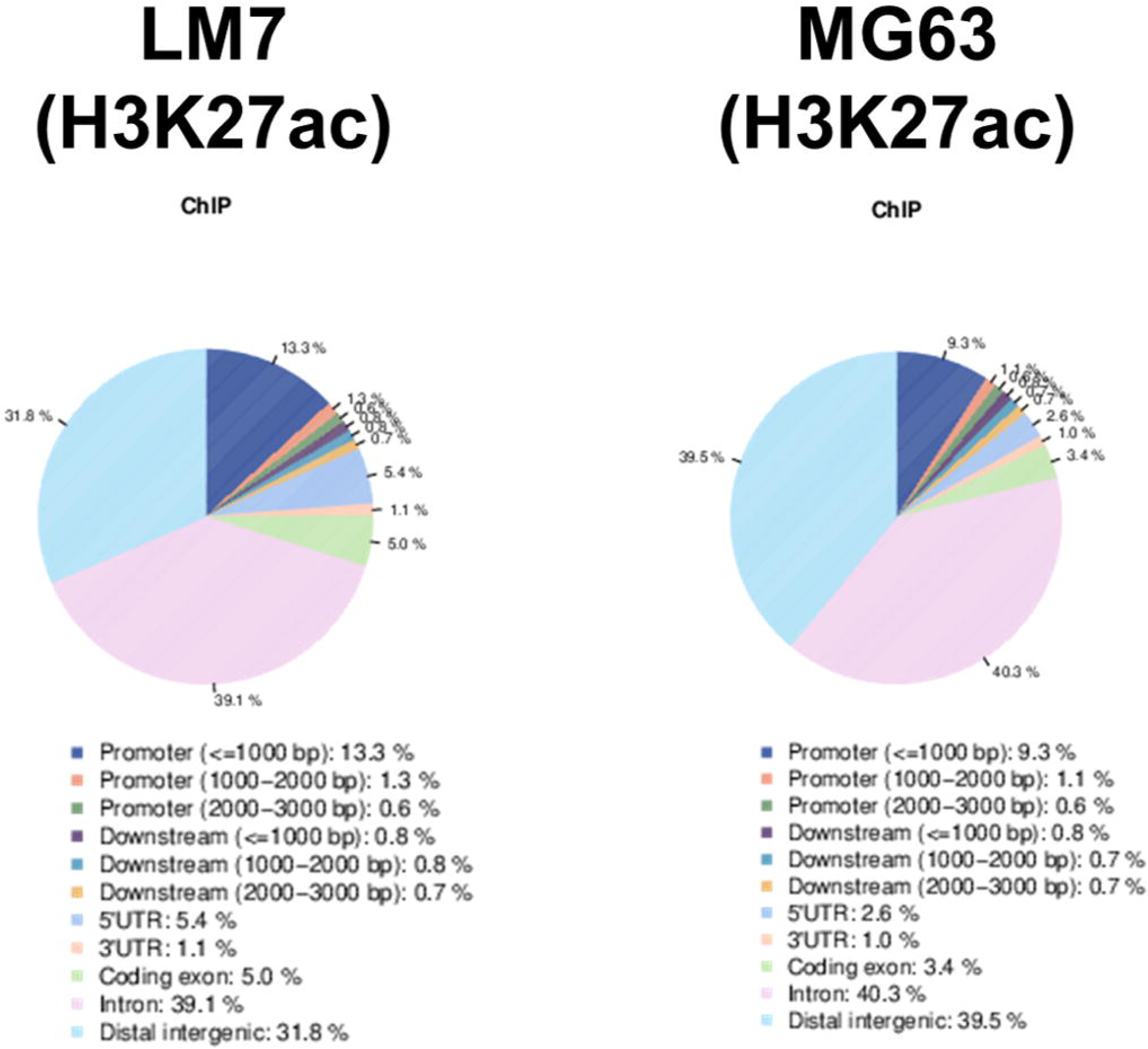

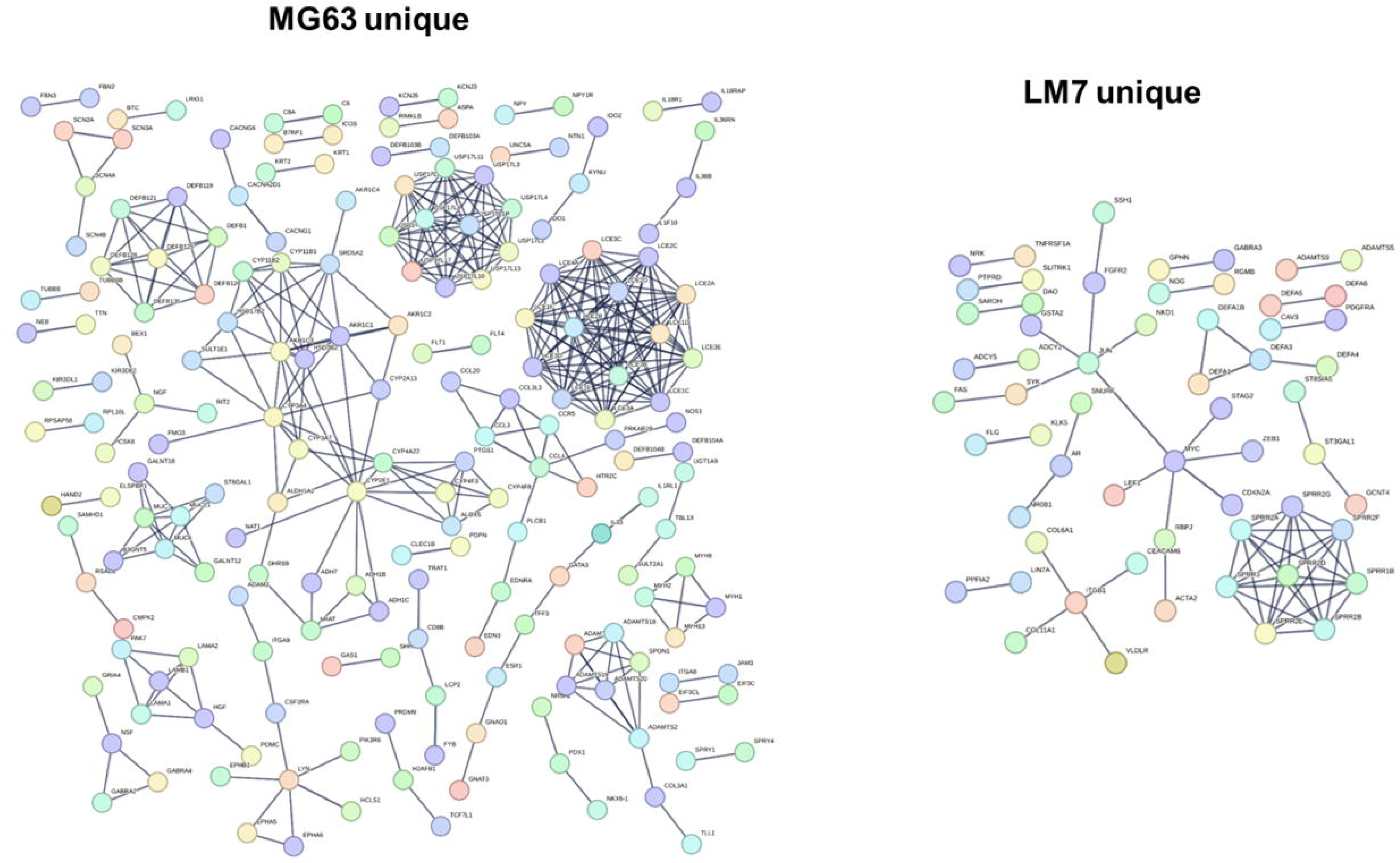

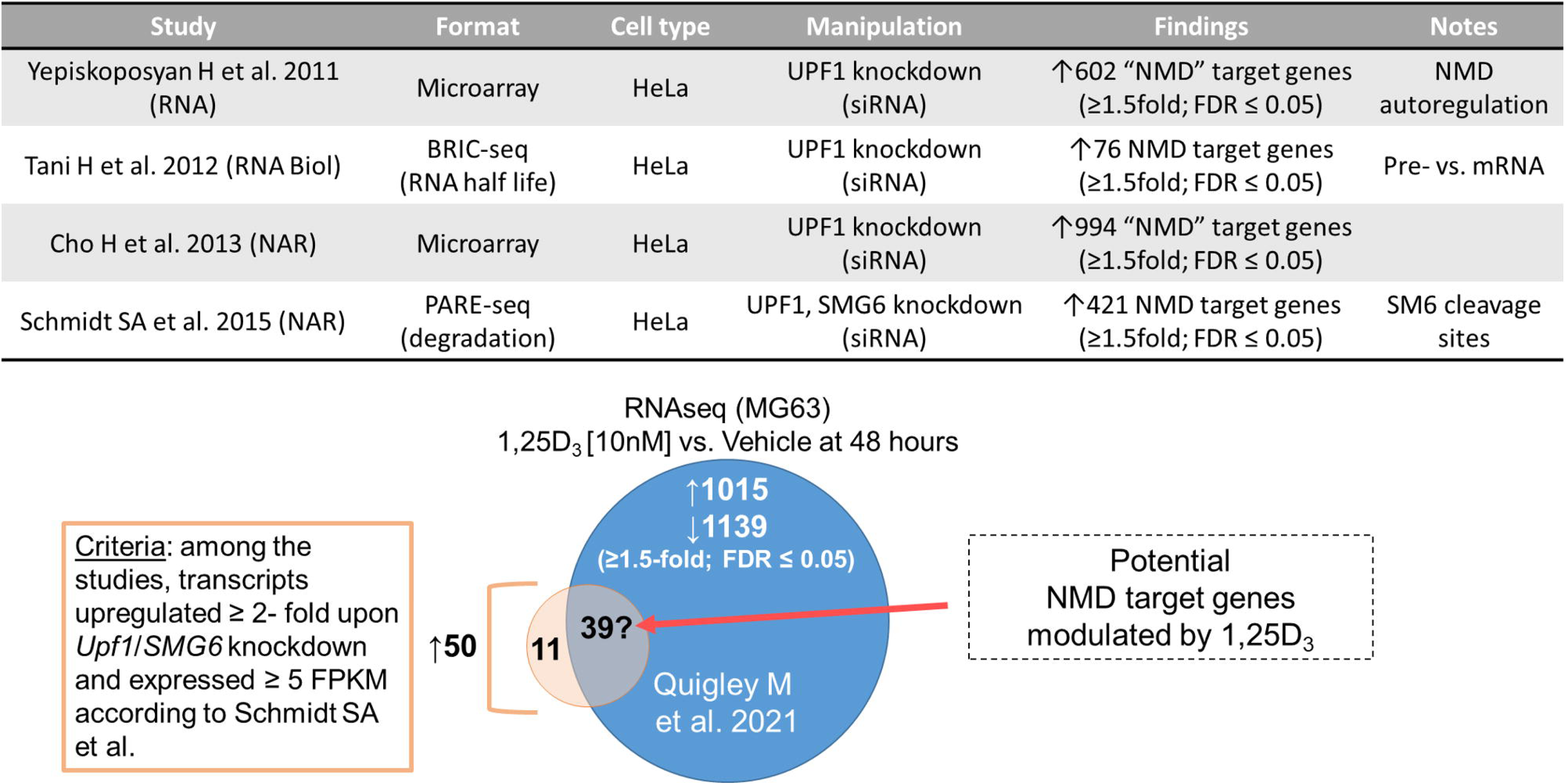

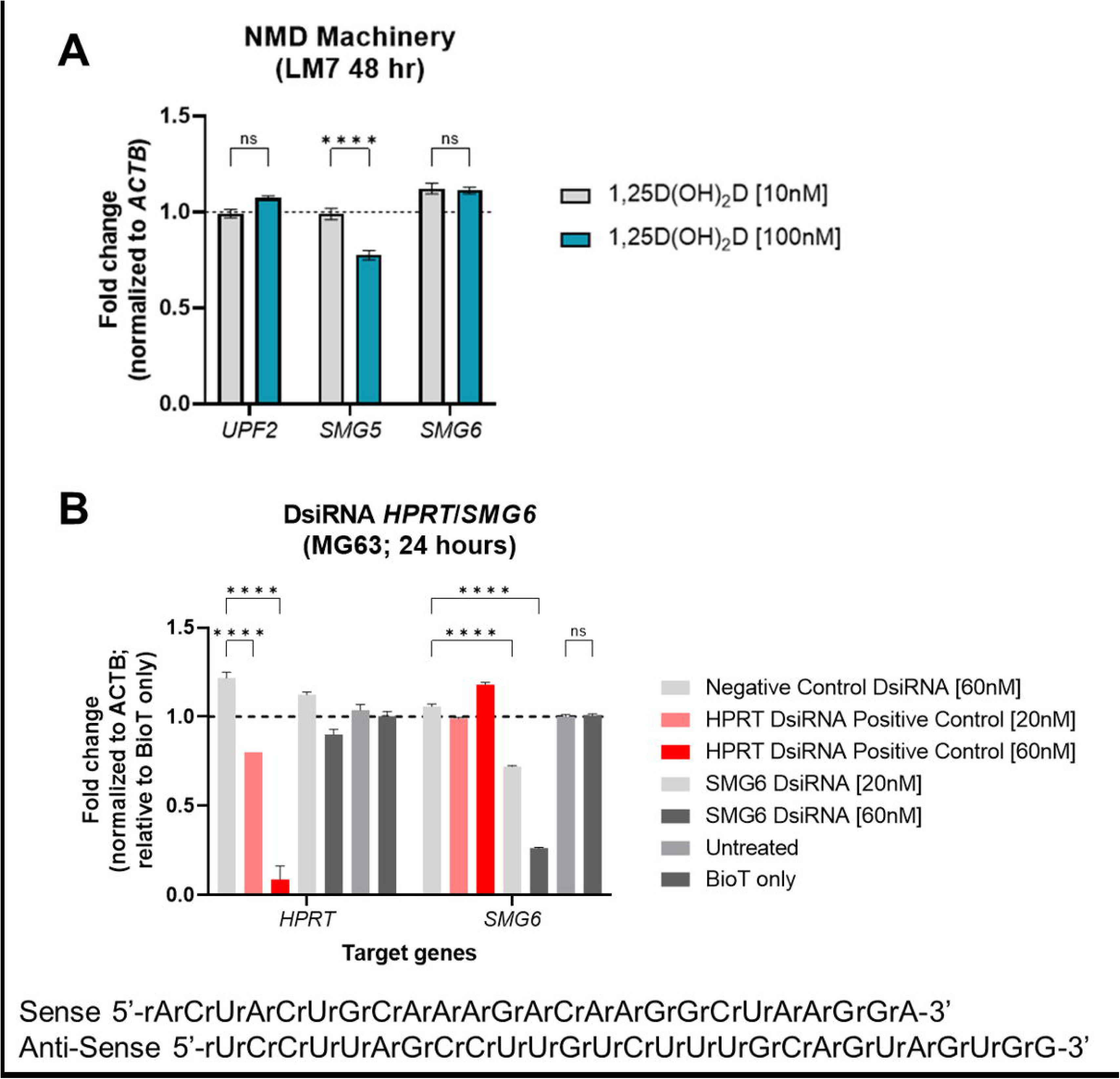

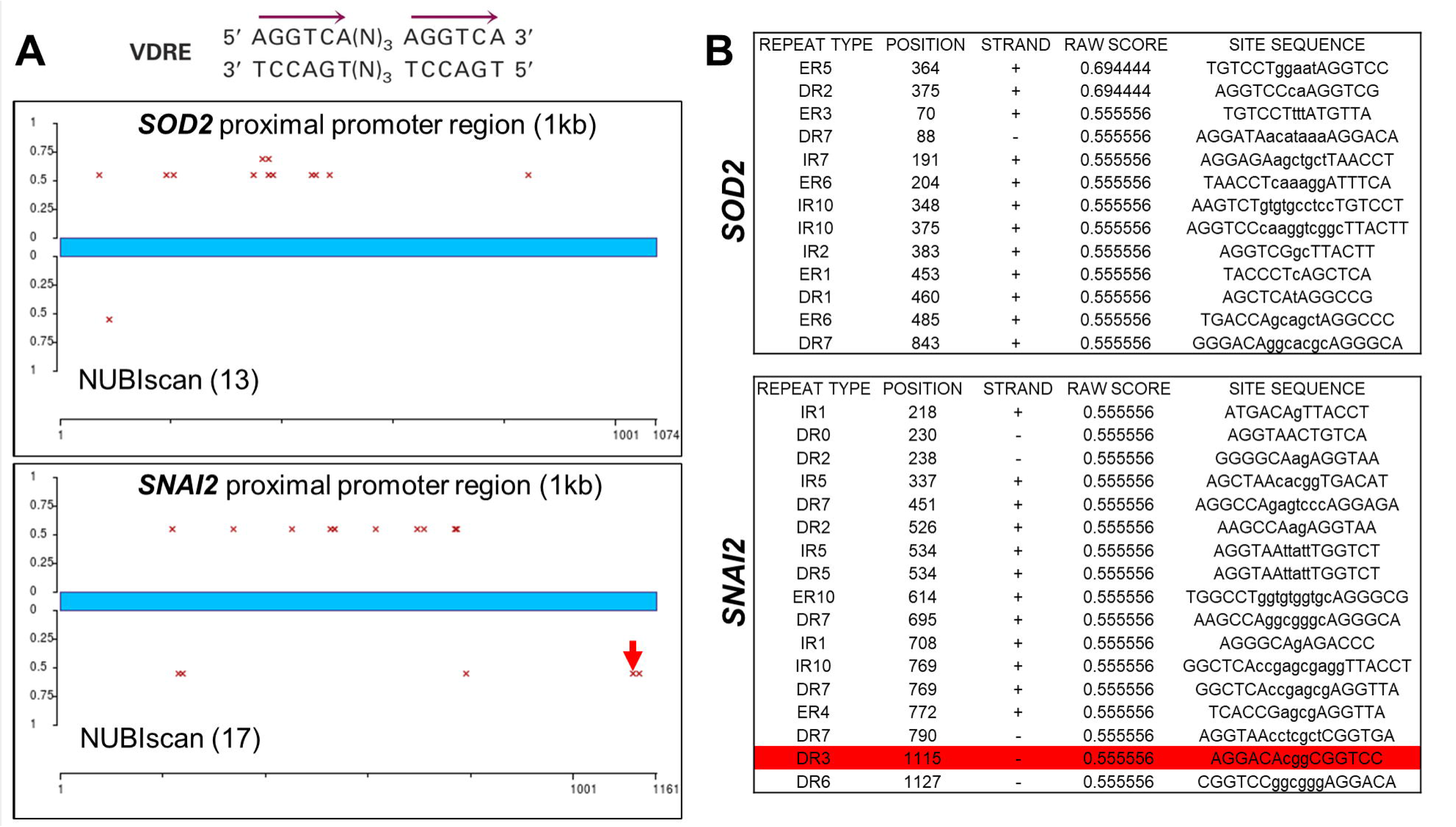

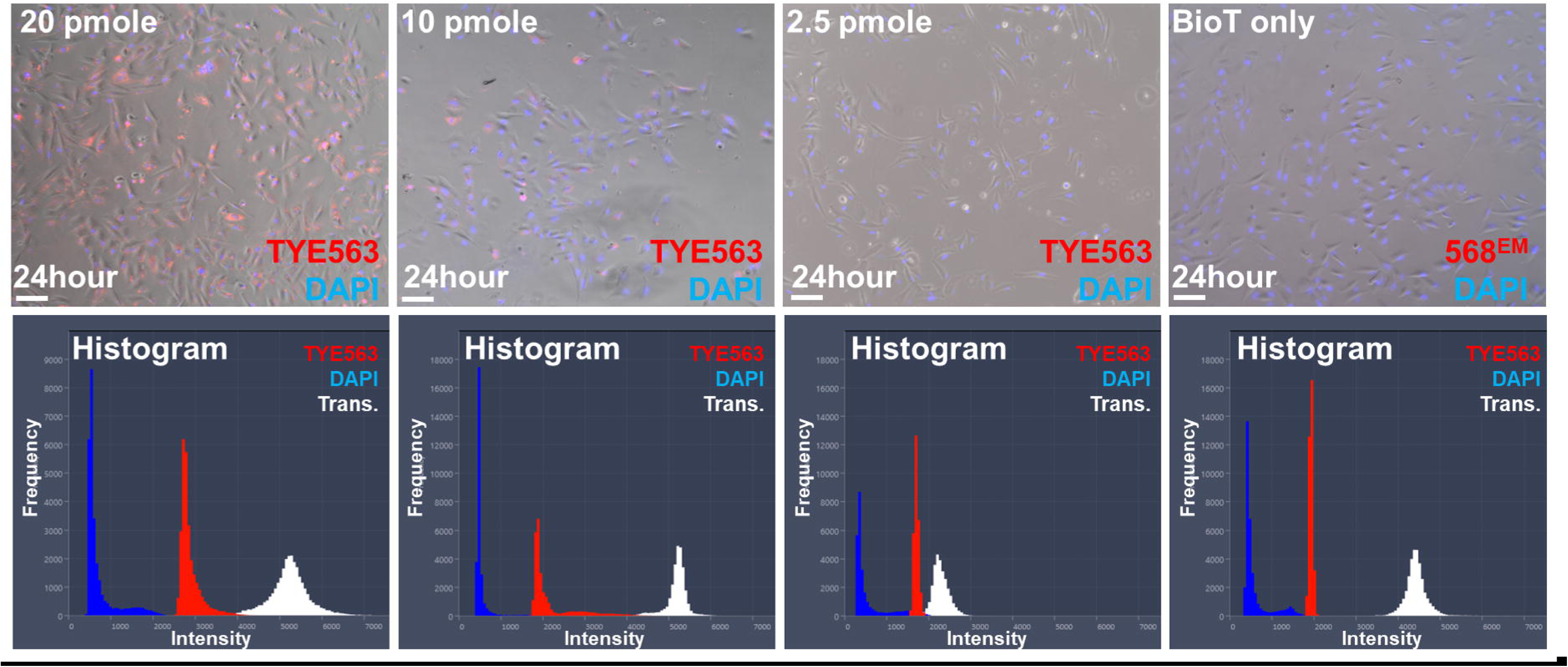

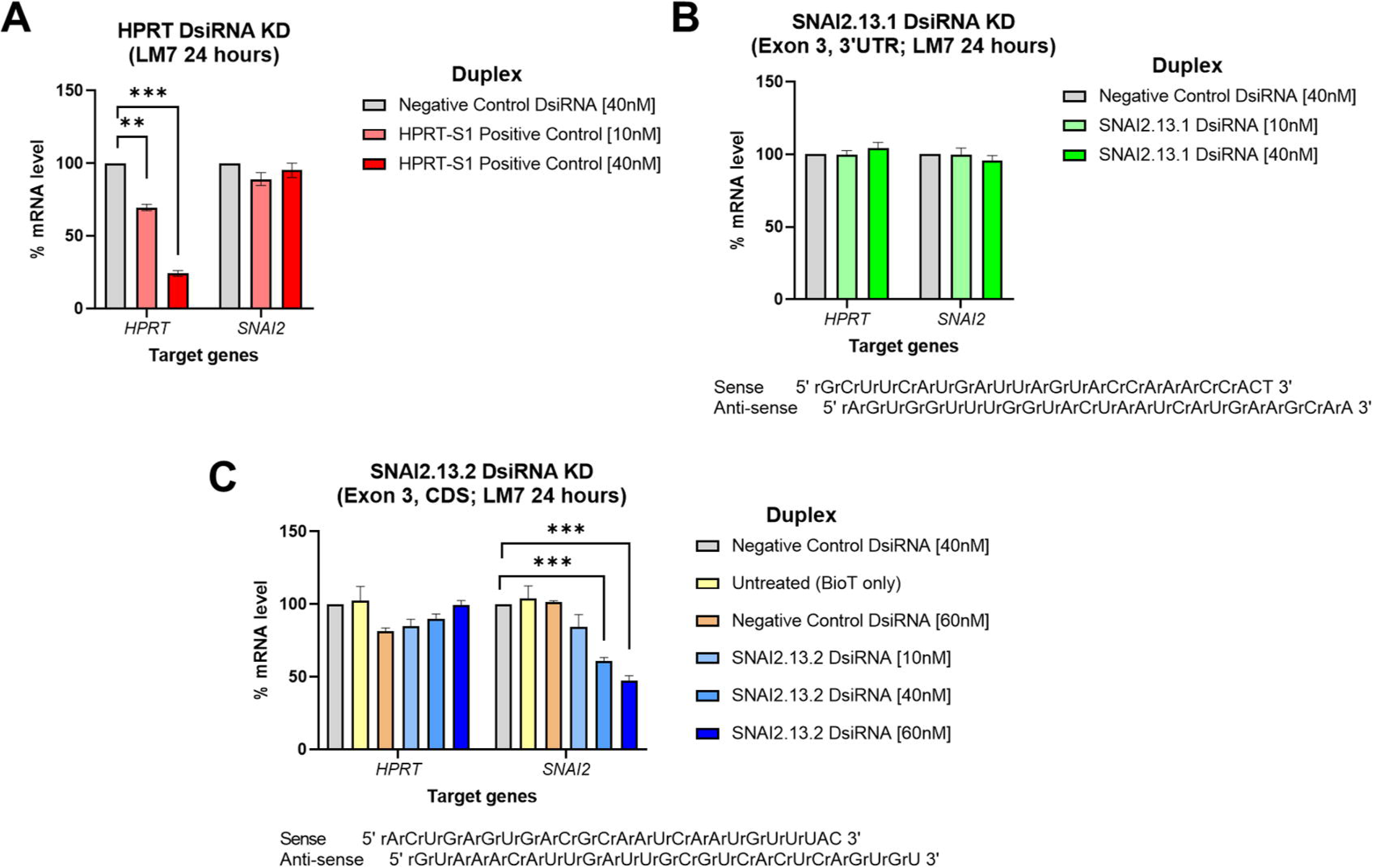

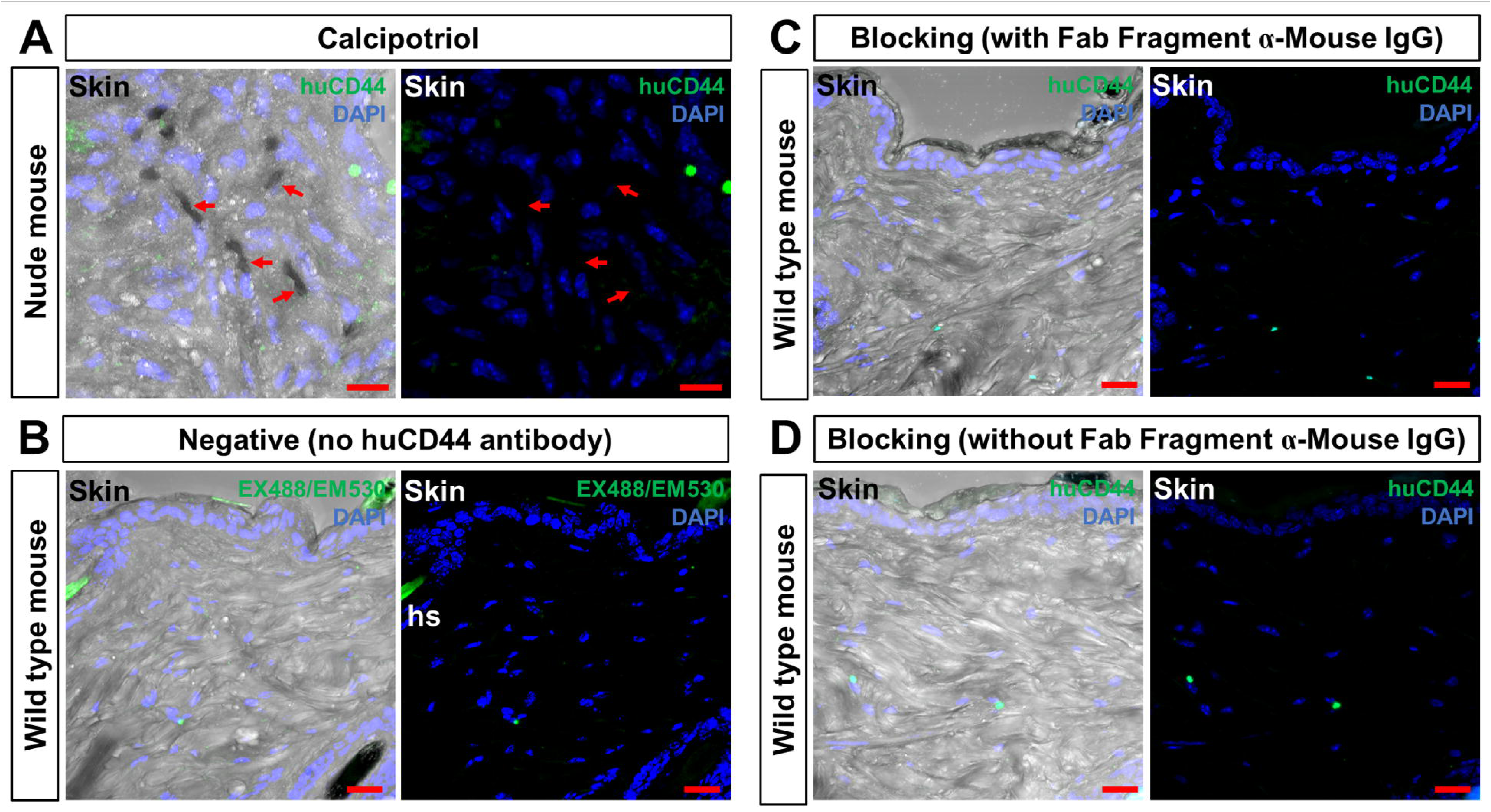

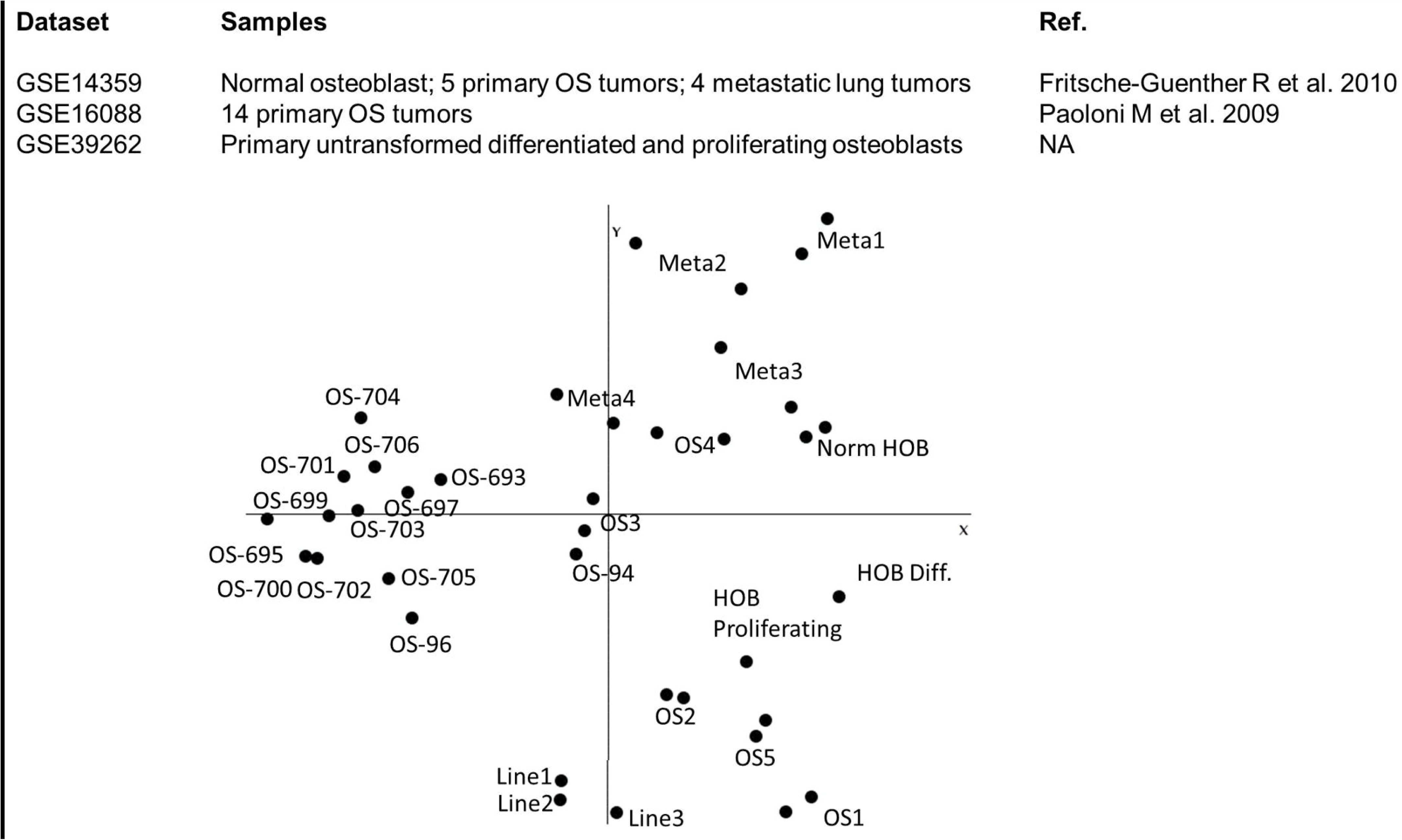

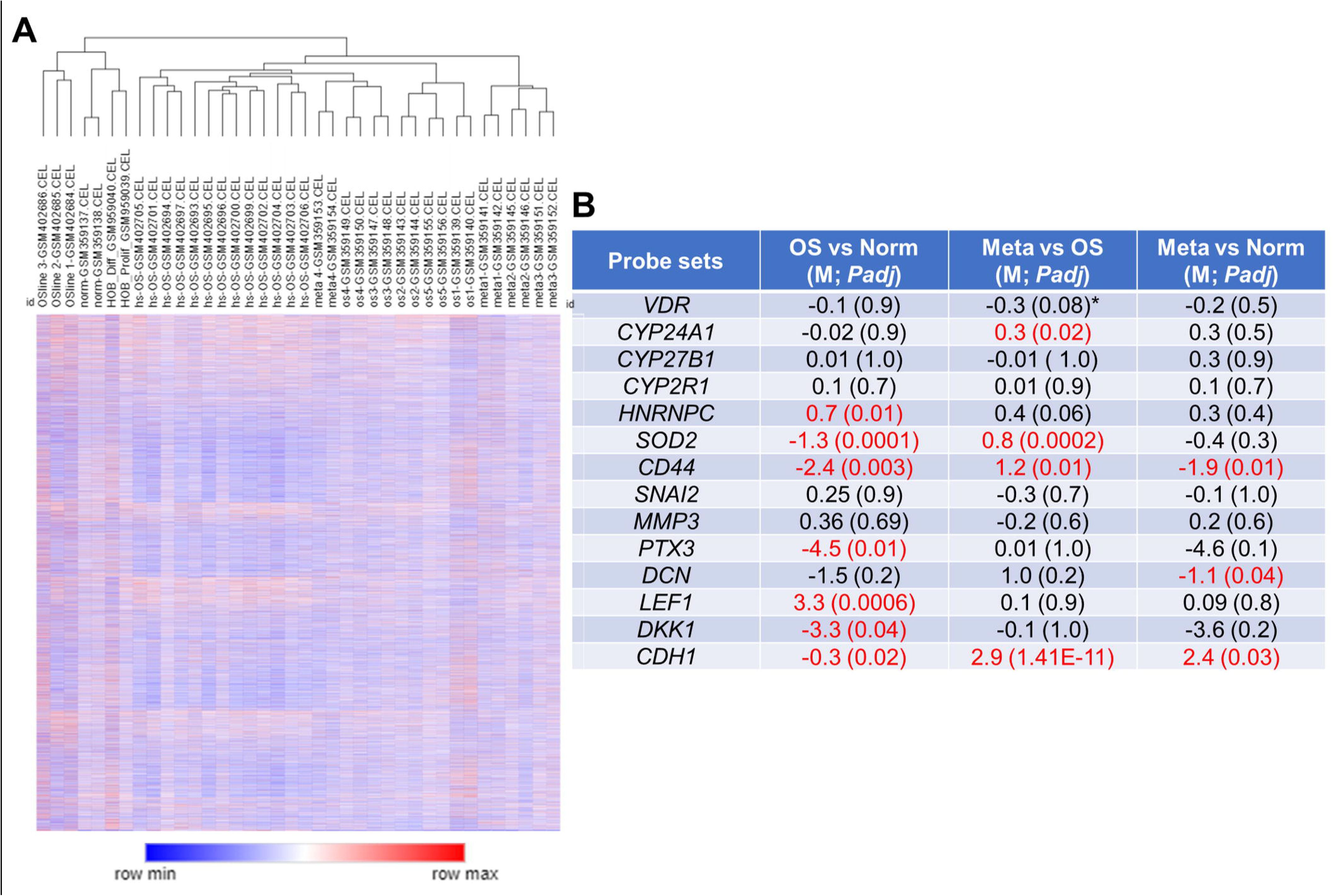

