## Supplemental File 1 for "Vitamin D inhibits osteosarcoma by reprogramming nonsense-mediated RNA decay and SNAI2-mediated epithelial-to-mesenchymal transition"

### Network Stats

| | |
|---|---|
| number of nodes: | 555 |
| number of edges: | 64 |
| average node degree: | 0.231 |
| avg. local clustering coefficient: | 0.0907 |
| expected number of edges: | 24 |
| PPI enrichment p-value: | 1.15e-11 |

### Functional enrichments in network

#### Biological Process (Gene Ontology)

| *GO-term description* | *count in network* | *strength* | *false discovery rate* |
|---|---|---|---|
| GO:0051673 Membrane disruption in other organism | 7 of 12 | 1.31 | 0.00071 |
| GO:0021889 Olfactory bulb interneuron differentiation | 5 of 12 | 1.17 | 0.0197 |
| GO:0021772 Olfactory bulb development | 8 of 32 | 0.95 | 0.0050 |
| GO:0002227 Innate immune response in mucosa | 6 of 26 | 0.91 | 0.0431 |
| GO:1905606 Regulation of presynapse assembly | 7 of 34 | 0.86 | 0.0267 |
| GO:0051965 Positive regulation of synapse assembly | 13 of 66 | 0.84 | 0.00067 |
| GO:0021879 Forebrain neuron differentiation | 8 of 46 | 0.79 | 0.0256 |
| GO:0050832 Defense response to fungus | 8 of 48 | 0.77 | 0.0307 |
| GO:0021872 Forebrain generation of neurons | 9 of 58 | 0.74 | 0.0229 |
| GO:0019731 Antibacterial humoral response | | | |

| | | | | |
|---|---|---|---|---|
| 9 of 59 | 0.73 | 0.0248 | GO:0009620 | Response to fungus |
| 9 of 61 | 0.72 | 0.0284 | GO:0051963 | Regulation of synapse assembly |
| 14 of 106 | 0.67 | 0.0032 | GO:0070268 | Cornification |
| 14 of 113 | 0.64 | 0.0050 | GO:0007416 | Synapse assembly |
| 11 of 96 | 0.61 | 0.0381 | GO:0035821 | Modulation of process of other organism |
| 12 of 115 | 0.57 | 0.0431 | GO:0021953 | Central nervous system neuron differentiation |
| 18 of 183 | 0.54 | 0.0051 | GO:0007156 | Homophilic cell adhesion via plasma membrane adhesion molecules |
| 16 of 164 | 0.54 | 0.0120 | GO:0098742 | Cell-cell adhesion via plasma-membrane adhesion molecules |
| 24 of 257 | 0.52 | 0.0014 | GO:0031424 | Keratinization |
| 20 of 226 | 0.49 | 0.0064 | GO:0050807 | Regulation of synapse organization |
| 20 of 228 | 0.49 | 0.0068 | GO:0007608 | Sensory perception of smell |
| 33 of 411 | 0.45 | 0.00067 | GO:0030216 | Keratinocyte differentiation |
| 21 of 268 | 0.44 | 0.0159 | GO:0007187 | G protein-coupled receptor signaling pathway, coupled to cyclic nucleotide second messenger |

| | | | |
|---|---|---|---|
| GO:0099537 Trans-synaptic signaling | 19 of 254 | 0.42 | 0.0446 |
| GO:0007268 Chemical synaptic transmission | 32 of 436 | 0.41 | 0.0024 |
| GO:0098609 Cell-cell adhesion | 30 of 418 | 0.4 | 0.0041 |
| GO:0050906 Detection of stimulus involved in sensory perception | 35 of 505 | 0.39 | 0.0024 |
| GO:0007606 Sensory perception of chemical stimulus | 35 of 497 | 0.39 | 0.0024 |
| GO:0030900 Forebrain development | 34 of 484 | 0.39 | 0.0024 |
| GO:0061564 Axon development | 27 of 387 | 0.39 | 0.0115 |
| GO:0043588 Skin development | 28 of 421 | 0.37 | 0.0159 |
| GO:0030855 Epithelial cell differentiation | 25 of 382 | 0.36 | 0.0379 |
| GO:0050877 Nervous system process | 42 of 673 | 0.34 | 0.0025 |
| GO:0051606 Detection of stimulus | 80 of 1352 | 0.32 | 4.49e-06 |
| GO:0007600 Sensory perception | 39 of 659 | 0.32 | 0.0088 |
| GO:0007155 Cell adhesion | 53 of 923 | 0.31 | 0.0024 |
| | 53 of 925 | 0.31 | 0.0024 |

| | | | | |
|---|---|---|---|---|
| GO:0030182 | Neuron differentiation | 56 of 1019 | 0.29 | 0.0025 |
| GO:0003008 | System process | 104 of 1942 | 0.28 | 3.03e-06 |
| GO:0007186 | G protein-coupled receptor signaling pathway | 68 of 1255 | 0.28 | 0.00072 |
| GO:0009888 | Tissue development | 88 of 1760 | 0.25 | 0.00067 |
| GO:0060429 | Epithelium development | 56 of 1109 | 0.25 | 0.0115 |
| GO:0007417 | Central nervous system development | 50 of 988 | 0.25 | 0.0238 |
| GO:0009887 | Animal organ morphogenesis | 49 of 967 | 0.25 | 0.0256 |
| GO:0048699 | Generation of neurons | 72 of 1551 | 0.21 | 0.0120 |
| GO:0022008 | Neurogenesis | 74 of 1657 | 0.2 | 0.0248 |
| GO:0009653 | Anatomical structure morphogenesis | 93 of 2165 | 0.18 | 0.0137 |
| GO:0048869 | Cellular developmental process | 151 of 3757 | 0.15 | 0.0025 |
| GO:0030154 | Cell differentiation | 149 of 3702 | 0.15 | 0.0026 |
| GO:0032501 | Multicellular organismal process | 270 of 6933 | 0.14 | 2.79e-06 |

| | count in network | strength | false discovery rate |
|---|---|---|---|
| **GO:0048513**<br>**Animal organ development** | **126 of 3197** | **0.14** | **0.0212** |
| **GO:0048731**<br>**System development** | **171 of 4426** | **0.13** | **0.0038** |
| **GO:0048856**<br>**Anatomical structure development** | **200 of 5402** | **0.12** | **0.0051** |
| **GO:0007275**<br>**Multicellular organism development** | **188 of 5023** | **0.12** | **0.0055** |
| **GO:0032502**<br>**Developmental process** | **211 of 5841** | **0.11** | **0.0100** |

### KEGG Pathways

| *Pathway description* | *count in network* | *strength* | *false discovery rate* |
|---|---|---|---|
| hsa00982<br>Drug metabolism - cytochrome P450 | 9 of 64 | 0.7 | 0.0294 |
| hsa05204<br>Chemical carcinogenesis | 9 of 75 | 0.63 | 0.0426 |
| hsa04740<br>Olfactory transduction | 32 of 420 | 0.43 | 0.00041 |
| hsa04080<br>Neuroactive ligand-receptor interaction | 22 of 330 | 0.37 | 0.0376 |

### Reactome Pathways

| *Pathway description* | *count in network* | *strength* | *false discovery rate* |
|---|---|---|---|
| HSA-1462054<br>Alpha-defensins | | | |

| | | | |
|---|---|---|---|
| | 5 of 9 | 1.29 | 0.0069 |
| HSA-388844 | | | |
| Receptor-type tyrosine-protein phosphatases | | | |
| | 6 of 19 | 1.05 | 0.0117 |
| HSA-6794362 | | | |
| Protein-protein interactions at synapses | | | |
| | 12 of 85 | 0.7 | 0.0045 |
| HSA-6809371 | | | |
| Formation of the cornified envelope | | | |
| | 15 of 127 | 0.62 | 0.0040 |
| HSA-6805567 | | | |
| Keratinization | | | |
| | 21 of 210 | 0.55 | 0.0014 |
| HSA-381753 | | | |
| Olfactory Signaling Pathway | | | |
| | 32 of 391 | 0.46 | 0.00062 |
| HSA-418555 | | | |
| G alpha (s) signalling events | | | |
| | 38 of 532 | 0.4 | 0.00062 |
| HSA-388396 | | | |
| GPCR downstream signalling | | | |
| | 58 of 1094 | 0.27 | 0.0038 |
| HSA-372790 | | | |
| Signaling by GPCR | | | |
| | 60 of 1166 | 0.26 | **0.0041** |
