## Supplemental File 2 for "Vitamin D inhibits osteosarcoma by reprogramming nonsense-mediated RNA decay and SNAI2-mediated epithelial-to-mesenchymal transition"

**Network Stats**
number of nodes: 1102
number of edges: 367
average node degree: 0.666
avg. local clustering coefficient: 0.142
expected number of edges: 92
PPI enrichment p-value: < 1.0e-16

**Functional enrichments in network**

### Biological Process (Gene Ontology)

| *GO-term description* | *count in network* | *strength* | *false discovery rate* |
|---|---|---|---|
| GO:2001030 Negative regulation of cellular glucuronidation | 6 of 8 | 1.12 | 0.0097 |
| GO:1904224 Negative regulation of glucuronosyltransferase activity | 6 of 8 | 1.12 | 0.0097 |
| GO:0052696 Flavonoid glucuronidation | 5 of 7 | 1.1 | 0.0378 |
| GO:1902644 Tertiary alcohol metabolic process | 6 of 13 | 0.91 | 0.0471 |
| GO:0009812 Flavonoid metabolic process | 6 of 13 | 0.91 | 0.0471 |
| GO:0033141 Positive regulation of peptidyl-serine phosphorylation of stat protein | 9 of 21 | 0.88 | 0.0047 |
| GO:0002323 Natural killer cell activation involved in immune response | 10 of 24 | 0.87 | 0.0023 |
| GO:0042573 Retinoic acid metabolic process | 11 of 32 | 0.79 | 0.0033 |

GO:0002548
Monocyte chemotaxis

14 of 43 0.76 0.00051
GO:0001580
Detection of chemical stimulus involved in sensory perception of bitter taste

12 of 39 0.74 0.0033
GO:0086010
Membrane depolarization during action potential

9 of 30 0.73 0.0279
GO:0050913
Sensory perception of bitter taste

13 of 44 0.72 0.0023
GO:0042100
B cell proliferation

11 of 43 0.66 0.0200
GO:0031424
Keratinization

55 of 226 0.64 1.99e-13
GO:0043330
Response to exogenous dsrna

11 of 46 0.63 0.0294
GO:0048247
Lymphocyte chemotaxis

11 of 50 0.59 0.0471
GO:0030101
Natural killer cell activation

12 of 56 0.58 0.0351
GO:0030216
Keratinocyte differentiation

56 of 268 0.57 1.57e-11
GO:0034308
Primary alcohol metabolic process

17 of 82 0.57 0.0047
GO:0070252
Actin-mediated cell contraction

18 of 90 0.55 0.0042

GO:0031640
Killing of cells of other organism

18 of 91 0.55 0.0047

GO:0050829
Defense response to gram-negative bacterium

19 of 98 0.54 0.0037

GO:0034754
Cellular hormone metabolic process

24 of 125 0.53 0.00051

GO:0009913
Epidermal cell differentiation

59 of 315 0.52 1.28e-10

GO:0061844
Antimicrobial humoral immune response mediated by antimicrobial peptide

21 of 113 0.52 0.0026

GO:0042180
Cellular ketone metabolic process

14 of 76 0.51 0.0420

GO:0120254
Olefinic compound metabolic process

19 of 106 0.5 0.0077

GO:1990869
Cellular response to chemokine

16 of 90 0.5 0.0265

GO:0001508
Action potential

17 of 97 0.49 0.0202

GO:0043588
Skin development

65 of 382 0.48 2.51e-10

GO:0019730
Antimicrobial humoral response

27 of 160 0.48 0.00084

| | | | | |
|---|---|---|---|---|
| GO:0008544 | Epidermis development | 69 of 419 | 0.47 | 2.02e-10 |
| GO:0006959 | Humoral immune response | 46 of 275 | 0.47 | 1.11e-06 |
| GO:0001906 | Cell killing | 21 of 126 | 0.47 | 0.0077 |
| GO:0006805 | Xenobiotic metabolic process | 18 of 112 | 0.46 | 0.0301 |
| GO:0042445 | Hormone metabolic process | 31 of 194 | 0.45 | 0.00051 |
| GO:0030048 | Actin filament-based movement | 19 of 121 | 0.45 | 0.0278 |
| GO:0097529 | Myeloid leukocyte migration | 19 of 123 | 0.44 | 0.0315 |
| GO:0035637 | Multicellular organismal signaling | 20 of 137 | 0.41 | 0.0403 |
| GO:0060326 | Cell chemotaxis | 29 of 204 | 0.4 | 0.0049 |
| GO:0030855 | Epithelial cell differentiation | 92 of 673 | 0.39 | 2.51e-10 |
| GO:0050907 | Detection of chemical stimulus involved in sensory perception | 60 of 434 | 0.39 | 2.36e-06 |
| GO:0009593 | Detection of chemical stimulus | | | |

63 of 470 0.38 2.55e-06

GO:0070374

Positive regulation of erk1 and erk2 cascade

28 of 209 0.38 0.0132

GO:0006813

Potassium ion transport

23 of 174 0.37 0.0475

GO:0007606

Sensory perception of chemical stimulus

63 of 484 0.36 4.80e-06

GO:0050906

Detection of stimulus involved in sensory perception

64 of 497 0.36 5.01e-06

GO:0042742

Defense response to bacterium

36 of 277 0.36 0.0035

GO:0007608

Sensory perception of smell

50 of 411 0.33 0.00057

GO:0007187

G protein-coupled receptor signaling pathway, coupled to cyclic nucleotide second messenger

30 of 254 0.32 0.0424

GO:0007186

G protein-coupled receptor signaling pathway

142 of 1255 0.3 9.60e-11

GO:0051606

Detection of stimulus

74 of 659 0.3 4.18e-05

GO:0070372

Regulation of erk1 and erk2 cascade

33 of 292 0.3 0.0433

GO:0050877

Nervous system process

145 of 1352 0.28 1.12e-09

| | | | | |
|---|---|---|---|---|
| GO:0007600 | Sensory perception | 100 of 923 | 0.28 | 2.55e-06 |
| GO:0006935 | Chemotaxis | 57 of 545 | 0.27 | 0.0049 |
| GO:0010817 | Regulation of hormone levels | 55 of 524 | 0.27 | 0.0057 |
| GO:0003008 | System process | 199 of 1942 | 0.26 | 5.75e-12 |
| GO:0060429 | Epithelium development | 114 of 1109 | 0.26 | 2.55e-06 |
| GO:0042391 | Regulation of membrane potential | 45 of 440 | 0.26 | 0.0351 |
| GO:0098542 | Defense response to other organism | 88 of 900 | 0.24 | 0.00055 |
| GO:0006952 | Defense response | 123 of 1296 | 0.23 | 2.45e-05 |
| GO:0009888 | Tissue development | 155 of 1760 | 0.19 | 3.09e-05 |
| GO:0051707 | Response to other organism | 108 of 1256 | 0.18 | 0.0051 |
| GO:0007267 | Cell-cell signaling | 98 of 1145 | 0.18 | 0.0117 |
| GO:0034220 | Ion transmembrane transport | | | |

| | | | |
|---|---|---|---|
| GO:0006811 Ion transport | 87 of 1010 | 0.18 | 0.0230 |
| GO:0032501 Multicellular organismal process | 109 of 1344 | 0.16 | 0.0279 |
| GO:0048513 Animal organ development | 553 of 6933 | 0.15 | 2.01e-18 |
| GO:0030154 Cell differentiation | 254 of 3197 | 0.15 | 8.70e-06 |
| GO:0048869 Cellular developmental process | 290 of 3702 | 0.14 | 2.55e-06 |
| GO:0006928 Movement of cell or subcellular component | 291 of 3757 | 0.14 | 5.67e-06 |
| GO:0009605 Response to external stimulus | 118 of 1501 | 0.14 | 0.0424 |
| GO:0007275 Multicellular organism development | 174 of 2310 | 0.13 | 0.0150 |
| GO:0048731 System development | 373 of 5023 | 0.12 | 2.55e-06 |
| GO:0048856 Anatomical structure development | 328 of 4426 | 0.12 | 2.78e-05 |
| GO:0023052 Signaling | 396 of 5402 | 0.11 | 2.55e-06 |
| | 381 of 5239 | 0.11 | 1.06e-05 |

| | | | |
|---|---|---|---|
| GO:0007154 Cell communication | 383 of 5320 | 0.11 | 2.60e-05 |
| GO:0032502 Developmental process | 417 of 5841 | 0.1 | 1.15e-05 |
| GO:0007165 Signal transduction | 343 of 4876 | 0.1 | 0.0013 |
| GO:0042221 Response to chemical | 308 of 4333 | 0.1 | 0.0022 |
| GO:0050896 Response to stimulus | 522 of 8046 | 0.06 | 0.0059 |

**KEGG Pathways**

| *Pathway description* | *count in network* | *strength* | *false discovery rate* |
|---|---|---|---|
| **hsa00140** **Steroid hormone biosynthesis** | **20 of 59** | **0.78** | **1.28e-06** |
| **hsa00053** **Ascorbate and aldarate metabolism** | **7 of 26** | **0.68** | **0.0424** |
| **hsa00830** **Retinol metabolism** | **16 of 64** | **0.65** | **0.00028** |
| **hsa04742** **Taste transduction** | **19 of 81** | **0.62** | **0.00020** |
| **hsa00980** **Metabolism of xenobiotics by cytochrome P450** | **16 of 69** | **0.61** | **0.00045** |
| **hsa05204** **Chemical carcinogenesis** | **17 of 75** | **0.6** | **0.00036** |

| Pathway | Description | count in network | strength | false discovery rate |
|---|---|---|---|---|
| hsa00590 | Arachidonic acid metabolism | 13 of 61 | 0.58 | 0.0055 |
| hsa00982 | Drug metabolism - cytochrome P450 | 13 of 64 | 0.56 | 0.0061 |
| hsa04976 | Bile secretion | 16 of 89 | 0.5 | 0.0055 |
| hsa04623 | Cytosolic DNA-sensing pathway | 11 of 62 | 0.5 | 0.0424 |
| hsa04060 | Cytokine-cytokine receptor interaction | 38 of 282 | 0.38 | 0.00028 |
| hsa04740 | Olfactory transduction | 51 of 420 | 0.33 | 0.00020 |
| hsa04080 | Neuroactive ligand-receptor interaction | 37 of 330 | 0.3 | 0.0055 |

Reactome Pathways

| *Pathway* | *description* | *count in network* | *strength* | *false discovery rate* |
|---|---|---|---|---|
| HSA-211935 | Fatty acids | 6 of 15 | 0.85 | 0.0495 |
| HSA-156588 | Glucuronidation | 8 of 23 | 0.79 | 0.0182 |
| HSA-445095 | Interaction between L1 and Ankyrins | 10 of 31 | 0.76 | 0.0064 |
| HSA-912694 | Regulation of IFNA signaling | 8 of 25 | 0.75 | 0.0242 |
| HSA-5576892 | Phase 0 - rapid depolarisation | 10 of 32 | 0.74 | 0.0075 |
| HSA-1461957 | Beta defensins | | | |

| | | | | |
|---|---|---|---|---|
| HSA-933541 | TRAF6 mediated IRF7 activation | 10 of 34 | 0.72 | 0.0101 |
| HSA-6805567 | Keratinization | 8 of 28 | 0.71 | 0.0440 |
| HSA-1461973 | Defensins | 54 of 210 | 0.66 | 1.71e-14 |
| HSA-420499 | Class C/3 (Metabotropic glutamate/pheromone receptors) | 11 of 43 | 0.66 | 0.0124 |
| HSA-6809371 | Formation of the cornified envelope | 10 of 39 | 0.66 | 0.0216 |
| HSA-6803157 | Antimicrobial peptides | 30 of 127 | 0.62 | 5.60e-07 |
| HSA-211897 | Cytochrome P450 - arranged by substrate type | 20 of 87 | 0.61 | 0.00020 |
| HSA-211945 | Phase I - Functionalization of compounds | 13 of 65 | 0.55 | 0.0216 |
| HSA-1296071 | Potassium Channels | 19 of 105 | 0.51 | 0.0048 |
| HSA-211859 | Biological oxidations | 18 of 103 | 0.49 | 0.0094 |
| HSA-375276 | Peptide ligand-binding receptors | 32 of 214 | 0.42 | 0.00063 |
| HSA-381753 | Olfactory Signaling Pathway | 28 of 193 | 0.41 | 0.0038 |
| HSA-500792 | GPCR ligand binding | 50 of 391 | 0.36 | 0.00011 |

| | | | |
|---|---|---|---|
| | 56 of 455 | 0.34 | 8.31e-05 |
| HSA-418555 G alpha (s) signalling events | 64 of 532 | 0.33 | 3.36e-05 |
| HSA-373076 Class A/1 (Rhodopsin-like receptors) | 39 of 323 | 0.33 | 0.0041 |
| HSA-112316 Neuronal System | 46 of 406 | 0.3 | 0.0038 |
| HSA-372790 Signaling by GPCR | 125 of 1166 | 0.28 | 4.45e-08 |
| HSA-388396 GPCR downstream signalling | 115 of 1094 | 0.27 | 5.60e-07 |
| HSA-418594 G alpha (i) signalling events | 42 of 396 | 0.27 | 0.0184 |
| HSA-1266738 Developmental Biology | 105 of 1087 | 0.23 | 8.31e-05 |
| HSA-162582 Signal Transduction | 195 of 2741 | 0.1 | 0.0440 |
